# Origin of the nuclear proteome on the basis of pre-existing nuclear localization signals in prokaryotic proteins

**DOI:** 10.1101/2020.01.07.896449

**Authors:** Olga M. Lisitsyna, Margarita A. Kurnaeva, Eugene A. Arifulin, Maria Y. Shubina, Yana R. Musinova, Andrey A. Mironov, Eugene V. Sheval

## Abstract

The origin of the selective nuclear protein import machinery, which consists of nuclear pore complexes and adaptor molecules interacting with the nuclear localization signals (NLSs) of cargo molecules, was one of the most important events in the evolution of the eukaryotic cell. How the proteins were selected for import into the forming nuclei remains an open question. Here, we demonstrate that functional NLSs may be integrated inside nucleotide-binding domains of both eukaryotic and prokaryotic proteins and may co-evolve with these domains. We propose that the pre-existence of NLSs inside prokaryotic proteins dictated, at least partially, the nuclear proteome composition.

## 1. Introduction

The acquisition of the cell nucleus enabled spatial segregation of transcription and translation and likely permitted the evolution of more sophisticated mechanisms of gene expression regulation [1]. Since proteins are translated in the cytoplasm, the emergence of a reliable and efficient nuclear import mechanism was the essential event leading to the origin of the eukaryotic cell. Nucleocytoplasmic transport across the nuclear envelope occurs predominantly through nuclear pore complexes (NPCs). Small proteins can freely diffuse through NPCs, but globular molecules larger than ~40 kDa are selectively transferred by an energy-dependent mechanism that requires additional transport factors, called karyopherins, which recognize nuclear localization signals (NLSs) in their cargo proteins [2]. Past studies have revealed some important events in the evolution of the nuclear envelope and the possible ancestors of the key elements of import machinery: NPCs and karyopherins [3–7]. However, it remains unclear how the proteins were selected for import into the forming nuclei, i.e., how the nuclear proteome evolved.

## 2. Materials and Methods

Human proteins containing NLSs were collected from NLSdb (https://rostlab.org/services/nlsdb/) and the UniProt database. The annotations of protein domain structure were obtained from the Uniprot/Swiss-Prot database. The regions between nearest annotated domains were analysed here as out-of-domain regions. The orthologues of human proteins with NLSs were found in the *B. floridae*, *D. rerio*, *X. laevis*, *P. sinensis*, and *G. gallus* proteomes by using OrthoDB release 10 (https://www.orthodb.org/) (Supplementary Table S1).

Multiple alignment of orthologous sequences was performed with Clustal Omega. The conservation degree of multiple alignments was evaluated as the *IC* [8], which was calculated as follows:

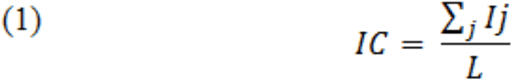

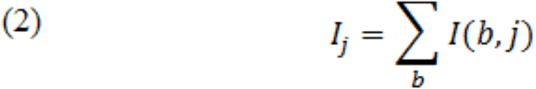

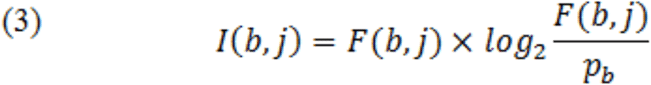

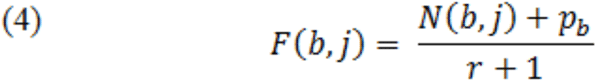

where *I*_*j*_ is the IC of the *j*^*th*^ alignment column, *L* – length of multiple alignment, *I*_*(b,j)*_ is the *IC* of amino acid residues in the *j*^*th*^ alignment column, *F*_*(b, j)*_ is the frequency of amino acid residues in the *j*^*th*^ alignment column, *N*_*(b, j)*_ is the number of one type of amino acid residue taking into account the base frequency of amino acids *(p_b_)* as the pseudo count [9], and *r* is the number of rows in the alignment.

The *Thermococcus sibiricus* lineage was kindly provided by E.A. Bonch-Osmolovskaya. Genomic DNA of *Synechococcus* sp. and *Anabaena* sp. was provided by O.A. Koksharova, and that of *Vibrio harveyi*, by Y.V. Bertsova and A.V. Bogachev. Genes of target prokaryotic proteins were amplified by PCR from corresponding genomic DNA and inserted into the pEGFP-C1 vector (Clontech). Mutated genes of prokaryotic proteins were obtained by PCR site-specific mutagenesis. Double-stranded oligonucleotides encoding predicted NLSs of prokaryotic proteins were inserted into the pEGFP-C1 vector. DNA fragments encoding M9M and Bimax2 peptides were inserted into the pTagRFP-C vector (Evrogen).

HeLa cells were grown in Dulbecco’s modified Eagle’s medium supplemented with L-glutamine, 10% foetal calf serum (HyClone) and antibiotic/antimycotic solution (Gibco). Cellular transfection was performed using Lipofectamine 2000 reagent (Thermo Fisher Scientific) according to the manufacturer’s instructions. Images of at least 20 living HeLa cells expressing EGFP-fused proteins were acquired in two different experiments using a Nikon C2 confocal laser scanning microscope. The ratio of nucleoplasmic EGFP concentration to cytoplasmic EGFP concentration (F_nuc_/F_cyt_) was measured as described elsewhere [10]. Statistical analysis and graph preparations were performed using Prism 6 (GraphPad software).

## 3. Results

To address this question, we analysed data on NLSs and their localization relative to protein domains. We collected a dataset consisting of 592 annotated NLSs from 496 human proteins, among which 234 NLSs were identified experimentally and the other 358 NLS sequences were annotated *in silico* (Supplementary Table S1). Forty-five percent of all NLSs overlapped with some annotated domains (19% with nucleotide-binding domains and 26% with domains involved in protein-protein interactions), while the other 55% of NLSs exhibited out-of-domain localization (Fig. 1a). The majority (77%) of the nucleotide-binding domains matched with NLSs were annotated as DNA-binding domains (Fig. 1a). Our data are in agreement with the published data about co-localization of NLS with DNA- and RNA-binding domains [11, 12].

**Figure 1.**
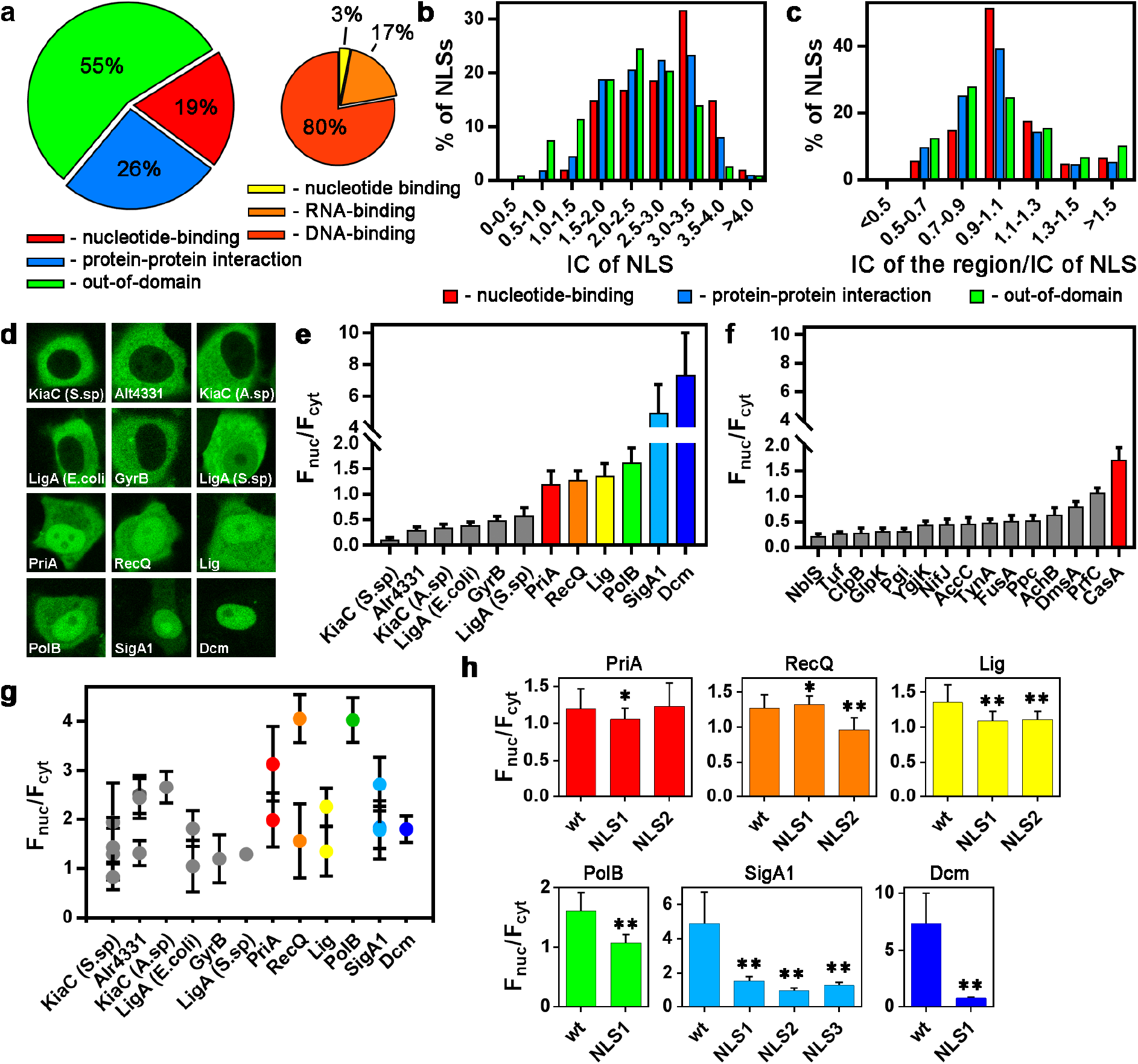
Evolutionary integration of NLSs inside annotated domains of eukaryotic and prokaryotic proteins. *(a)* Distribution of NLSs according to their localization in protein sequences relative to annotated protein domains. *(b)* IC distributions of NLSs that overlapped with either nucleotide-binding domains, domains involved in protein-protein interactions or out-of-domain regions. The shift of distributions of NLS overlapping with nucleotide-binding domains and domains involved in protein-protein interactions towards higher values of IC suggest that in-domain NLSs are more conservative relative to out-of-domain regions (one-way ANOVA test, p < 0.05, followed by the Bonferroni correction for multiple comparisons) *(c)*. Distribution of the ratio of IC of the surrounding NLS region to that of the NLS. *(d)* Localization of prokaryotic proteins that were expressed as EGFP fusions in living HeLa cells. *(e)* Estimation of nuclear accumulation (F_nuc_/F_cyt_) of prokaryotic proteins fused with EGFP. The results are presented as the mean±s.d. (n>20). Proteins with an F_nuc_/F_cyt_ ≤ 1.16 were classified as non-accumulated inside nuclei (grey bars), whereas those with an F_nuc_/F_cyt_ > 1.16 were classified as accumulated inside nuclei (coloured bars). *(f)* Estimation of nuclear accumulation of different prokaryotic proteins for which the presence of NLS(s) was not predicted using cNLS Mapper software (mean±s.d.) (n>20). *(g)* Estimation of nuclear accumulation of EGFP fused with the predicted NLSs from different prokaryotic proteins (mean±s.d.) (n>20). *(h)* Mutations in predicted NLSs influence the nuclear accumulation (F_nuc_/F_cyt_) of prokaryotic proteins. Each value represents the mean±s.d. (n>20). single asterisk: p<0.05, double asterisk: p<0.0001, Mann-Whitney test.

We hypothesized that there was an evolutionary link between NLSs and domains. To test this hypothesis, the conservation of NLSs and surrounding regions were analysed by comparing the human protein sequences with their orthologues from five different species of phylum Chordata (*Branchiostoma floridae*, *Danio rerio*, *Xenopus laevis*, *Pelodiscus sinensis* and *Gallus gallus*). The degree of conservation of NLSs and surrounding regions (domains or out-of-domain regions) was evaluated as the information content (IC) of the obtained multiple alignment when the most conserved position in the alignment had a higher value of information content. Comparison of the calculated information content distribution in three groups of NLSs demonstrated that NLSs overlapping the annotated domains (both nucleotide-binding domains and domains involved in protein-protein interactions) were more conservative than NLSs located outside annotated domains (one-way ANOVA test, p < 0.05, followed by the Bonferroni correction for multiple comparisons) (Fig. 1b). To compare conservation between protein regions and the NLSs overlapping these regions, the ratio of IC of each region to IC of the corresponding NLS was calculated (Fig. 1c). Approximately half of all NLSs overlapping nucleotide-binding domains had the same conservation degree as the corresponding domains (for 51% of NLSs, the ratio was within the 0.9-1.1 interval). These NLSs were integrated into nucleotide-binding domains, and their evolution might depend on the evolution of the domains. The NLSs overlapping domains involved in protein-protein interactions demonstrated lower similarity with the surrounding protein regions (for 39% of NLSs, the ratio was within the 0.9-1.1 interval), and the NLSs located outside domains did not demonstrate substantial similarity with the surrounding regions (the ratio was within the 0.9-1.1 interval only for 25% of NLSs).

NLSs are short and structurally simple sequences. For example, a monopartite ‘classical’ NLS has a degenerate consensus sequence of K(K/R)X(K/R) [13]. Since nucleotide-binding domains are enriched in positively charged amino acids, the occasional appearance of such NLSs inside such domains seems probable. Since similar nucleotide-binding domains may be found inside prokaryotic proteins, it seems plausible that these domains already contain sequences that can potentially play a role in NLSs. If this supposition is correct, then after expression inside eukaryotic cells, such prokaryotic proteins would accumulate inside cell nuclei. We cloned 12 large (>45 kDa) prokaryotic proteins with nucleotide-binding domains. In all these proteins, the NLSs were predicted using cNLS Mapper [14], and at least one of these NLSs overlapped with nucleotide-binding domains (Supplementary Table S2). To produce a control group of proteins, we cloned and analysed 15 large (>45 kDa) proteins without predicted NLSs (Supplementary Table S2). The proteins were fused with enhanced green fluorescent protein (EGFP), and their localization was investigated in living HeLa cells. Approximately half of all the proteins accumulated inside the nuclei, although to different degrees (Fig. 1d). To quantify the efficiency of nuclear accumulation in the nucleus, the ratio of nucleoplasmic to cytoplasmic (F_nuc_/F_cyt_) fluorescence was measured for all the proteins as described elsewhere [10]. Proteins with F_nuc_/F_cyt_ values higher than that of EGFP, i.e., > 1.16, were classified as being accumulated inside nuclei (Fig. 1e, Supplementary Table S2). No correlation between the efficiency of nuclear accumulation and the molecular weight of prokaryotic proteins was detected (Pearson correlation coefficient = 0.13), indicating that the transfer of proteins into the nuclei was not due to diffusion but rather due to an active process. Among these proteins, only one was accumulated inside nuclei (F_nuc_/F_cyt_ = 1.71±0.25), and this protein was the only one with DNA-binding activity among the 15 control proteins (Fig. 1f; Supplementary Table S2). These data are in agreement with published results indicating that NLSs are present not only in the proteins of eukaryotes but also in the proteins of prokaryotes [15–22] and bacteriophages [23].

Nuclear import of large (>40 kDa) proteins depends on the presence of NLSs. NLSs were predicted in all investigated prokaryotic proteins using cNLS Mapper [13] (Supplementary Table S2). To confirm that these protein regions are indeed functionally active NLSs, we constructed plasmids coding predicted NLSs fused with EGFP. All predicted NLSs were able to accumulate EGFP inside nuclei (Fig. 1g, Supplementary Table S3), but the F_nuc_/F_cyt_ values of the predicted NLSs did not correlate with the F_nuc_/F_cyt_ values of the corresponding full-length proteins (Supplementary Fig. S1) (Pearson’s correlation coefficient between the F_nuc_/F_cyt_ of the strongest among all predicted NLSs and the F_nuc_/F_cyt_ of the proteins = 0.14). Therefore, these results can be considered only an indication of the potential NLS activity. Therefore, we also used site-specific mutagenesis to directly detect the presence of NLSs.

Substitutions of all positively charged amino acids inside each predicted NLS with alanine decreased the nuclear accumulation (F_nuc_/F_cyt_) of all proteins that had been classified as accumulated inside nuclei (F_nuc_/F_cyt_ > 1.16) (Fig. 1h, Supplementary Table S4).

Nuclear import of proteins containing a classical NLS depends on the interaction of the NLS sequence with karyopherin-*α* and karyopherin-*β*. The non-classical NLSs directly interact with karyopherin-*β* for import to the nuclei. We used the inhibitors Bimax2 [24] and M9M [25], which are highly specifically bound to karyopherin-*α* and karyopherin-*β2*, respectively. Nuclear accumulation of PriA, Lig, PolB and SigA1 was decreased by both co-expressed Bimax2 (Supplementary Fig. S2) and M9M (Supplementary Fig. S3). The nuclear accumulation of Dcm was decreased by only Bimax2. Thus, these proteins accumulated inside nuclei via the ‘classical’ karyopherin-*α*/*β*-dependent pathway. Additionally, nuclear accumulation of RecQ was decreased by co-expression of M9M but not co-expression of Bimax2, indicating the presence of a non-classical NLS in this protein.

## 4. Discussion

Overall, our data indicate that regions enriched with positively charged amino acids of nucleotide-binding domains can indeed serve as genuine NLSs. These NLSs are integrated into domains, and their evolution might depend on the evolution of the domains. Such NLSs, even if they are integrated inside prokaryotic proteins, can interact with karyopherins. Karyopherins have many functions in the cell and, in particular, can act as chaperones [26, 27]. The protein domains interacting with karyopherins might have evolved before the origin of the nuclear envelope, and these domains contained sequences that can potentially play a role in NLSs. The pre-existence of NLSs interacting with karyopherins might have dictated, at least in part, which proteins were accumulated and compartmentalized inside nuclei, i.e., the content of proteins for forming the nuclear proteome. Proteins that did not have such integrated NLSs could have acquired them *de novo* after nuclear envelope formation, and these NLSs can be considered as separate units of genome evolution.

## Acknowledgments

We are grateful to E.A. Bonch-Osmolovskaya for providing *Thermococcus sibiricus* lineage, O.A. Koksharova for providing genomic DNA of *Anabaena* sp. and *Synechoccus* sp., Y.V. Bertsova and A.V. Bogachev for providing genomic DNA of *Vibrio harveyi*. We thank Y.S. Vassetzky for valuable discussion.

