## Supplementary Figures for "Origin of the nuclear proteome on the basis of pre-existing nuclear localization signals in prokaryotic proteins"

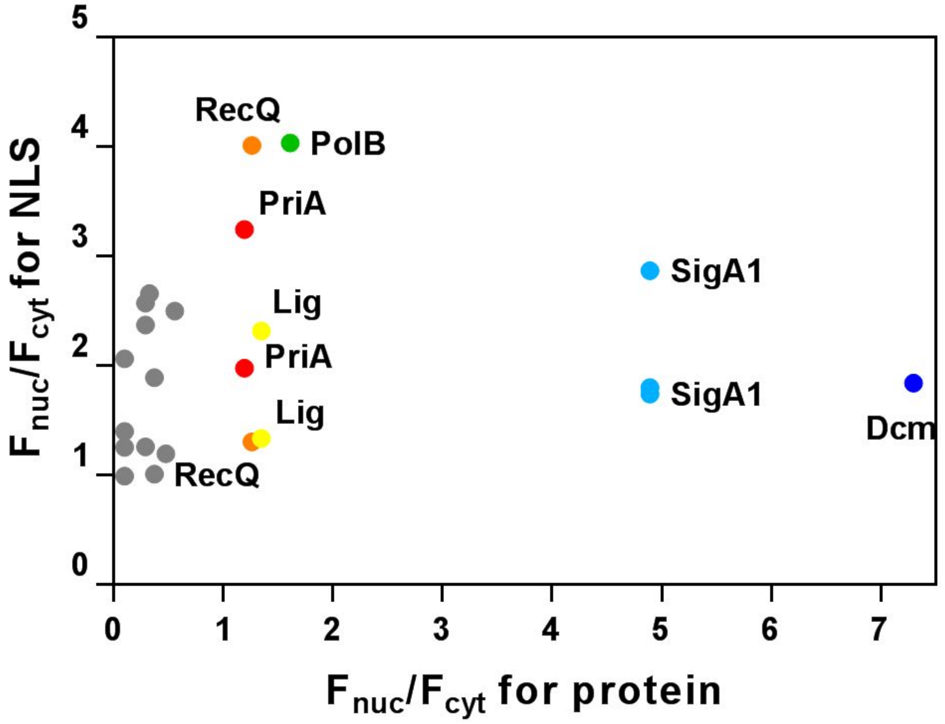


**Supplementary Fig. S1.** Comparison of nuclear accumulation ($F_{nuc}/F_{cyt}$) of all predicted NLSs and nuclear accumulation ($F_{nuc}/F_{cyt}$) of the full-length proteins fused with EGFP in living HeLa cells.

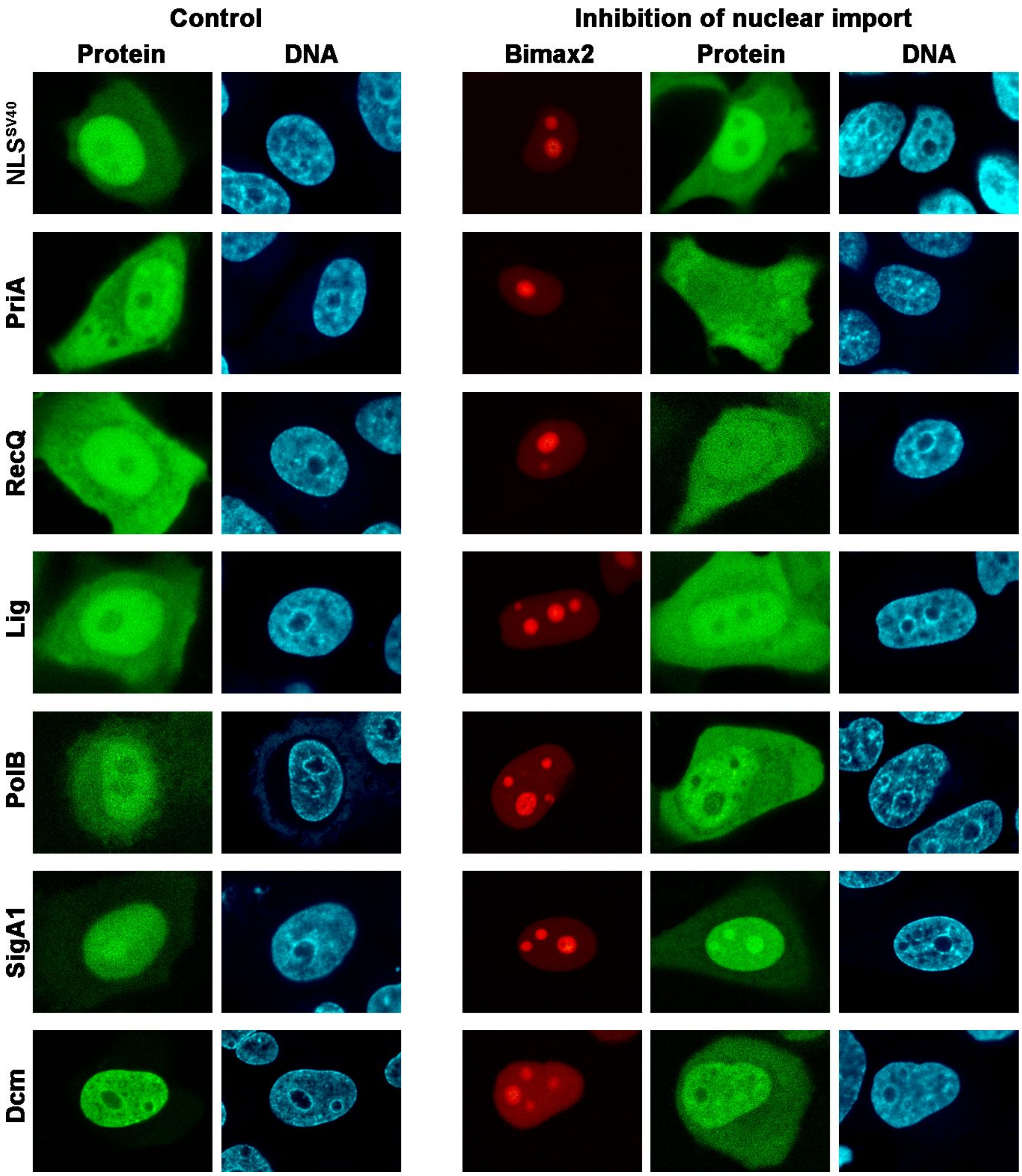


**Supplementary Fig. S2.** Decrease in nuclear accumulation of prokaryotic proteins by a peptide inhibitor of karyopherin-*α* (Bimax2). NLS from the T antigen of SV40 virus ($NLS^{SV40}$) fused with EGFP was used as a positive control. Expression of TagRFP-Bimax2 leads to a decrease in the nuclear accumulation of $NLS^{SV40}$. A decrease in nuclear accumulation was also detected for five prokaryotic proteins, namely, PriA, Lig, PolB, SigA1 and Dcm.

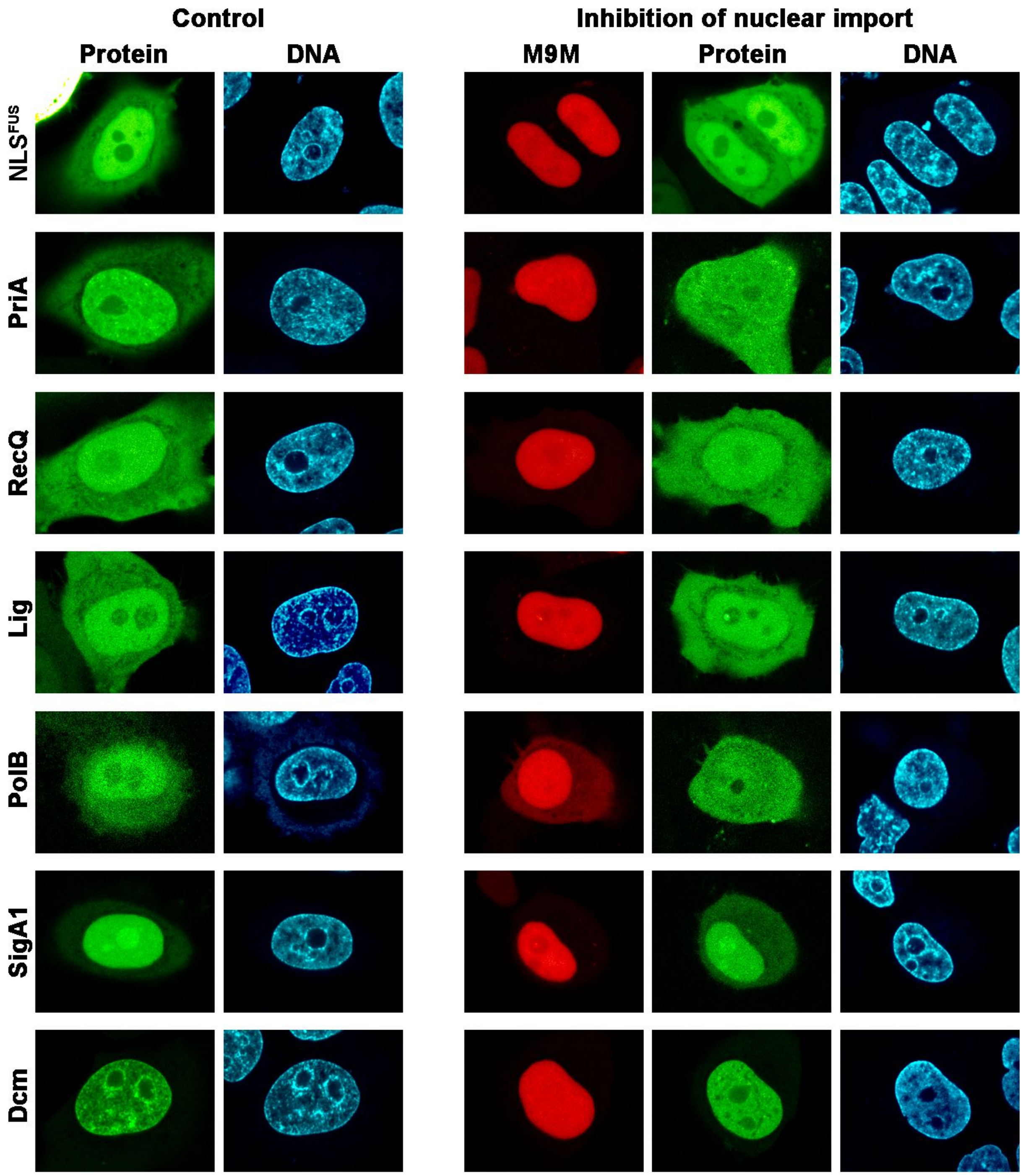


**Supplementary Fig. S3.** Decrease in the nuclear accumulation of prokaryotic proteins by a peptide inhibitor of karyopherin-*β2* (M9M). The NLS from FUS protein (NLS$^{FUS}$) fused with EGFP was used as a positive control. The expression of TagRFP-M9M leads to a decrease in the nuclear accumulation of NLS$^{FUS}$. A decrease in nuclear accumulation was also detected for five prokaryotic proteins, namely, PriA, RecQ, Lig, PolB and SigA1.
