## Supplementary Table S1 for "Origin of the nuclear proteome on the basis of pre-existing nuclear localization signals in prokaryotic proteins"

**Supplementary Table S1.** NLSs identified experimentally or predicted *in silico*.

**NLSs identified experimentally**

| № of NLS | NLS sequence | № of protein | Uniprot ID | Protein name | Protein function | Ortholog from *Danio rerio* (Uniprot/Uniprac ID) | Ortholog from *Branchiostoma floridae* (Uniprot/Uniprac ID) | Ortholog from *Xenopus leavis* (Uniprot/Uniprac ID) | Ortholog from *Pelodiscus sinesis* (Uniprot/Uniprac ID) | Ortholog from *Gallus gallus* (Uniprot/Uniprac ID) |
|---|---|---|---|---|---|---|---|---|---|---|
| 1 | KKRK LE | 1 | P02545 | Prelamin-A/C | Nuclear assembly, chromatin organization, nuclear membrane and telomere dynamics | Q90XD7 | C3ZXL7 | P11048 | K7G3U5 | P13648 |
| 2 | RRKLKN RV | 2 | P17861 | X-box-binding protein 1 | Transcription factor, regulates the unfolded protein response. | Q90X27 | C3XVS7 | Q7ZYC2 | - | F1NAC1 |
| 3 | RKKR | 3 | Q9BZS1 | Forkhead box protein P3 | Crucial for the development and inhibitory function of regulatory T-cells | A8WFW7 | C3XQK2 | Q5W1J5 | K7FP67 | Q58NQ4 |
| 4 | KSGKAPRR | 4 | O43524 | Forkhead box protein O3 | Transcriptional activator which triggers apoptosis in the absence of survival factors, including neuronal cell death upon oxidative stress | Q5I0F7 | C3YI59 | Q6EUW1 | K7F1W1 | F1NNE8 |
| 5 | QSSNFGPMKG GNFGGRSSGP YGGGGQYFAK PRNQGGY | 5 | P09651 | Heterogeneous nuclear ribonucleoprotein A1 | Involved in the packaging of pre-mRNA into hnRNP particles, transport of poly(A) mRNA from the nucleus to the cytoplasm and may modulate splice site selection | Q803K3 | - | Q7ZX33 | - | A0A1D5P5G1 |
| 6 | KKRK | 6 | Q13620 | Cullin-4B | Core component of multiple cullin-RING-based E3 ubiquitin-protein ligase complexes which mediate the ubiquitination and subsequent proteasomal degradation of target proteins | F8W428 | C3ZF52 | Q498E8 | K7GFE2 | E1BQK9 |
| 7 | KRK | 7 | Q15911 | Zinc finger homeobox protein 3 | Transcriptional regulator which can act as an activator or a repressor | A0A0R4ISP1 | C3ZBH5 | A0A1L8GKV0 | - | F1N9R6 |
| 8 | NYK RPMDGTYGPP AKRHEGE | 8 | O14497 | AT-rich interactive domain-containing protein 1A | Involved in transcriptional activation and repression of select genes by chromatin remodeling (alteration of DNA-nucleosome topology) | A0A0R4IVD0 | C3ZM58 | - | K7GH06 | A0A1D5NZ44 |
| 9 | KRK | 9 | O95644 | Nuclear factor of activated T-cells, cytoplasmic 1 | Plays a role in the inducible expression of cytokine genes in T-cells, especially in the induction of the IL-2 [illegible] | Q19AW4 | - | A0A1L8FSH1 | - | A0A1D5PI24 |
| 10 | KRKR | | | | | | | | | |
| 11 | KVPWLKPGRSPLPSHARSQPG LCNMYK | 10 | P02686 | Myelin basic protein | The most abundant protein component of the myelin membrane in the CNS. Protein has a role in both its formation and stabilization | E7EYA2 | - | A0A1L8FY91 | - | A0A1D5NWU0 |
| 12 | RRVPQRKEVS RCRKCRK | 11 | Q9NUL5 | Repressor of yield of DENV protein | Exhibits antiviral activity against dengue virus (DENV) and can inhibit the replication of all DENV serotypes | A7YY07 | - | A0A1L8H2R0 | - | - |
| 13 | RKKSIPLSI KNLKRKHKRK KNKITR | 12 | Q9C0C9 | E3-independent) E2 ubiquitin-conjugating enzyme | E2/E3 hybrid ubiquitin-protein ligase that displays both E2 and E3 ligase activities and mediates monoubiquitination of target proteins | A0A0G2KHN8 | C3Z6Y3 | A0A1L8EMX4 | - | F1P5I5 |
| 14 | KKQKKQHEG SKKK | 13 | P49591 | Serine-tRNA ligase, cytoplasmic | Catalyzes the attachment of serine to tRNA(Ser) in a two-step reaction: serine is first activated by ATP to form Ser-AMP and then transferred to the acceptor end of tRNA(Ser) | Q6DRC0 | C3YPI3 | Q7ZY49 | K7FUL9 | Q5ZM75 |
| 15 | RKRSPKDK KEKDLDGAGK RRKT | 14 | Q7RTP6 | [F-actin]-monooxygenase MICAL3 | Monooxygenase that promotes depolymerization of F-actin by mediating oxidation of specific methionine residues on actin to form methionine-sulfoxide, resulting in actin filament disassembly and preventing repolymerization | E7F9T0 | C3Z6G9 | - | - | Q5F3F3 |
| 16 | PKVRRRI | 15 | Q3KP66 | Innate immunity activator protein | Expressed in peripheral macrophages and intestinal myeloid-derived cells, is required for optimal PRR (pattern recognition receptor)-induced signaling, [illegible] | F1QAI0 | C3Z2X7 | - | K7FHM9 | A0A1L1RYD9 |
| 17 | PYEKPRK | | | | | | | | | |
| 18 | RRRPTH | | | | | | | | | |
| 19 | KRAR | 16 | O15381 | Nuclear valosin-containing protein-like | Participates in the assembly of the telomerase holoenzyme and effecting of telomerase activity via its [illegible] | Q803I9 | C3XWH4 | Q7ZXI4 | K7FKS9 | A0A1L1RMM8 |
| 20 | RK GKLKNKGSKR KK | | | | | | | | | |
| 21 | KRANS NLVAAYEKAK KK | 17 | P23508 | Colorectal mutant cancer protein | Suppresses cell proliferation and the Wnt/b-catenin pathway in colorectal cancer cells. Inhibits DNA binding of b-catenin/TCF/LEF transcription factors. Involved in cell migration independently of RAC1, CDC42 and p21-activated kinase (PAK) activation | A0A075BWY1 | C3YKE8 | - | - | F1NS46 |
| 22 | LSKKR R | 18 | P22314 | Ubiquitin-like modifier-activating enzyme 1 | Catalyzes the first step in ubiquitin conjugation to mark cellular proteins for degradation through the ubiquitin-proteasome system | Q6P9P3 | C3ZZI8 | Q6P285 | K7F4G1 | F1NPI6 |

| | | | | | | | | | | |
|---|---|---|---|---|---|---|---|---|---|---|
| 23 | RP LQSEGAAALV TKGCQRLAAQ GARPEAPKRK WAEDGGDAPS PSKR | 19 | Q96KS0 | Egl nine homolog 2 | Cellular oxygen sensor that catalyzes, under normoxic conditions, the post-translational formation of 4-hydroxyproline in hypoxia-inducible factor (HIF) alpha proteins | E7F8S1 | C3YH16 | A0A1L8F3X2 | K7GGI0 | E1C8C0 |
| 24 | RKRRKR | 20 | Q6X4W1 | NMDA receptor synaptonuclear signaling and neuronal migration factor | Couples NMDA-sensitive glutamate receptor signaling to the nucleus and triggers long-lasting changes in the cytoarchitecture of dendrites and spine synapse processes | B8PYG1 | C3ZTG8 | A0A1L8F0Y0 | A0A1V4J7D7 | - |
| 25 | NKRKR K | 21 | Q9HAJ7 | Histone deacetylase complex subunit SAP30L | unctions as transcription repressor | Q6NYV5 | C3YZU5 | A0A1L8HU31 | - | R4GLF5 |
| 26 | PP ARRRR | 22 | Q8NAG6 | Ankyrin repeat and LEM domain-containing protein 1 | Endonuclease that probably plays a role in the DNA damage response and DNA repair. | Q6NUU1 | C3YQ02 | A0A1L8HWS5 | - | - |
| 27 | KKPKW DDFKKKKK | 23 | Q15397 | Pumilio homolog 3 | Inhibits the poly(ADP-ribosyl)ation activity of PARP1 and the degradation of PARP1 by CASP3 following genotoxic stress. Binds to double-stranded RNA or DNA without sequence specificity | X1WGX5 | C3YCI8 | Q6DFI0 | K7GDL9 | E1C1R5 |
| 28 | RKRK | 24 | Q6NTE8 | MRN complex-interacting protein | Plays a role in the cellular response to DNA damage and the maintenance of genome stability through its association with the MRN damage-sensing complex | Q32PP1 | - | - | - | - |
| 29 | PRRRVQR KR | 25 | Q5TAQ9 | DDB1- and CUL4-associated factor 8 | May function as a substrate receptor for CUL4-DDB1 E3 ubiquitin-protein ligase complex | F1QF18 | C3XZJ2 | Q6NRH1 | K7FA09\| | A0A1D5PZY7 |
| 30 | KRK | 26 | Q5T5X7 | BEN domain-containing protein 3 | Transcriptional repressor which associates with the NoRC (nucleolar remodeling complex) complex and plays a key role in repressing rDNA transcription | B7ZD94 | - | A0A1L8G991 | - | F1NMS3 |
| 31 | RKVKKLR | 27 | Q86TI2 | Dipeptidyl peptidase 9 | Dipeptidyl peptidase that cleaves off N-terminal dipeptides from proteins having a Pro or Ala residue at position 2 | B0V1L8 | C3YDM8 | Q6GR22 | K7FG08 | A0A1D5P419 |
| 32 | KKR K | 28 | Q5T3J3 | Ligand-dependent nuclear receptor-interacting factor 1 | Together with SMCHD1, involved in chromosome X inactivation in females by promoting the compaction of | E7F853 | - | A0A1L8HBT1 | K7G507 | - |
| 33 | KKRK | | | | | | | | | |
| 34 | NF QDDPHAGL | 29 | O43257 | Zinc finger HIT domain-containing protein 1 | Seems to play a role in p53-mediated apoptosis induction. Binds to NR1D2 and relieves it of its inhibitory effect on the transcription of APOC3 without affecting its DNA-binding activity | Q561S5 | - | A0A1L8H473 | - | - |
| 35 | KRGRKREDK ED | 30 | Q9NVM9 | Integrator complex subunit 13 | Crucial regulator of the mitotic cell cycle and development. At prophase, required for dynein anchoring to the nuclear envelope important for proper centrosome-nucleus coupling. At G2/M phase, may be required for proper spindle formation and execution of cytokinesis. Probable component of the Integrator (INT) complex, a complex involved in the small nuclear RNAs (snRNA) U1 and U2 transcription and in their 3'-box-dependent processing | Q7ZVZ8 | C3YV85 | Q6GLY5 | K7GIR3 | F1P1V8 |
| 36 | KKPRKCDLTPFLVLKARKK | 31 | Q9NR20 | Dual specificity tyrosine-phosphorylation-regulated kinase 4 | Possible non-essential role in spermiogenesis | A0A0R4ITS5 | C3YLM7 | Q5PPW5 | K7F6T0 | Q5ZIU3 |
| 37 | KKE IARQMKAYEK QRGTRREEQK E | 32 | Q9Y314 | Nitric oxide synthase-interacting protein | E3 ubiquitin-protein ligase that is essential for proper development of the forebrain, the eye, and the face. Negatively regulates nitric oxide production by inducing NOS1 and NOS3 translocation to actin cytoskeleton and inhibiting their enzymatic activity | B2GNS3 | C3Y5M1 | A0A1L8FGZ1 | - | - |
| 38 | RPKGKTLQ KRKPK | 33 | Q6ZN17 | Protein lin-28 homolog B | Suppressor of microRNA (miRNA) biogenesis, including that of let-7 and possibly of miR107, miR-143 and miR-200c. Binds primary let-7 transcripts (pri-let-7), including pri-let-7g and pri-let-7a-1, and sequester them in the nucleolus, away from the microprocessor complex, hence preventing their processing into mature miRNA | S0BDM9 | - | A0A1L8G938 | K7F3U2 | A0A1D5PHV1 |
| 39 | RKR QR | 34 | O75937 | DnaJ homolog subfamily C member 8 | Suppresses polyglutamine (polyQ) aggregation of ATXN3 in neuronal cells | Q1ED27 | C3XW14 | A0A1L8HE93 | K7GI79 | F1ND15 |
| 40 | KRER | | | | | | | | | |
| 41 | R HRKIYTKKHH | 35 | Q8NFU5 | Inositol polyphosphate multikinase | Inositol phosphate kinase with a broad substrate specificity. Has a preference for inositol 1,4,5-trisphosphate (Ins(1,4,5)P3) and inositol 1,3,4,6-tetrakisphosphate (Ins(1,3,4,6)P4) | Q6PBN6 | C3Z3L6 | Q6GQ51 | K7G124 | F1NFX3 |

| | | | | | | | | | | |
|---|---|---|---|---|---|---|---|---|---|---|
| 42 | KRRRCA | 36 | Q53H80 | Akirin-2 | Required for the innate immune response. Downstream effector of the Toll-like receptor (TLR), TNF and IL-1 beta signaling pathways leading to the production of IL-6 | A5WVY2 | C3YH63 | Q6GQB5 | - | E1C249 |
| 43 | KK | 37 | Q9C0C6 | CLOCK-interacting pacemaker | Transcriptional repressor which may act as a negative-feedback regulator of CLOCK-ARNTL/BMAL1 transcriptional activity in the circadian-clock mechanism | E7FAN7 | - | A0A1L8F9G1 | - | F1NAE8 |
| 44 | RRRQKQKGGA SRRR | 38 | Q9BWC9 | Coiled-coil domain-containing protein 106 | Promotes the degradation of p53/TP53 protein and inhibits its transactivity | Q5EB04 | - | - | - | - |
| 45 | PKKRRKKK | 39 | Q12796 | Proline-rich nuclear receptor coactivator 1 | Nuclear receptor coactivator. May play a role in signal transduction | Q5TZ15 | - | - | - | Q5ZII5 |
| 46 | SRERK | 40 | O75467 | Zinc finger protein 324A | May be involved in transcriptional regulation. May be involved in regulation of cell proliferation | A2RUZ2 | C3XSR4 | A0A1L8HBB8 | K7FDC2 | A0A1D5PJZ9 |
| 47 | PLLKKIKQ | 41 | Q708E1 | C-myb v-myb myeloblastosis viral oncogene homologue (avian), exon 1 and joined CDS | DNA binding | F1QP24 | C3ZI75 | Q05935 | K7FZH8 | P01103 |
| 48 | KRPMNAFMVWAQAARRK | 42 | P48436 | Transcription factor SOX-9 | Transcriptional regulator that plays a role in chondrocytes differentiation and skeletal development | Q9DFH2 | C3Z730 | B7ZR65 | K7G5M4 | P48434 |
| 49 | PRRRK | | | | | | | | | |
| 50 | PPQKKIKS | 43 | Q9UMQ5 | N-myc protein | DNA-binding transcription factor activity | Q6PBB3 | - | - | - | - |
| 51 | PAAKRVKLD | 44 | P01106 | Myc proto-oncogene protein | Transcription factor that binds DNA in a non-specific manner, yet also specifically recognizes the core sequence 5'-CAC[GA]TG-3'. Activates the transcription of growth-related genes. Binds to the VEGFA promoter, promoting VEGFA production and subsequent sprouting angiogenesis | B2GQB7 | C3Y235 | Q2TAU1 | K7GG15 | F1NXY9 |
| 52 | QRKRQK | 45 | P19838 | Nuclear factor NF-kappa-B p105 subunit | NF-kappa-B is a pleiotropic transcription factor present in almost all cell types and is the endpoint of a series of signal transduction events that are initiated by a vast array of stimuli related to many biological processes such as inflammation, immunity, differentiation, cell growth, tumorigenesis and apoptosis | A0A0R4IN79 | C3XR39 | A0A1L8F873 | K7FNJ5 | Q04861 |
| 53 | RRSMKRK | 46 | P11473 | Vitamin D3 receptor | Nuclear receptor for calcitriol, the active form of vitamin D3 which mediates the action of this vitamin on cells. Enters the nucleus upon vitamin D3 binding where it forms heterodimers with the retinoid X receptor/RXR. The VDR-RXR heterodimers bind to specific response elements on DNA and activate the transcription of vitamin D3-responsive target genes. Recruited to promoters via its interaction with BAZ1B/WSTF which mediates the interaction with acetylated histones, an essential step for VDR-promoter association. Plays a central role in calcium homeostasis | Q1L673 | - | B7ZPA5 | - | A0A1D5NUY7 |
| 54 | SKRVAKRKL | 47 | P10827 | Thyroid hormone receptor alpha | Nuclear hormone receptor that can act as a repressor or activator of transcription. High affinity receptor for thyroid hormones, including triiodothyronine and thyroxine | A0A286Y807 | C3YYX7 | A0A1L8EKN8 | K7FLZ6 | A0A1D5PCG7 |
| 55 | VNEAFETLKRC | 48 | P15172 | Myoblast determination protein 1 | Acts as a transcriptional activator that promotes transcription of muscle-specific target genes and plays a [illegible] | Q90477 | - | Q6GN48 | E5RLQ9 | P16075 |
| 56 | CKRKTTNADRRKA | | | | | | | | | |
| 57 | RRMKWKK | 49 | Q9UMQ3 | Homeobox protein BarH-like 2 | Transcription factor. Binds optimally to the DNA consensus sequence 5'-YYTAATGRTTTTY-3' | F6NKZ9 | C3Y0U7 | A0A1L8FLC4 | K7F6T6 | Q9W694 |
| 58 | KRKx{11}KKKSKK | 50 | P09874 | Poly [ADP-ribose] polymerase 1 | Involved in the base excision repair (BER) pathway, by catalyzing the poly(ADP-ribosyl)ation of a limited number of acceptor proteins involved in chromatin architecture and in DNA metabolism. | Q5RHR0 | C3Z2P3 | Q2NLA7 | - | A0A1L1RQW |
| 59 | KRMRNRIAASKCRKRKL | 51 | P05412 | Transcription factor AP-1 | Transcription factor that recognizes and binds to the enhancer heptamer motif 5'-TGA[CG]TCA-3'. Promotes activity of NR5A1 when phosphorylated by HIPK3 leading to increased steroidogenic gene expression upon cAMP signaling pathway stimulation. Involved in activated KRAS-mediated transcriptional activation of USP28 in colorectal cancer (CRC) cells | B0V1Q6 | C3YQP4 | A0JMS2 | K7EY56 | P18870 |

| | | | | | | | | | | |
|---|---|---|---|---|---|---|---|---|---|---|
| 60 | KHRKHPG | 52 | P46776 | 60S ribosomal protein L27a | RNA binding | Q6P2M0 | C3ZKJ7 | P47830 | - | A0A1D5PSU0 |
| 61 | HKKKKIRTSPTFTTPKTLRLRR QPKYPRKSAPRRNKLDHY | 53 | P62750 | 60S ribosomal protein L23a | Binds a specific region on the 26S rRNA. May promote p53/TP53 degradation possibly through the stimulation of MDM2-mediated TP53 polyubiquitination | Q6Q416 | C3YD43 | A0A1L8H4S4 | - | E1BS06 |
| 62 | KKKHPDASVNFSEFSKK | 54 | P09429 | High mobility group protein B1 | In the nucleus is one of the major chromatin-associated non-histone proteins and acts as a DNA chaperone | X1WFI9 | C3YZU6 | Q6GNQ5 | K7FJB3 | Q9YH06 |
| 63 | EKSKKKK | | | | | | | | | |
| 64 | RKG AKSRLSSSKS KSRTSPYPQY | 55 | P81133 | Single-minded homolog 1 | Transcriptional factor that may have pleiotropic effects during embryogenesis and in the adult. | F1QMF7 | - | A0A1L8G2Y4 | - | E1C5G3 |
| 65 | KGKYK EGDTGGPAVW KTRLR | 56 | Q00978 | Interferon regulatory factor 9 | Transcription factor that mediates signaling by type I IFNs (IFN-alpha and IFN-beta) | F1Q573 | C3XS91 | A0A1L8HPG8 | K7FYQ2 | Q5ZJM5 |
| 66 | CRLRKC YEAGMTLGAR KLKKL | 57 | D3YPQ2 | Androgen receptor isoform 1 transcript variant 1 | DNA binding | A4GVF3 | - | - | - | - |
| 67 | CRLQKCFE VGMSKESVRN DRNKKKKEVP KP | 58 | Q6I9R7 | RARA protein | DNA binding | T2DKU4 | Q8MX80 | Q92019 | K7F2Z9 | Q90966 |
| 68 | RRRHSDEN DGGQPHKRRK | 59 | Q09161 | Nuclear cap-binding protein subunit 1 | Component of the cap-binding complex (CBC), which binds cotranscriptionally to the 5'-cap of pre-mRNAs and is involved in various processes such as pre-mRNA splicing, translation regulation, nonsense-mediated mRNA decay, RNA-mediated gene silencing (RNAi) by microRNAs (miRNAs) and mRNA export | UPI0007EEE498 | C3Y2C6 | A0A1L8HX89 | - | Q5ZJZ6 |
| 69 | GKK RKRPHVFESN PSIRKR | 60 | Q16656 | Nuclear respiratory factor 1 | Transcription factor that activates the expression of the EIF2S1 (EIF2-alpha) gene | Q90X44 | C3ZAI3 | A0A1L8GZK2 | K7GFU4 | Q5ZL67 |
| 70 | KKPKR | 61 | Q9Y6D6 | Brefeldin A-inhibited guanine nucleotide-exchange protein 1 | Promotes guanine-nucleotide exchange on ARF1 and ARF3 | E7FGL2 | C3Y295 | A0A1L8FT76 | K7FVJ2 | A0A1D5P6U6 |
| 71 | KP KKLKV | 62 | P55263 | Adenosine kinase | ATP dependent phosphorylation of adenosine and other related nucleoside analogs to monophosphate derivatives. Serves as a potential regulator of concentrations of extracellular adenosine and intracellular adenine nucleotides. | Q6P3J9 | C3XQH1 | Q6DJM1 | - | Q5ZMK9 |
| 72 | RQHMRK AVPFQVAQK | 63 | Q8WTP8 | Apoptosis-enhancing nuclease | Exonuclease with activity against single- and double-stranded DNA and RNA | E7F5Q4 | C3YLI2 | A0A1L8H0Y1 | K7GFG9 | A0A1L1RN59 |
| 73 | MDSLLMNRRK FLYQFKNVRW AKGRRETYLC | 64 | Q9GZX7 | Single-stranded DNA cytosine deaminase | Involved in somatic hypermutation (SHM), gene conversion, and class-switch recombination (CSR) in B-lymphocytes by deaminating C to U during transcription of Ig-variable (V) and Ig-switch (S) region DNA. | Q5YF88 | - | Q4VUI3 | - | E1BVI0 |
| 74 | KRGKKGAV AED | 65 | P27695 | DNA-(apurinic or apyrimidinic site) lyase | The two major activities in DNA repair and redox regulation of transcriptional factors. | F1QTR9 | C3YX76 | Q6IP62 | - | - |
| 75 | KLV KPKNTKMKTK LRTNPY | 66 | Q14190 | Single-minded homolog 2 | Transcription factor that may be a master gene of CNS development in cooperation with Arnt. It may have pleiotropic effects in the tissues expressed during development | C1JIE2 | C3XTH0 | A0A1L8HCG5 | - | E1BSJ1 |
| 76 | RRVRLK | 67 | Q13568 | Interferon regulatory factor 5 | Transcription factor involved in the induction of interferons IFNA and INFB and inflammatory cytokines upon virus infection. Activated by TLR7 or TLR8 signaling | B0S682 | C3ZAX5 | A0A1L8GTJ1 | K7F6Y8 | E1BUT7 |
| 77 | KKHRRRPSKK KRHWKPY | 68 | O94992 | Protein HEXIM1 | Transcriptional regulator which functions as a general RNA polymerase II transcription inhibitor | A5D8S8 | C3Y140 | - | - | - |
| 78 | YWE ECPREGKSFK AKYKLVNHIR VH | 69 | O60481 | Zinc finger protein ZIC 3 | Acts as transcriptional activator. Required in the earliest stages in both axial midline development and left-right (LR) asymmetry specification. Binds to the minimal GLI-consensus sequence 5'-GGGTGGTC-3' | Q9IAT0 | - | O57311 | K7FBG1 | A0A1D5PRL4 |
| 79 | PFPGCGKIFA RSENLKIHKR TH | | | | | | | | | |
| 80 | RV RYNPYTTRPN RRGDTWHDRD RIHVTVRRDR | 70 | Q9UBU9 | Nuclear RNA export factor 1 | Involved in the nuclear export of mRNA species bearing retroviral constitutive transport elements (CTE) and in the export of mRNA from the nucleus to the cytoplasm (TAP/NFX1 pathway) | Q0P3Y8 | C3ZF40 | Q6IRQ2 | - | - |
| 81 | KSDKFL IIPLHSLM | 71 | Q9H2U1 | ATP-dependent RNA helicase DHX36 | Proposed to have a global role in regulating mRNA expression including transcriptional regulation and mRNA stability. Binds with high affinity to and resolves tetramolecular RNA and DNA quadruplex structures | A2BIE5 | C3Y332 | A0A1L8GAK8 | K7FY91 | A0A1D5PHB6 |
| 82 | KRARCRR HQR | 72 | Q8NHV9 | Rhox homeobox family member 1 | Transcription factor maybe involved in reproductive processes. Modulates expression of target genes encoding proteins involved in processes relevant to spermatogenesis | Q7SZN8 | - | - | - | - |

| | | | | | | | | | |
|---|---|---|---|---|---|---|---|---|---|
| 83 | RRRx{12}KRPRR | 73 | O14746 | Telomerase reverse transcriptase | Telomerase is a ribonucleoprotein enzyme essential for the replication of chromosome termini in most eukaryotes | A2THE9 | - | Q9DE32 | - | A0A1D5PY42 |
| 84 | RR KEKKRRRK | 74 | Q96B42 | Transmembrane protein 18 | Transcription repressor. Sequence-specific ssDNA and dsDNA binding protein, with preference for GCT end CTG repeats | Q641M3 | C3XYA5 | A0A1L8G500 | - | Q5F410 |
| 85 | KPSKGQ KRHLSTCDGQ NPPKK | 75 | O15226 | NF-kappa-B-repressing factor | Interacts with a specific negative regulatory element (NRE) 5'-AATTCCTCTGA-3' to mediate transcriptional repression of certain NK-kappa-B responsive genes | A0A0R4IUL8 | C3YZJ2 | A0A1L8F334 | - | F1NL52 |
| 86 | KYRKGAC ENCGAMTHKK KDCFER | 76 | O95391 | Pre-mRNA-splicing factor SLU7 | Participates in the second catalytic step of pre-mRNA splicing, when the free hydroxyl group of exon I attacks the 3'-splice site to generate spliced mRNA and the excised lariat intron | F1RCR0 | C3XZQ3 | A0A1L8GX04 | - | Q5ZIG2 |
| 87 | KKRKH | 77 | Q9Y2G1 | Myelin regulatory factor | Myelin regulatory factor: Constitutes a precursor of the transcription factor. Mediates the autocatalytic cleavage [illegible] | UPI000383813F | C3Y630 | Q66IV1 | - | E1C867 |
| 88 | R KKGK | | | | | | | | | |
| 89 | RRAMK RNARLRCPFR KGACEITRKT RR | 78 | O75469 | Nuclear receptor subfamily 1 group I member 2 | Nuclear receptor that binds and is activated by variety of endogenous and xenobiotic compounds. Transcription factor that activates the transcription of multiple genes involved in the metabolism and secretion of potentially harmful xenobiotics, drugs and endogenous compounds | B0S5T9 | - | A0A1L8HIC5 | - | Q9DFH3 |
| 90 | LKRKLQR | 79 | P26367 | Paired box protein Pax-6 | Competes with PAX4 in binding to a common element in the glucagon, insulin and somatostatin promoters | A0A0R4IXU2 | C3ZKE8 | A0A1L8GJ29 | K7G268 | O42348 |
| 91 | RKKRGGRRYR KMKER | 80 | Q8WWY3 | U4/U6 small nuclear ribonucleoprotein Prp31 | Involved in pre-mRNA splicing as component of the spliceosome. Required for the assembly of the U4/U5/U6 tri-snRNP complex, one of the building blocks of the spliceosome | B2GS36 | C3Z7L6 | Q5U5C5 | - | - |
| 92 | RRKNYPTL ANIERKKKLK | 81 | Q9UHK0 | Nuclear fragile X mental retardation-interacting protein 1 | Binds RNA | A8KB24 | C3YJZ2 | A0A1L8HHI9 | K7FJF1 | E1C0G7 |
| 93 | RKRTAPE DFLGKGPTKK | 82 | Q96FV9 | THO complex subunit 1 | Required for efficient export of polyadenylated RNA. Acts as component of the THO subcomplex of the TREX complex which is thought to couple mRNA transcription, processing and nuclear export, and which specifically associates with spliced mRNA and not with unspliced pre-mRNA | Q7SYB2 | C3Y1Y4 | Q498H9 | - | - |
| 94 | RKRKK | 83 | P20264 | POU domain, class 3, transcription factor 3 | Transcription factor that acts synergistically with SOX11 and SOX4. Plays a role in neuronal development. Is implicated in an enhancer activity at the embryonic met-mesencephalic junction; the enhancer element contains the octamer motif (5'-ATTTGCAT-3') | A2VD21 | C3ZW98 | A0A1L8H8P3 | K7EWD4 | - |
| 95 | KRVL QVWFQNARAK FRR | 84 | P50458 | LIM/homeobox protein Lhx2 | Acts as a transcriptional activator. Stimulates the promoter of the alpha-glycoprotein gene. Transcriptional regulatory protein involved in the control of cell differentiation in developing lymphoid and neural cell types | B0R107 | C3ZG21 | A0A1L8F6B6 | K7FB66 | A0A1D5PPZ9 |
| 96 | RQRK | 85 | Q8IWS0 | PHD finger protein 6 | Transcriptional regulator that associates with ribosomal RNA promoters and suppresses ribosomal RNA (rRNA) transcription | Q803W6 | C3ZZY1 | A0A1L8F3A3 | - | V9GVP7 |
| 97 | PQPKKKP | 86 | P04637 | Cellular tumor antigen p53 | Acts as a tumor suppressor in many tumor types; induces growth arrest or apoptosis depending on the physiological circumstances and cell type. Involved in cell cycle regulation as a trans-activator that acts to negatively regulate cell division by controlling a set of genes required for this process | G1K2L5 | C3XPU2 | P07193 | K7G3P4 | A0A167VDT2 |
| 98 | LEKKVKKKFDW | 87 | P60900 | Proteasome subunit alpha type-6 | Component of the 20S core proteasome complex involved in the proteolytic degradation of most intracellular proteins | Q7SXN6 | C3XZ66 | Q6AZR7 | - | F1NEQ6 |
| 99 | YLTQETNKVETYKEQPLKTPG KKKKGKP | 88 | P12272 | Parathyroid hormone-related protein | Neuroendocrine peptide which is a critical regulator of cellular and organ growth, development, migration, differentiation and survival and of epithelial calcium ion transport. Regulates endochondral bone development and epithelial-mesenchymal interactions during the formation of the mammary glands and teeth | Q1L5E7 | - | A0A1L8GR30 | - | P17251 |

| | | | | | | | | | | |
|---|---|---|---|---|---|---|---|---|---|---|
| 100 | RPRRK | 89 | Q05066 | Sex-determining region Y protein | Promotes DNA bending. SRY HMG box recognizes DNA by partial intercalation in the minor groove. Also | Q6EJB7 | - | - | - | - |
| 101 | KRPMNAFIVWSRDQRRK | | | | | | | | | |
| 102 | RRERNKMAAAKCRNRRR | 90 | P01100 | Proto-oncogene c-Fos | Nuclear phosphoprotein which forms a tight but non-covalently linked complex with the JUN/AP-1 transcription factor. In the heterodimer, FOS and JUN/AP-1 basic regions each seems to interact with symmetrical DNA half sites. | A2CEM8 | C3Y0C5 | Q91639 | K7GHV7 | F1NFV6 |
| 103 | KKKRERLD | 91 | P11171 | Protein 4.1 | Protein 4.1 is a major structural element of the erythrocyte membrane skeleton. It plays a key role in regulating membrane physical properties of mechanical stability and deformability by stabilizing spectrin-actin interaction. Recruits DLG1 to membranes. Required for dynein-dynactin complex and NUMA1 recruitment at the mitotic cell cortex during anaphase | E7F634 | C3ZA69 | P11434 | K7FVB5 | P12264 |
| 104 | KRPACTLKPECVQQLLVCSQEAKK | 92 | P32320 | Cytidine deaminase | This enzyme scavenges exogenous and endogenous cytidine and 2'-deoxycytidine for UMP synthesis | Q6P0R4 | C3XPZ1 | A0A1L8FHN2 | K7FC96 | E1C6V1 |
| 105 | KRQRx{20}KKSKK | 93 | P49454 | Centromere protein F | Required for kinetochore function and chromosome segregation in mitosis. Required for kinetochore localization of dynein, LIS1, NDE1 and NDEL1 | F1Q7X5 | - | A0A1L8G630 | - | A0A1L1RMT7 |
| 106 | KRSAEGSNPPKPLKKLR | 94 | P06400 | Retinoblastoma-associated protein | Key regulator of entry into cell division that acts as a tumor suppressor. Promotes G0-G1 transition when phosphorylated by CDK3/cyclin-C. Acts as a transcription repressor of E2F1 target genes | B3DI38 | C3XU15 | A0A1L8EML3 | K7FQ20 | Q90600 |
| 107 | KRKCSQTQCPRKVIK | 95 | P29590 | Protein PML | Functions via its association with PML-nuclear bodies (PML-NBs) in a wide range of important cellular processes, including tumor suppression, transcriptional regulation, apoptosis, senescence, DNA damage response, and viral defense mechanisms | F1RBJ0 | - | - | - | A0A1L1RZK6 |
| 108 | PGRKRKW | 96 | Q9HAN9 | Nicotinamide/nicotinic acid mononucleotide adenylyltransferase 1 | Catalyzes the formation of NAD+ from nicotinamide mononucleotide (NMN) and ATP | E9QH56 | C3Y2B3 | A0A1L8FKV8 | K7G6T7 | R4GGL5 |
| 109 | ILIKRRQQ KIRKYTMRRL LQ | 97 | P04626 | Receptor tyrosine-protein kinase erbB-2 | Protein tyrosine kinase that is part of several cell surface receptor complexes, but that apparently needs a coreceptor for ligand binding. Essential component of a neuregulin-receptor complex, although neuregulins do not interact with it alone | Q7SZF7 | C3YZ48 | A0A1L8FRS8 | - | A0A1D5PEH0 |
| 110 | NKL PRRRVNIFGE ELERLLKKKY | 98 | P50616 | Protein Tob1 | Anti-proliferative protein; the function is mediated by association with deadenylase subunits of the CCR4-NOT complex | Q7SZR1 | C3Z7P3 | A0A1L8EUN5 | K7FHV8 | Q90VW0 |
| 111 | RRQQEQLR YEQERRPGRR | 99 | Q14151 | Scaffold attachment factor B2 | Binds to scaffold/matrix attachment region (S/MAR) DNA. Can function as an estrogen receptor corepressor and can also inhibit cell proliferation. | Q803X6 | C3Z0S4 | A0A1L8HWX4 | K7FD64 | H9KZ12 |
| 112 | ELCRRRSLLE LHKRRKALTE | 100 | P53350 | Serine/threonine-protein kinase PLK1 | Serine/threonine-protein kinase that performs several important functions throughout M phase of the cell cycle, including the regulation of centrosome maturation and spindle assembly, the removal of cohesins from chromosome arms, the inactivation of anaphase-promoting complex/cyclosome (APC/C) inhibitors, and the regulation of mitotic exit and cytokinesis | Q4KMI8 | C3ZJT7 | P70032 | K7FID3 | F1NB57 |
| 113 | SKTFAMK ILKKRHIVDT RQQ | 101 | Q13976-2 | cGMP-dependent protein kinase 1 | Serine/threonine protein kinase that acts as key mediator of the nitric oxide (NO)/cGMP signaling pathway | A0A286Y9K1 | C3Z4R1 | Q6GPV8 | - | E1C4W9 |
| 114 | RKKEA PGPREELRSR GR | 102 | P54259 | Atrophin-1 | Transcriptional corepressor. Recruits NR2E1 to repress transcription | UPI00038385C7 | - | A0A1L8FIG3 | K7G1I3 | - |
| 115 | RKPDRRKR SR | 103 | Q92560 | Ubiquitin carboxyl-terminal hydrolase BAP1 | Deubiquitinating enzyme that plays a key role in chromatin by mediating deubiquitination of histone H2A and HCFC1. Catalytic component of the PR-DUB complex, a complex that specifically mediates deubiquitination of histone H2A monoubiquitinated at 'Lys-119' (H2AK119ub1) | F1QJ12 | C4A0D9 | A0A1L8GHI0 | - | Q5F3N6 |
| 116 | KRFRREFGD SKRVFGERRR ND | 104 | Q9NRA8 | Eukaryotic translation initiation factor 4E transporter | Nucleoplasmic shuttling protein, which inhibits translation initiation. Mediates the nuclear import of EIF4E by a piggy-back mechanism | E7FDR0 | C3YCP5 | A0A1L8I094 | - | E1BTK8 |

| | | | | | | | | | | |
|---|---|---|---|---|---|---|---|---|---|---|
| 117 | HRKR | 105 | Q7Z6G8 | Ankyrin repeat and sterile alpha motif domain-containing protein 1B | Ephrin receptor binding | A0A0R4IE55 | C3ZFZ7 | A0A1L8H6M2 | - | A0A1D5PKC1 |
| 118 | KRASEDT TSGSPPKKSS AGPKR | 106 | Q9BZZ5 | Apoptosis inhibitor 5 | Antiapoptotic factor that may have a role in protein assembly. Negatively regulates ACIN1. By binding to ACIN1, it suppresses ACIN1 cleavage from CASP3 and ACIN1-mediated DNA fragmentation. Also known to efficiently suppress E2F1-induced apoptosis. Its depletion enhances the cytotoxic action of the chemotherapeutic drugs | R4GDP5 | C4A049 | Q6DDM4 | - | Q5ZMW3 |
| 119 | RRPR RRTFPGVASR RNPERR | 107 | Q9BWT1 | Cell division cycle-associated protein 7 | Participates in MYC-mediated cell transformation and apoptosis; induces anchorage-independent growth and clonogenicity in lymphoblastoid cells. Insufficient to induce tumorigenicity when overexpressed but contributes to MYC-mediated tumorigenesis. May play a role as transcriptional regulator | B0S4Y9 | C3Y254 | A0JMU9 | K7FL67 | A0A1D5PK49 |
| 120 | KEL LKHKFILRNA KK | 108 | Q9Y6E0 | Serine/threonine-protein kinase 24 | Serine/threonine-protein kinase that acts on both serine and threonine residues and promotes apoptosis in response to stress stimuli and caspase activation | A0A0R4IN78 | C3YYD0 | A0A1L8H969 | K7FJN7 | A0A1D5NWG8 |
| 121 | KRRK | 109 | Q14678 | KN motif and ankyrin repeat domain-containing protein 1 | Involved in the control of cytoskeleton formation by regulating actin polymerization | E9QCJ6 | C3ZZY7 | A0A1L8HWJ6 | K7G0R3 | F1NA35 |
| 122 | N ESTLKSIMKK KDGNKDSNGA KK | | | | | | | | | |
| 123 | KSKKRIR | 110 | Q6NYC1 | Bifunctional arginine demethylase and lysyl-hydroxylase JMJD6 | Dioxygenase that can both act as a histone arginine demethylase and a lysyl-hydroxylase | Q6PFM0 | C3ZFA3 | A0A1L8EUJ7 | K7FSE8 | Q5ZMK5 |
| 124 | KRKYR | | | | | | | | | |
| 125 | RRKKRR | | | | | | | | | |
| 126 | PKRRK | | | | | | | | | |
| 127 | KRRP | | | | | | | | | |
| 128 | RKE KVKRE | 111 | P34947 | G protein-coupled receptor kinase 5 | Serine/threonine kinase that phosphorylates preferentially the activated forms of a variety of G-protein-coupled receptors (GPCRs | Q1XHM0 | C3Y5N3 | Q6DCW4 | K7FMD2 | Q5ZKL2 |
| 129 | RQQRKR | 112 | Q14980 | Nuclear mitotic apparatus protein 1 | Microtubule (MT)-binding protein that plays a role in the formation and maintenance of the spindle poles and the alignement and the segregation of chromosomes during mitotic cell division | A0A0R4IH76 | - | A0A1L8HJJ6 | - | D8MIU8 |
| 130 | KRKRKNSR VTFSEDDEI | 113 | Q12972 | Nuclear inhibitor of protein phosphatase 1 | Inhibitor subunit of the major nuclear protein phosphatase-1 (PP-1). It has RNA-binding activity but does not cleave RNA and may target PP-1 to RNA- | A5PF58 | - | Q6P9H8 | K7GAA8 | F1N9I1 |
| 131 | RFRNMVQTAV VPVKKKRVEG | | | | | | | | | |
| 132 | KRKR | 114 | Q6DJT9 | Zinc finger protein PLAG1 | Transcription factor whose activation results in up-regulation of target genes, such as IGFII, leading to uncontrolled cell proliferation: when overexpressed in cultured cells, higher proliferation rate and transformation are observed | Q29YB2 | C3ZWC1 | A0A1L8FZ09 | K7GIK1 | Q58NQ5 |
| 133 | KKKYT R | 115 | Q13177 | Serine/threonine-protein kinase PAK 2 | Serine/threonine protein kinase that plays a role in a variety of different signaling pathways including cytoskeleton regulation, cell motility, cell cycle progression, apoptosis or proliferation. Acts as downstream effector of the small GTPases CDC42 and RAC1 | Q6DG42 | C3Z873 | Q6PAZ2 | K7FAU6 | E1C3P6 |
| 134 | RKSK | 116 | Q9H2X6 | Homeodomain-interacting protein kinase 2 | Serine/threonine-protein kinase involved in transcription regulation, p53/TP53-mediated cellular [illegible] | F1Q4N0 | C3Y5H6 | A0A1L8H6D1 | K7GA98 | F1P4G3 |
| 135 | KRVK | | | | | | | | | |
| 136 | KKAILKKKEE K | 117 | O00141 | Serine/threonine-protein kinase Sgk1 | Serine/threonine-protein kinase which is involved in the regulation of a wide variety of ion channels, membrane transporters, cellular enzymes, transcription factors, neuronal excitability, cell growth, proliferation, survival, migration and apoptosis | Q7ZTW4 | C3Y291 | Q6GPN6 | K7G4F8 | Q6U1I9 |
| 137 | RGKK K | 118 | Q9UBP0 | Spastin | ATP-dependent microtubule severing protein that specifically recognizes and cuts microtubules that are [illegible] | Q6NW58 | C3YR60 | Q6AZT2 | K7EXQ8 | Q5ZK92 |
| 138 | RK KK | | | | | | | | | |
| 139 | KRP R | 119 | Q9UQ84 | Exonuclease 1 | 5'->3' double-stranded DNA exonuclease which may also possess a cryptic 3'->5' double-stranded DNA exonuclease activity | Q803U7 | C3Z4R4 | Q9W6K2 | K7G1H9 | E1C1G7 |

| | | | | | | | | | |
|---|---|---|---|---|---|---|---|---|---|
| 140 | KLRRR | 120 | O95239 | Chromosome-associated kinesin KIF4A | Motor protein that translocates PRC1 to the plus ends of interdigitating spindle microtubules during the metaphase to anaphase transition, an essential step for the formation of an organized central spindle midzone and midbody and for successful cytokinesis. May play a role in mitotic chromosomal positioning and bipolar spindle stabilization | F8W409 | C3Z125 | Q91784 | K7FYR1 | O95239 |
| 141 | K VKTKKSTK | 121 | P42338 | Phosphatidylinositol 4,5-bisphosphate 3-kinase catalytic subunit beta isoform | Phosphoinositide-3-kinase (PI3K) that phosphorylates PtdIns (Phosphatidylinositol), PtdIns4P (Phosphatidylinositol 4-phosphate) and PtdIns(4,5)P2 (Phosphatidylinositol 4,5-bisphosphate) to generate phosphatidylinositol 3,4,5-trisphosphate (PIP3). | F1QAD7 | C3XYJ1 | A0A1L8G3Z7 | K7F3A1 | E1C093 |
| 142 | TKRPRW | 122 | Q86W56 | Poly(ADP-ribose) glycohydrolase | Poly(ADP-ribose) synthesized after DNA damage is only present transiently and is rapidly degraded by poly(ADP-ribose) glycohydrolas | E7F2B6 | C3ZES4 | Q4KLP9 | K7FAS7 | F1NY76 |
| 143 | KR KTKISRKTR | 123 | O00443 | Phosphatidylinositol 4-phosphate 3-kinase C2 domain-containing subunit alpha | Generates phosphatidylinositol 3-phosphate (PtdIns3P) and phosphatidylinositol 3,4-bisphosphate (PtdIns(3,4)P2) that act as second messengers | F1QWN6 | C3ZMJ4 | A0A1L8GQ56 | - | E1C1G0 |
| 144 | KQKQRF EEKRFK | 124 | Q8NHY2 | E3 ubiquitin-protein ligase COP1 | E3 ubiquitin-protein ligase that mediates ubiquitination and subsequent proteasomal degradation of target | A4QP94 | C3Y342 | A0A1L8GMK1 | K7FZ14 | F1NQ64 |
| 145 | GS RKR | | | | | | | | | |
| 146 | KRKK | 125 | Q13352 | Centromere protein R | Transcription coregulator that can have both coactivator and corepressor functions | X1WGM0 | - | A0A1L8GLX9 | - | F1NP14 |
| 147 | RKKRK GK | 126 | Q14118 | Dystroglycan | The dystroglycan complex is involved in a number of processes including laminin and basement membrane assembly, sarcolemmal stability, cell survival, peripheral nerve myelination, nodal structure, cell migration, and epithelial polarization | Q8JHU7 | C3YV90 | A0A1L8GP44 | K7GFA6 | A4VAR9 |
| 148 | KKKGK | 127 | O00757 | Fructose-1,6-bisphosphatase isozyme 2 | Catalyzes the hydrolysis of fructose 1,6-bisphosphate to fructose 6-phosphate in the presence of divalent cations and probably participates in glycogen synthesis from carbohydrate precursors, such as lactate. | Q68EJ6 | - | Q7SZA0 | K7F7B3 | F1NYI2 |
| 149 | R SAGCRKGKHK RKK | 128 | P56645 | Period circadian protein homolog 3 | Kinase binding. Ubiquitin protein ligase binding | Q9I8L4 | - | A0A1L8FGK4 | - | E1C8E2 |
| 150 | RRKR | 129 | Q14934 | Nuclear factor of activated T-cells, cytoplasmic 4 | Plays a role in the inducible expression of cytokine genes in T-cells, especially in the induction of the IL-2 | E7F059 | C3YFD8 | A0A1L8HZL8 | K7FYL4 | A0A1D5NT38 |
| 151 | KRR Y | | | | | | | | | |
| 152 | PPVKRERTS | 130 | Q9H6Z4 | Ran-binding protein 3 | Acts as a cofactor for XPO1/CRM1-mediated nuclear export, perhaps as export complex scaffolding protein | Q6PCR4 | C3ZFL9 | Q6IP83 | - | F1NJ42 |
| 153 | PQSRKKLR | 131 | P61244 | Protein max | Transcription regulator. Forms a sequence-specific DNA-binding protein complex with MYC or MAD which recognizes the core sequence 5'-CAC[GA]TG-3'. | Q6NY29 | C3XRZ9 | Q07016 | K7FRK5 | P52162 |
| 154 | KVTKRKHDNEGSGSKRPK | 132 | P12956 | X-ray repair cross-complementing protein 6 | Single-stranded DNA-dependent ATP-dependent helicase. Has a role in chromosome translocation. | F1QZ60 | C3Z1F8 | Q6AZR6 | K7GE42 | O93257 |
| 155 | RAIKRRPGLDFDDDGEGNSKFLR | 133 | P27540 | Aryl hydrocarbon receptor nuclear translocator | Required for activity of the Ah (dioxin) receptor. This protein is required for the ligand-binding subunit to translocate from the cytosol to the nucleus after ligand binding. | Q9DG12 | C3Y283 | A0A1L8H0U4 | K7G3H3 | E1C1C5 |
| 156 | RVHPYQR | 134 | Q96PU8 | Protein quaking | RNA-binding protein that plays a central role in myelinization | Q6P104 | C3XVZ1 | A0A1L8G7T1 | - | A0A1D5P2B1 |
| 157 | KKQTTLAFKPIKKGKKR | 135 | P11388 | DNA topoisomerase 2-alpha | Control of topological states of DNA by transient breakage and subsequent rejoining of DNA strands. Topoisomerase II makes double-strand breaks. Essential during mitosis and meiosis for proper segregation of daughter chromosomes. May play a role in regulating the period length of ARNTL/BMAL1 transcriptional oscillation | Q1LUT2 | C3YYJ6 | A0A1L8FW96 | K7FJR3 | E1BZ19 |
| 158 | KKEKKKSKK | 136 | O60832 | H/ACA ribonucleoprotein complex subunit DKC1 | Catalytic subunit of H/ACA small nucleolar ribonucleoprotein (H/ACA snoRNP) complex, which catalyzes pseudouridylation of rRNA | F1Q749 | C3Y7R6 | - | - | Q5ZJH9 |
| 159 | KKx{15}KKRK | 137 | P46063 | ATP-dependent DNA helicase Q1 | DNA helicase that may play a role in the repair of DNA that is damaged by ultraviolet light or other mutagens. Exhibits a magnesium-dependent ATP-dependent DNA-helicase activity that unwinds single- and double-stranded DNA in a 3'-5' direction | Q498Z7 | C3YKR1 | Q32NN1 | K7F4Y6 | Q8AYS4 |

| | | | | | | | | | | |
|---|---|---|---|---|---|---|---|---|---|---|
| 160 | SGYG KVSRRGGHQN SYKPY | 138 | Q14103 | Heterogeneous nuclear ribonucleoprotein D0 | Binds with high affinity to RNA molecules that contain AU-rich elements (AREs) found within the 3'-UTR of many proto-oncogenes and cytokine mRNAs. | Q7SXN2 | C3ZWI1 | Q7ZYE9 | K7FUP0 | F6QGM9 |
| 161 | QR KVKPQPPLSD AYLSGMPAKR RKT | 139 | P46379 | Large proline-rich protein BAG6 | ATP-independent molecular chaperone preventing the aggregation of misfolded and hydrophobic patches-containing proteins. | A3KPW9 | - | Q6PA26 | - | - |
| 162 | KRKMDETDA SSAVKVKR | 140 | Q13148 | TAR DNA-binding protein 43 | DNA and RNA-binding protein which regulates transcription and splicing. | Q802C7 | C3ZIE9 | A0A1L8FFB1 | K7FJ67 | Q5ZLN5 |
| 163 | RGGRGG GDRGGFGPGK MDSRGEHRQD RRERPY | 141 | P35637 | RNA-binding protein FUS | Binds both single-stranded and double-stranded DNA and promotes ATP-independent annealing of complementary single-stranded DNAs and D-loop formation in superhelical double-stranded DNA. May play a role in maintenance of genomic integrity. | F1R0M4 | C3Z8P8 | Q7ZXQ2 | - | Q6J4Y8 |
| 164 | KKPK | 142 | P05230 | Fibroblast growth factor 1 | Plays an important role in the regulation of cell survival, cell division, angiogenesis, cell differentiation and cell migration | Q5TLE2 | - | A0A1L8HW95 | K7FNQ6 | P48801 |
| 165 | PSKRQRV | 143 | P07910 | Heterogeneous nuclear ribonucleoproteins C1/C2 | Binds pre-mRNA and nucleates the assembly of 40S hnRNP particles | Q802Y4 | C3Z5T3 | A0A1L8I0B0 | K7FZZ4 | - |
| 166 | KRKRPR | 144 | O60682 | Musculin | Transcription repressor capable of inhibiting the transactivation capability of TCF3/E47. May play a role in regulating antigen-dependent B-cell differentiation. | Q32PV5 | C3Y214 | A0A1L8G8Q2 | K7F6Z1 | A0A1D5PSY8 |
| 167 | KKKKKK | 145 | Q96GQ7 | Probable ATP-dependent RNA helicase DDX27 | Probable ATP-dependent RNA helicase. Component of the nucleolar ribosomal RNA (rRNA) processing machinery that regulates 3' end formation of ribosomal 47S rRNA | Q6DRN0 | C3ZBT1 | Q4V836 | - | Q5ZLD0 |
| 168 | RKRRR | 146 | Q9H8V3 | Protein ECT2 | Guanine nucleotide exchange factor (GEF) that catalyzes the exchange of GDP for GTP | A0A0R4IKD5 | C3YJI7 | Q5XGV4 | K7FLF0 | E1BWT7 |
| 169 | PRKRP | | | | | | | | | |
| 170 | PSQQKRK | 147 | Q92547 | DNA topoisomerase 2-binding protein 1 | Required for DNA replication. Plays a role in the rescue of stalled replication forks and checkpoint control | F1R135 | C3Z1H4 | A0A1L8FWJ6 | K7FLF0 | A0A1D5P3M9 |
| 171 | KRPR | | | | | | | | | |
| 172 | RKRARAEKKA LKKKKKI | 148 | Q15020 | Squamous cell carcinoma antigen recognized by T-cells 3 | U6 snRNP-binding protein that functions as a recycling factor of the splicing machinery. Promotes the initial reassembly of U4 and U6 snRNPs following their ejection from the spliceosome during its maturation | Q6ZM52 | C3Y679 | A0A1L8HZW2 | - | A0A1D5P378 |
| 173 | SGKRGGRKRR SSRRSAGGGA GPGGAAGGGV GGGDEPGSPA QGKRGKKSAG C | 149 | Q15672 | Twist-related protein 1 | Acts as a transcriptional regulator. | Q9PTE3 | C3YC05 | A0A1L8FR58 | K7G204 | Q9PVG4 |
| 174 | LKKD KSRKRSYSPD GKESPSDKKS KTD | 150 | P43243 | Matrin-3 | May play a role in transcription or may interact with other nuclear matrix proteins to form the internal fibrogranular network | Q800H6 | - | - | K7G839 | Q8UWC5 |
| 175 | KKKLKKVD | | | | | | | | | |
| 176 | RRGRK RKAEKQ | 151 | O75475 | PC4 and SFRS1-interacting protein | Transcriptional coactivator involved in neuroepithelial stem cell differentiation and neurogenesis. | F8W309 | C3Z0T2 | Q5XXA8 | K7GK32 | R4GK64 |
| 177 | NN GEQIIRRRTR KRLNPEAL | 152 | Q9UHF7-2 | Zinc finger transcription factor Trps1 | Transcriptional repressor. Binds specifically to GATA sequences and represses expression of GATA-regulated genes at selected sites and stages in vertebrate development. | D4P0Z5 | - | A0A1L8FZU9 | K7G5X8 | A0A1D5PND7 |
| 178 | MGLRTAKKRG LGGGGKWKRE | 153 | P28329 | Choline O-acetyltransferase | Catalyzes the reversible synthesis of acetylcholine (ACh) from acetyl CoA and choline at cholinergic synapses | Q642H6 | C3ZX27 | A0A1L8GMF7 | K7GIT1 | Q90YJ9 |
| 179 | KRAADE VLAEAKKPR | 154 | Q6P1J9 | Parafibromin | Tumor suppressor probably involved in transcriptional and post-transcriptional control pathways | X1WHN7 | C3ZG24 | A0A1L8GND2 | - | Q5ZLM0 |
| 180 | LKRAGSPEVQ GAMGSPAPKR | 155 | O43593 | Lysine-specific demethylase hairless | Histone demethylase that specifically demethylates both mono- and dimethylated 'Lys-9' of histone H3. May act as a transcription regulator controlling hair biology (via targeting of collagens), neural activity, and cell cycle | UPI0000F2058B | - | - | - | - |
| 181 | YRK CLQAGMNLEA RKTKKKIKGI QQAT | 156 | P04150 | Glucocorticoid receptor | Has a dual mode of action: as a transcription factor that binds to glucocorticoid response elements (GRE), both for nuclear and mitochondrial DNA, and as a modulator of other transcription factors | Q1XHK0 | - | A0A1L8GV81 | K7GAE8 | Q38HX7 |
| 182 | KTR REKRAR | 157 | Q9Y266 | Nuclear migration protein nudC | Necessary for correct formation of mitotic spindles and chromosome separation during mitosis. Necessary for cytokinesis and cell proliferation | Q7ZVD2 | C3ZLJ1 | Q6IRP6 | - | Q5ZIN1 |

| | | | | | | | | | | |
|---|---|---|---|---|---|---|---|---|---|---|
| 183 | PVAQKRK | 158 | P41218 | Myeloid cell nuclear differentiation antigen | May act as a transcriptional activator/repressor in the myeloid lineage. Plays a role in the granulocyte/monocyte cell-specific response to interferon. Stimulates the DNA binding of the transcriptional repressor protein YY1 | UPI00038371A7 | - | - | - | - |
| 184 | RRAAPT TPPPPVKRRD RE | 159 | Q9NZQ3 | NCK-interacting protein with SH3 domain | Has an important role in stress fiber formation induced by active diaphanous protein homolog 1 (DRF1) | UPI0007EEE691 | C3Z2V1 | A0A1L8GHQ7 | - | E1C7M3 |
| 185 | KKKSRK | 160 | Q01954 | Zinc finger protein basonuclin-1 | Transcriptional activator | A0A0G2KIM8 | C3ZGK9 | A0A1L8GST0 | K7FIP4 | E1C4G7 |
| 186 | RKRKKRKKGE KQIPGEEKGR KRRRVKEDKK KRDRDR | 161 | Q9NPI1 | Bromodomain-containing protein 7 | Acts both as coactivator and as corepressor | B0S7E1 | C3YFI8 | Q802C4 | - | Q5ZKG2 |
| 187 | KG VKRKADTTTP | 162 | Q58F21 | Bromodomain testis-specific protein | Testis-specific chromatin protein that specifically binds histone H4 acetylated at 'Lys-5' and 'Lys-8' (H4K5ac and H4K8ac, respectively) and plays a key role in spermatogenesis | F1QQP6 | C3ZPB3 | A1L1G8 | - | E1C8U8 |
| 188 | K RRIRRRPR | 163 | P41002 | | | Q6NYX6 | C3Z7N7 | Q7T0L6 | K7G0H4 | A0A1D5PBK0 |
| 189 | RRT KRKR | | | | | | | | | |
| 190 | KLKRR | 164 | Q49MI3 | Ceramide kinase-like protein | Has no detectable ceramide-kinase activity. Overexpression of CERKL protects cells from apoptosis in oxidative stress conditions | A5WVZ0 | C3YV82 | Q6GLV1 | K7G4B6 | Q5F3H3 |
| 191 | RRTTRRHP NTQQRASKKK PK | 165 | Q15021 | Condensin complex subunit 1 | Regulatory subunit of the condensin complex, a complex required for conversion of interphase chromatin into mitotic-like condense chromosomes. | B0V123 | C3XPS2 | Q9YHY6 | K7GDZ8 | Q5F475 |
| 192 | RKRL KSK | 166 | Q13309 | S-phase kinase-associated protein 2 | Substrate recognition component of a SCF (SKP1-CUL1-F-box protein) E3 ubiquitin-protein ligase complex which mediates the ubiquitination and subsequent proteasomal degradation of target proteins involved in cell cycle progression, signal transduction and transcription | A3KP53 | C3YM20 | Q075B2 | - | F7BEK8 |
| 193 | KKSKRGR | 167 | Q9UBS0 | Ribosomal protein S6 kinase beta-2 | Phosphorylates specifically ribosomal protein S6. | Q802Y7 | C3YDD4 | Q9W6Y9 | K7FKX2 | Q5ZLW5 |
| 194 | KKKMKKH KNKSEAKKRK IEFLRPGYKA KRQK | 168 | O75691 | Small subunit processome component 20 homolog | Involved in 18S pre-rRNA processing. Associates with U3 snoRNA. | A0A0R4IQW7 | C3ZQ27 | - | K7GES8 | E1BVC5 |
| 195 | KVLGVNVMLR K | 169 | P29373 | Cellular retinoic acid-binding protein 2 | Transports retinoic acid to the nucleus. Regulates the access of retinoic acid to the nuclear retinoic acid receptors. | A4VCG2 | C3Y8B2 | A0A1L8FCB0 | K7FJ80 | R4GL88 |
| 196 | G KMDKGEHRQE RRDRPY | 170 | Q01844 | RNA-binding protein EWS | Might normally function as a transcriptional repressor | Q4QRG0 | C3Z8P8 | Q2VPG6 | - | A0A1D5NUX9 |
| 197 | RKRR | 171 | Q14493 | Histone RNA hairpin-binding protein | RNA-binding protein involved in the histone pre-mRNA processing | A1L2F4 | C3Y0M4 | A0A1L8HT76 | - | Q6B7Z7 |
| 198 | RYKRK | | | | | | | | | |
| 199 | KR GLNSSFETSP KKVKW | 172 | Q8IV63 | Inactive serine/threonine-protein kinase VRK3 | Inactive kinase that suppresses ERK activity by promoting phosphatase activity of DUSP3 which specifically dephosphorylates and inactivates ERK in the nucleus | Q5BLE9 | C3XZ03 | A0A1L8GK79 | K7G2S1 | F1NDK1 |
| 200 | RRKSI KKKR | 173 | Q15303 | Receptor tyrosine-protein kinase erbB-4 | Tyrosine-protein kinase that plays an essential role as cell surface receptor for neuregulins and EGF family members and regulates development of the heart, the central nervous system and the mammary gland, gene transcription, cell proliferation, differentiation, migration and apoptosis | A0A0A0MPK4 | C3YZ4 | A0A1L8FRS8 | - | A0A1D5PKH0 |
| 201 | STKKRER | 174 | O60934 | Nibrin | Component of the MRE11-RAD50-NBN (MRN complex) which plays a critical role in the cellular response to DNA damage and the maintenance of chromosome integrity | Q5I2W8 | C3YFG1 | Q6XV80 | - | A0A1D5P1G5 |
| 202 | K RKR | 175 | Q96C86 | m7GpppX diphosphatase | Decapping scavenger enzyme that catalyzes the cleavage of a residual cap structure following the degradation of mRNAs by the 3'->5' exosome-mediated mRNA decay pathway | F1R776 | C3ZTI1 | Q4KLD2 | - | - |
| 203 | RKRKPS | 176 | Q7Z5L9 | Interferon regulatory factor 2-binding protein 2 | Acts as a transcriptional corepressor in a IRF2-dependent manner; this repression is not mediated by histone deacetylase activities | Q6NZT6 | - | A0A1L8G830 | K7EX81 | A0A1D5PGK9 |
| 204 | SDPKGPPAKI ARLEQNGSPL | 177 | Q01826 | DNA-binding protein SATB1 | Binds to DNA at special AT-rich sequences, the consensus SATB1-binding sequence (CSBS), at nuclear matrix- or scaffold-associated regions. | E1ANH4 | - | A0A1L8FW95 | K7GJC9 | E1BZL3 |

| | | | | | | | | | |
|---|---|---|---|---|---|---|---|---|---|
| 205 | KRQRKRR | 178 | P10244 | Myb-related protein B | Transcription factor involved in the regulation of cell survival, proliferation, and differentiation. Transactivates the expression of the CLU gene. | Q4KMF7 | C3ZI75 | Q08759 | K7GDA9 | Q03237 |
| 206 | RRR | 179 | Q12778 | Forkhead box protein O1 | Transcription factor that is the main target of insulin signaling and regulates metabolic homeostasis in response to oxidative stress | A3RK75 | C3YI59 | A0A1L8HIX1 | K7EZG6 | F1NNE8 |
| 207 | RKRTPRVDGQT GENDMNKRRR K | 180 | O94851 | [F-actin]-monooxygenase MICAL2 | Nuclear monooxygenase that promotes depolymerization of F-actin by mediating oxidation of specific methionine residues on actin to form methionine-sulfoxide, resulting in actin filament disassembly and preventing repolymerization | A0A0A0MPT7 | C3Z6G9 | - | - | Q5F3F3 |
| 208 | KKRKK | 181 | Q969P5 | F-box only protein 32 | Substrate recognition component of a SCF (SKP1-CUL1-F-box protein) E3 ubiquitin-protein ligase complex which mediates the ubiquitination and subsequent proteasomal degradation of target proteins. Probably recognizes and binds to phosphorylated target proteins during skeletal muscle atrophy. Recognizes TERF1. | Q7ZTY2 | C3Y232 | A0A1L8FTW6 | K7F351 | Q5ZHM6 |
| 209 | K AFFQSEDSPS PKRQRLSHSV F | 182 | Q9H0F5 | E3 ubiquitin-protein ligase RNF38 | Acts as an E3 ubiquitin-protein ligase able to ubiquitinate p53/TP53 which promotes its relocalization to discrete foci associated with PML | E7FDX7 | - | A0A1L8GY19 | K7GE35 | F1NND0 |
| 210 | RNRRSPP VRRQRGRRD R | | | | | | | | | |
| 211 | RKKNKY RPISHTPS | 183 | Q96K30 | RBPJ-interacting and tubulin-associated protein 1 | Tubulin-binding protein that acts as a negative regulator of Notch signaling pathway. Shuttles between the cytoplasm and the nucleus and mediates the nuclear export of RBPJ/RBPSUH, thereby preventing the interaction between RBPJ/RBPSUH and NICD product of Notch proteins (Notch intracellular domain), leading to down-regulate Notch-mediated transcription. May play a role in neurogenesis. | Q1JPZ6 | C3Y4L6 | A0A1L8I064 | - | - |
| 212 | K KRGKKKKRGR | 184 | Q9HBH9 | MAP kinase-interacting serine/threonine-protein kinase 2 | Serine/threonine-protein kinase that phosphorylates SFPQ/PSF, HNRNPA1 and EIF4E | Q7ZVV9 | C3YM17 | A0A1L8HWS4 | K7F5G5 | F1NCK3 |
| 213 | RKRKK | 185 | Q16666 | Gamma-interferon-inducible protein 16 | Binds double-stranded DNA. Binds preferentially to supercoiled DNA and cruciform DNA structures | UPI000383722A | - | - | - | - |
| 214 | KRK K | | | | | | | | | |
| 215 | KEK | | | | | | | | | |
| 216 | K GSK | | | | | | | | | |
| 217 | RKRGRGRPKG RGRGRPRGRP RGSKR | 186 | Q9UKY1 | Zinc fingers and homeoboxes protein 1 | Acts as a transcriptional repressor. Increases DNMT3B-mediated repressive transcriptional activity when DNMT3B is tethered to DNA. May link molecule between DNMT3B and other co-repressor proteins | A5D8R5 | C3Z1B2 | A0A1L8G021 | K7FK68 | A0A1D5PKX3 |
| 218 | RKA KRAR | 187 | Q86WB0 | Nuclear-interacting partner of ALK | Essential component of a SCF-type E3 ligase complex, SCF(NIPA), a complex that controls mitotic entry by mediating ubiquitination and subsequent degradation of cyclin B1 (CCNB1) | Q08BA2 | - | A0A1L8GTM4 | - | E1C3I1 |
| 219 | RKK | 188 | Q9UNZ2 | NSFL1 cofactor p47 | Together with UBXN2B/p37, regulates the centrosomal levels of kinase AURKA/Aurora A during | Q5U3U1 | C3Y288 | Q7ZYT4 | K7G8S4 | A0A1L1RX58 |
| 220 | KRQH | | | | | | | | | |
| 221 | KPNKRK | 189 | P51858 | Hepatoma-derived growth factor | Heparin-binding protein, with mitogenic activity for fibroblasts. Acts as a transcriptional repressor | Q5XXA7 | C3Z0T2 | A0A1L8HW52 | K7FIL0 | A0A1D5PAE4 |
| 222 | KRRAGD LLEDSPKRPK | | | | | | | | | |
| 223 | KRQEME SGITTPPKMR R | 190 | P21359 | Neurofibromin | Stimulates the GTPase activity of Ras. NF1 shows greater affinity for Ras GAP, but lower specific activity. May be a regulator of Ras activity. | E7FFY1 | C3ZV81 | A0A1L8HGF2 | - | A0A1D5P1U6 |
| 224 | KK KGRTQKEKKA ARARSKGK | 191 | Q9UNL4 | Inhibitor of growth protein 4 | Component of the HBO1 complex which has a histone H4-specific acetyltransferase activity, a reduced activity toward histone H3 and is responsible for the bulk of histone H4 acetylation in vivo. | Q4VBS0 | C3ZSF3 | A0A1L8FDZ0 | K7FSP4 | Q5ZKY4 |
| 225 | KRRRRR RREKRKR | 192 | Q96GM8 | Target of EGR1 protein 1 | Inhibits cell growth rate and cell cycle. Induces CDKN1A expression as well as TGF-beta expression. Mediates the inhibitory growth effect of EGR1. Involved in the maturation of snRNAs and snRNA 3'-tail processing | B8A5U4 | C3YIK6 | A0A1L8GFN0 | - | F1N9P8 |
| 226 | GKN EAKKRK | 193 | P52739 | Zinc finger protein 131 | Plays a role during development and organogenesis as well as in the function of the adult central nervous | Q7SXM6 | - | A0A1L8I2R2 | - | E1C6B5 |
| 227 | KK QRTGKKIH | | | | | | | | | |

| | | | | | | | | | | |
|---|---|---|---|---|---|---|---|---|---|---|
| 228 | KG RRGRLPSKPK | 194 | P43354 | Nuclear receptor subfamily 4 group A member 2 | Transcriptional regulator which is important for the differentiation and maintenance of meso-diencephalic dopaminergic (mdDA) neurons during development. It is crucial for expression of a set of genes such as SLC6A3, SLC18A2, TH and DRD2 which are essential for development of mdDA neurons | Q6GMG3 | C3YEV5 | Q7T0V3 | K7FLA5 | A0A1D5NYR5 |
| 229 | KKKP AK | 195 | Q9BRS8 | La-related protein 6 | Regulates the coordinated translation of type I collagen alpha-1 and alpha-2 mRNAs, CO1A1 and CO1A2. Stabilizes mRNAs through high-affinity binding of a stem-loop structure in their 5' UTR. This regulation requires VIM and MYH10 filaments, and the helicase DHX9 | Q6PHD4 | C3ZKS9 | A0A1L8H0A5 | - | F1NWE6 |
| 230 | RR R | 196 | Q9HC62 | Sentrin-specific protease 2 | Protease that catalyzes two essential functions in the SUMO pathway. The first is the hydrolysis of an alpha- [illegible] | E7F9K9 | C3XYA3 | Q804S7 | K7G2N0 | A0A1D5PCU8 |
| 231 | PAKRP R | | | | | | | | | |
| 232 | KRRH | 197 | Q12968 | Nuclear factor of activated T-cells, cytoplasmic 3 | Acts as a regulator of transcriptional activation. Plays a role in the inducible expression of cytokine genes in T- [illegible] | E7FE12 | C3YFD8 | A0A1L8GK58 | K7FYL4 | A0A1D5P8S9 |
| 233 | KRKK | | | | | | | | | |
| 234 | KRK | 198 | O43148 | mRNA cap guanine-N7 methyltransferase | Catalytic subunit of the mRNA-capping methyltransferase RNMT:RAMMET complex that methylates the N7 position of the added guanosine to the 5'-cap structure of mRNAs | Q1MTD3 | C3ZV89 | A0A1L8FYD5 | K7FIZ9 | E1BYM1 |

***In silico* predicted NLSs**

| | | | | | | | | | | |
|---|---|---|---|---|---|---|---|---|---|---|
| 1 | RKKRR | 1 | Q9UER7 | Death domain-associated protein 6 | Transcription corepressor known to repress transcriptional potential of several sumoylated [illegible] | Q567Z8 | - | Q2KHQ3 | - | - |
| 2 | PCK KSRK | | | | | | | | | |
| 3 | PKPS | 2 | P54253 | Ataxin-1 | Chromatin-binding factor that repress Notch signaling in the absence of Notch intracellular domain by acting as a CBF1 corepressor. | Q1L966 | - | A0A1L8FSD1 | - | E1BZ23 |
| 4 | VKRRRM | 3 | Q03001 | Dystonin | Cytoskeletal linker protein. Acts as an integrator of intermediate filaments, actin and microtubule cytoskeleton networks. | A0A0G2KQI8 | C3YBC4 | - | - | A0A1D5P0Y6 |
| 5 | K KELPSHVQAM HKRKEFL | 4 | Q7Z333 | Probable helicase senataxin | Probable RNA/DNA helicase involved in diverse aspects of RNA metabolism and genomic integrity. | E7FAV9 | C3ZM47 | A0A1L8HDV6 | - | Q5ZKD7 |
| 6 | RARLA NTQGKKAKRK AREKQLEEAR RLAALQKRRE LRAAGIEIQK KRKRKRGVDY NAEIPFEKKP ALGFYDTSEE NYQALDADFR KLRQQDLDGE LRSEKEGRDR KKDKQHLKRK K | 5 | Q99459 | Cell division cycle 5-like protein | DNA-binding protein involved in cell cycle control. May act as a transcription activator. | E9QIC1 | C3YHB1 | A1A5J6 | K7G2Y2 | E1C6A9 |
| 7 | KKLEV NEAELLRRRN FK | 6 | Q6NZI2 | Caveolae-associated protein 1 | Plays an important role in caveolae formation and organization. Essential for the formation of caveolae in [illegible] | E7F0K3 | - | A0A1L8EKH0 | K7GED2 | Q90885 |
| 8 | KKAFSKEK MEKTKVRTR | | | | | | | | | |
| 9 | RKKRR | 7 | Q0VD86 | Protein INCA1 | cyclin-dependent protein serine/threonine kinase inhibitor activity | E9QDB1 | - | - | - | - |
| 10 | QVGA GGEPLDACYL KALEGFVMVL | 8 | Q9Y2N7 | Hypoxia-inducible factor 3-alpha | Acts as a transcriptional regulator in adaptive response to low oxygen tension. | Q6EHI4 | - | Q6PI54 | K7FUG2 | Q9YIB9 |
| 11 | LQA NGPVAKKAKK KA | 9 | Q14978 | Nucleolar and coiled-body phosphoprotein 1 | Nucleolar protein that acts as a regulator of RNA polymerase I by connecting RNA polymerase I with enzymes responsible for ribosomal processing and modification | B8JIY2 | - | Q91803 | K7GC40 | A0A1D5PUU2 |
| 12 | TKNSSNK PAVTTKSPAV KPAAAPKQPV GGGQKLLTRK ADSSSSEEES SSSEEEKTKK MVATTKPKAT AKAALSLPAK QAPQGSRDSS SDSDSSSSEE EEEKTSKSAV KKKPQKVAGG AAPSKPASAK KGKAESSNSS SSDDSSEEEE EKLKGKGSPR PQAPKANGTS ALTAQNGKAA KNSEEEEEEK KKAAVVVSKS GSLKKRK | | | | | | | | | |
| 13 | KIKLQTPNT FPKRKKG | | | | | | | | | |
| 14 | LSPSL SPL | 10 | Q9Y261 | Hepatocyte nuclear factor 3-beta | Transcription factor that is involved in embryonic development, establishment of tissue-specific gene expression and regulation of gene expression in differentiated tissues. | Q6P0C9 | C3XR49 | A0A1L8FAN2 | K7FXK1 | Q9PWP8 |

| | | | | | | | | | |
|---|---|---|---|---|---|---|---|---|---|
| 15 | GP RAKVAKLNIQ SLSPVKKKKM VPGALGVPAD LAPVDVEFSF PKFSRLRRGL KAEAVKGPVP AAPARRRLQL PRLRVR | 11 | Q9BXM0 | Periaxin | Scaffolding protein that functions as part of a dystroglycan complex in Schwann cells, and as part of EZR and AHNAK-containing complexes in eye lens fiber cells. | F1QQL4 | C3YGB2 | A0A1L8G1U2 | - | - |
| 16 | KPRK | 12 | Q8TDY2 | RB1-inducible coiled-coil protein 1 | Involved in autophagy. Regulates early events but also late events of autophagosome formation through direct interaction with Atg16L1 | E7FFM2 | - | A0A1L8FYZ6 | K7G9N3 | E1C330 |
| 17 | RKRKG | 13 | Q8WYA1 | Aryl hydrocarbon receptor nuclear translocator-like protein 2 | Transcriptional activator which forms a core component of the circadian clock. | Q9I879 | C3Y283 | Q6P6Z6 | K7FUW5 | E1C1C5 |
| 18 | KQEDKN AMDVIVKVIK R | 14 | Q13609 | Deoxyribonuclease gamma | Has DNA hydrolytic activity. Is capable of both single- and double-stranded DNA cleavage, producing DNA [illegible] | Q6DH05 | C3Z4K2 | A0A1L8EZN6 | K7F8E0 | Q3Y8N1 |
| 19 | LRKKT KSKR | | | | | | | | | |
| 20 | RSKPKK KSKKHKDKE | 15 | P55199 | RNA polymerase II elongation factor ELL | Elongation factor component of the super elongation complex (SEC), a complex required to increase the catalytic rate of RNA polymerase II transcription by suppressing transient pausing by the polymerase at multiple sites along the DNA. | Q6PEG4 | C3YQ98 | A0A1L8HX53 | K7GHT8 | F1NBA3 |
| 21 | KLVPGLFKDE MKRRR | 16 | P35227 | Polycomb group RING finger protein 2 | Transcriptional repressor. Binds specifically to the DNA sequence 5'-GACTNGACT-3'. | B0UXV8 | C3YW14 | Q6NU75 | K7FHU3 | F1NWQ4 |
| 22 | KEDSKI LKKARKDGFL HETLLDRRAK L | 17 | Q14331 | Protein FRG1 | Binds to mRNA in a sequence-independent manner. | Q566W5 | C3XXG6 | A0A1L8HU14 | - | E1BY86 |
| 23 | KKKSKDKK RKR | | | | | | | | | |
| 24 | RKAEHW KHKKEESGRE RKKNW | 18 | Q96AQ6 | Pre-B-cell leukemia transcription factor-interacting protein 1 | Regulator of pre-B-cell leukemia transcription factors (BPXs) function. Inhibits the binding of PBX1-HOX complex to DNA and blocks the transcriptional activity of E2A-PBX1. | A5WW29 | - | A0A1L8FDD5 | - | A0A1D5P2Y6 |
| 25 | KALKK RSGKKDKHSQ SPRAAGPREG | | | | | | | | | |
| 26 | RRTDPECT APIKKQKKR | 19 | Q8N1G2 | Cap-specific mRNA (nucleoside-2'-O-)-methyltransferase 1 | S-adenosyl-L-methionine-dependent methyltransferase that mediates mRNA cap1 2'-O-ribose methylation to the 5'-cap structure of mRNAs. | Q803R5 | C3Y0P7 | Q6GQ76 | - | F1NAT2 |
| 27 | RKKIS NFT | 20 | Q9H079 | KATNB1-like protein 1 | Regulates microtubule-severing activity of KATNAL1 in a concentration-dependent manner in vitro. | Q5XJA1 | - | Q5EAV1 | K7FQB2 | Q5ZIG6 |
| 28 | RKKVHHPF | | | | | | | | | |
| 29 | RDGFRRK | 21 | Q8NEY8 | Periphilin-1 | Component of the HUSH complex, a multiprotein complex that mediates epigenetic repression. | A1A5V1 | - | Q6DJM5 | - | A0A1D5NZJ2 |
| 30 | KLVTKL QNSERKKRGA RR | 22 | O60356 | Nuclear protein 1 | Chromatin-binding protein that converts stress signals into a program of gene expression that empowers cells with resistance to the stress induced by a change in their microenvironment. | E9QCY4 | - | A0A1L8H8H5 | - | - |
| 31 | IKKKQKH | 23 | P55209 | Nucleosome assembly protein 1-like 1 | May be involved in modulating chromatin formation and contribute to regulation of cell proliferation. | A0A0R4IDK6 | C3Y698 | A0A1L8GYL4 | K7FK99 | E1BZS2 |
| 32 | KKRKGR | 24 | Q9H147 | Deoxynucleotidyltransferase terminal-interacting protein 1 | Increases DNTT terminal deoxynucleotidyltransferase activity (in vitro) | F1Q8X8 | C3YCS8 | Q6IP16 | K7GBG4 | - |
| 33 | KKQR | 25 | Q9UFW8 | CGG triplet repeat-binding protein 1 | Binds to nonmethylated 5'-d(CGG)(n)-3' trinucleotide repeats in the FMR1 promoter. May play a role in regulating FMR1 promoter. | - | - | - | - | A0A1D5NUG1 |
| 34 | KRRKKE EKR | 26 | Q9H6E4 | Coiled-coil domain-containing protein 134 | In extracellular secreted form, promotes proliferation and activation of CD8+ T cells, suggesting a cytokine-like function | A0A0G2KTT4 | C3YZU0 | A0A1L8GNP2 | K7GB64 | E1BVM7 |
| 35 | RKRR | 27 | Q96EU6 | Ribosomal RNA processing protein 36 homolog | Involved in the early processing steps of the pre-rRNA in the maturation pathway leading to the 18S rRNA. | Q1L876 | C3ZKP2 | A0A1L8G0R1 | - | R4GFB9 |
| 36 | KKRRK | 28 | O95935 | T-box transcription factor TBX18 | Acts as transcriptional repressor involved in developmental processes of a variety of tissues and organs, including the heart and coronary vessels, the ureter and the vertebral column. Required for embryonic development of the sino atrial node (SAN) head area. | Q8JFF9 | C3XV96 | A0A1L8G3B1 | K7G3Q5 | Q68SB0 |
| 37 | KRRR | 29 | Q14153 | Protein FAM53B | Acts as a regulator of Wnt signaling pathway by regulating beta-catenin (CTNNB1) nuclear localization. | F1QN48 | - | - | - | E1C7N4 |
| 38 | APK KMKTS | 30 | Q9NS26 | Sperm protein associated with the nucleus on the X chromosome A | spermatogenesis | - | - | - | - | - |
| 39 | RKRYGYRL | 31 | O95478 | Ribosome biogenesis protein NSA2 homolog | Involved in the biogenesis of the 60S ribosomal subunit. | F1QDL2 | C3Y211 | K7G2C3 | Q6GP86 | Q5ZMH9 |
| 40 | RKSSHHRKAG ESHRRRAG | 32 | Q8TC56 | Protein FAM71B | May be involved in RNA biogenesis. | - | - | - | - | - |

| | | | | | | | | | | |
|---|---|---|---|---|---|---|---|---|---|---|
| 41 | RRRRAS APISQWSSSR RSR | 33 | Q8N699 | Myc target protein 1 | May regulate certain MYC target genes, MYC seems to be a direct upstream transcriptional activator. | E7FG94 | - | A0A1L8G7Z9 | K7FBA2 | - |
| 42 | RRRFK | 34 | P55786 | Puromycin-sensitive aminopeptidase | Aminopeptidase with broad substrate specificity for several peptides. Involved in proteolytic events essential for cell growth and viability. | E7F2N2 | C3YCZ6 | A0A1L8I1H6 | K7FNK2 | A0A1D5PAZ7 |
| 43 | RQKRQARESN SDRVSASKRR SSPSKDGR | 35 | P61328 | Fibroblast growth factor 12 | Involved in nervous system development and function. Involved in the positive regulation of voltage-gated sodium channel activity. Promotes neuronal excitability by elevating the voltage dependence of neuronal sodium channel SCN8A fast inactivation. | A0AUQ1 | - | A0A1L8F810 | K7FYZ7 | Q9IAI5 |
| 44 | KKKKEK | 36 | Q8NI27 | THO complex subunit 2 | Required for efficient export of polyadenylated RNA and spliced mRNA. | F1QXF4 | C3YRS5 | A0A1L8F2J0 | K7GAE7 | A0A1D5PRL3 |
| 45 | QQKKQ | 37 | Q76L83 | Putative Polycomb group protein ASXL2 | Putative Polycomb group (PcG) protein. PcG proteins act by forming multiprotein complexes, which are required to maintain the transcriptionally repressive state of homeotic genes throughout development. PcG proteins are not required to initiate repression, but to maintain it during later stages of development. | E7FG88 | - | - | - | E1BTW9 |
| 46 | RRKRRRRR | 38 | Q9BQK8 | Phosphatidate phosphatase LPIN3 | Regulates fatty acid metabolism. Magnesium-dependent phosphatidate phosphatase enzyme which catalyzes the conversion of phosphatidic acid to diacylglycerol during triglyceride, phosphatidylcholine and phosphatidylethanolamine biosynthesis | F1RAK9 | C3YA84 | A0A1L8FYL5 | K7FWE2 | A0A1D5PV53 |
| 47 | LKKRAR | 39 | P57772 | Selenocysteine-specific elongation factor | Translation factor necessary for the incorporation of selenocysteine into proteins. It probably replaces EF-Tu for the insertion of selenocysteine directed by the UGA codon. SelB binds GTP and GDP. | Q32LX1 | C3YLP3 | Q6GPS7 | - | A0A1D5PXD0 |
| 48 | KKRRRRARK | 40 | Q9UMS6 | Synaptopodin-2 | Has an actin-binding and actin-bundling activity. Can induce the formation of F-actin networks in an isoform-specific manner | E9QFY4 | C3Z5E7 | A0A1L8FK03 | K7FHA8 | F1NH40 |
| 49 | KRK R | 41 | P49918 | Cyclin-dependent kinase inhibitor 1C | Potent tight-binding inhibitor of several G1 cyclin/CDK complexes (cyclin E-CDK2, cyclin D2-CDK4, and cyclin A-CDK2) and, to lesser extent, of the mitotic cyclin B-CDC2. | Q6IQT7 | - | Q91603 | K7FV40 | Q8JIV2 |
| 50 | RA RARARARA | 42 | Q9UH62 | Armadillo repeat-containing X-linked protein 3 | Regulates mitochondrial aggregation and transport in axons in living neurons. | A0A0R4IWE3 | C3ZBB8 | A0A1L8GZ89 | K7FUU6 | E1C223 |
| 51 | KKYGPK | 43 | Q16527 | Cysteine and glycine-rich protein 2 | Drastically down-regulated in response to PDGF-BB or cell injury, that promote smooth muscle cell proliferation and dedifferentiation. Seems to play a role in the development of the embryonic vascular system. | Q6P2T6 | C3ZK45 | A0A1L8GUD0 | K7G1K8 | A0A1D5NYM7 |
| 52 | RRRRHRNR | 44 | P61579 | Endogenous retrovirus group K member 25 Rec protein | Retroviral replication requires the nuclear export and translation of unspliced, singly-spliced and multiply-spliced derivatives of the initial genomic transcript. | - | - | - | - | - |
| 53 | KRS RRKRRR | 45 | Q6NSI3 | Protein FAM53A | May play an important role in neural development; the dorsomedial roof of the third ventricle. | F1QN48 | - | A0A1L8HT13 | - | Q5ZKN5 |
| 54 | KRRKR | 46 | Q14693 | Phosphatidate phosphatase LPIN1 | Plays important roles in controlling the metabolism of fatty acids at different levels. | X1WH54 | C3YA84 | A0A1L8G5J3 | K7FWE2 | A0A1D5P4T3 |
| 55 | IKKKRPPV | 47 | Q8IVN3 | Musculoskeletal embryonic nuclear protein 1 | May be involved in the development and regeneration of the musculoskeletal system. | Q7ZVB7 | C3YAD5 | Q7T0M7 | K7GIR2 | A0A1D5PW18 |
| 56 | VRKKR KP | 48 | P49321 | Nuclear autoantigenic sperm protein | Required for DNA replication, normal cell cycle progression and cell proliferation. Forms a cytoplasmic complex with HSP90 and H1 linker histones and stimulates HSP90 ATPase activity. | F1QNE0 | C3YIL0 | - | Q2VPH0 | - |
| 57 | RQRRSQNKFLRRQRKPQRK | 49 | Q6NW34 | Nucleolus and neural progenitor protein | May play a role in cortex development as part of the Notch signaling pathway. | Q567G6 | C3XS37 | A0A1L8HH29 | K7F2R4 | - |
| 58 | KRLVPQ KQASVAKKKK | 50 | Q86SE8 | Nucleoplasmin-2 | Core histones chaperone involved in chromatin reprogramming, specially during fertilization and early embryonic development. Probably involved in sperm DNA decondensation during fertilization. | Q5BLA5 | - | Q6GQG6 | K7FKB9 | P16039 |
| 59 | RRQL VHFAWEHFRP RCK | 51 | Q9NZT2 | Opioid growth factor receptor | Receptor for opioid growth factor (OGF), also known as Met-enkephalin. Seems to be involved in growth regulation. | E7EYZ3 | C3ZI17 | A0A1L8G3Q4 | - | F1NEX4 |

| | | | | | | | | | |
|---|---|---|---|---|---|---|---|---|---|
| 60 | RRR SKFANLGRIF KPWKWRKKK | 52 | Q9C0D0 | Phosphatase and actin regulator 1 | Binds actin monomers (G actin) and plays a role in the reorganization of the actin cytoskeleton and in formation of actin stress fibers. Plays a role in cell motility. Plays a role in the formation of tubules by endothelial cells. Regulates PPP1CA activity. Required for normal cell survival. | F1R9D7 | - | A0A1L8FY03 | - | F1NWB1 |
| 61 | RRRKRR | 53 | Q6MZT1 | Regulator of G-protein signaling 7-binding protein | Regulator of G protein-coupled receptor (GPCR) signaling. Regulatory subunit of the R7-Gbeta5 complexes that acts by controlling the subcellular location of the R7-Gbeta5 complexes. | B2GRF5 | - | A0A1L8I255 | K7FKC4 | R4GI70 |
| 62 | KKIKKKKKK KHR | 54 | Q5VWQ0 | Lysine-specific demethylase 9 | Histone demethylase that specifically demethylates dimethylated 'Lys-20' of histone H4 (H4K20me2), thereby modulating chromosome architecture. | A0A0G2KVT3 | - | A0A1L8HEY8 | - | A0A1D5PEL9 |
| 63 | VARRR | 55 | Q7Z7C7 | Stimulated by retinoic acid gene 8 protein homolog | Meiosis-inducer required for the transition into meiosis for both female and male germ cells. In female germ cells, required for premeiotic DNA replication and subsequent events in meiotic prophase. | - | C3Y531 | A0A1L8GQI6 | - | E1C4S4 |
| 64 | KKKRK | 56 | O75683 | Surfeit locus protein 6 | Binds to both DNA and RNA in vitro, with a stronger binding capacity for RNA. May represent a nucleolar [illegible] | Q7T3A6 | - | Q0IH42 | - | Q5ZHN7 |
| 65 | RKKRK | | | | | | | | | |
| 66 | KLKKKKMA | 57 | Q05952 | Nuclear transition protein 2 | Plays a key role in the replacement of histones to protamine in the elongating spermatids of mammals. In condensing spermatids, loaded onto the nucleosomes, where it promotes the recruitment and processing of protamines, which are responsible for histone eviction. | XP_021324227.1 | - | - | - | - |
| 67 | KKWK | 58 | Q15431 | Synaptonemal complex protein 1 | Major component of the transverse filaments of synaptonemal complexes (SCS), formed between homologous chromosomes during meiotic prophase. [illegible] | F1QYS1 | - | - | - | - |
| 68 | LE EV | | | | | | | | | |
| 69 | KK KRK | | | | | | | | | |
| 70 | KRK R | 59 | Q8IZU3 | Synaptonemal complex protein 3 | Component of the synaptonemal complexes (SCS), formed between homologous chromosomes during meiotic prophase. Required for centromere pairing during meiosis in male germ cells | Q1RLP7 | C3ZM03 | A0A1L8GIK4 | - | E1BWM3 |
| 71 | RKRLAG | 60 | Q9P016 | Thymocyte nuclear protein 1 | Specifically binds 5-hydroxymethylcytosine (5hmC), suggesting that it acts as a specific reader of 5hmC. | F1QDP5 | C3XST2 | A0A1L8FL07 | K7GB28 | Q90679 |
| 72 | KYKEGK | 61 | O60688 | Protein yippee-like 1 | May play a role in epithelioid conversion of fibroblasts. | Q6AXK6 | C3YVP3 | Q6DDQ0 | K7F5P3 | F1NA62 |
| 73 | RAKWR RHQRA | 62 | P0C7M4 | Rhox homeobox family member 2B | Transcription factor maybe involved in reproductive processes. Modulates expression of target genes encoding proteins involved in processes relevant to spermatogenesis. | Q6P295 | - | - | - | - |
| 74 | VHFKISGE KRPSADPGKK AKNPKKKKKKDP | 63 | Q96NM4 | TOX high mobility group box family member 2 | Putative transcriptional activator involved in the hypothalamo-pituitary-gonadal system. | B0V356 | - | A0A1L8FZ33 | K7FWE5 | A0A1D5PSC7 |
| 75 | RPKGRQRK | 64 | O15234 | Protein CASC3 | Core component of the splicing-dependent multiprotein exon junction complex (EJC) deposited at splice [illegible] | Q1ECZ4 | C3XUF7 | A0JMU8 | - | A0A1D5PM89 |
| 76 | KPRRIRK PR | | | | | | | | | |
| 77 | A RVCRRK | 65 | Q7Z2W4 | Zinc finger CCCH-type antiviral protein 1 | Antiviral protein which inhibits the replication of viruses by recruiting the cellular RNA degradation machineries to degrade the viral mRNAs. | A0A0R4IH74 | C3YKR0 | A0A1L8G4F7 | K7FBX9 | A0A1D5PEY4 |
| 78 | KKR IRPLETTQQI RKRHC | 66 | Q9BX63 | Fanconi anemia group J protein | DNA-dependent ATPase and 5' to 3' DNA helicase required for the maintenance of chromosomal stability. | B7ZDD5 | C3YXX3 | A0A1L8HG08 | K7G641 | Q3YK19 |
| 79 | RKR RRR | 67 | Q8N693 | Homeobox protein ESX1 | May coordinately regulate cell cycle progression and transcription during spermatogenesis. | Q6P984 | - | - | - | - |
| 80 | KKRK | 68 | Q14320 | Protein FAM50A | May be a DNA-binding protein or transcriptional factor. | Q568K9 | C3ZVJ9 | Q2VPH1 | - | - |
| 81 | PKVKRSRKA | 69 | P41223 | Protein BUD31 homolog | Involved in the pre-mRNA splicing process | A0A0R4I9A3 | C3Y9A1 | A0A1L8EXA3 | K7FWH2 | E1BV47 |
| 82 | KKKR KR | 70 | Q7LFL8 | CXXC-type zinc finger protein 5 | May indirectly participate in activation of the NF-kappa-B and MAPK pathways. | E7FC24 | - | A0A1L8HV25 | - | F1NCW6 |
| 83 | KAKRVSRN KSEKKRR | 71 | O15516 | Circadian locomoter output cycles protein kaput | Transcriptional activator which forms a core component of the circadian clock. | B3DH92 | C3XWF6 | Q9I9Q6 | K7FXZ9 | Q8QGQ6 |
| 84 | KRKHV VK | 72 | Q8N9N8 | Probable RNA-binding protein EIF1AD | Plays a role into cellular response to oxidative stress. Decreases cell proliferation. | Q7SY07 | C3XYZ9 | A0A1L8GK34 | K7F4B4 | Q6K1L7 |
| 85 | KYRKN IWIKR | | | | | | | | | |
| 86 | KRKKRRHR | 73 | Q9NZR4 | Visual system homeobox 1 | Binds to the 37-bp core of the locus control region (LCR) of the red/green visual pigment gene cluster. May regulate the activity of the LCR and the cone opsin genes at earlier stages of development. | O42250 | C3XZK7 | Q0P031 | - | Q9IAL2 |

| | | | | | | | | | | |
|---|---|---|---|---|---|---|---|---|---|---|
| 87 | RRR | 74 | Q9BSI4 | TERF1-interacting nuclear factor 2 | Component of the shelterin complex (telosome) that is involved in the regulation of telomere length and protection. | F1R9M2 | - | - | - | - |
| 88 | KT KRK | 75 | Q8IX15 | Homeobox and leucine zipper protein Homez | May function as a transcriptional regulator. | A5D8R5 | C3Z1B2 | A0A1L8G021 | K7FK68 | A0A1D5PYW2 |
| 89 | RKRR K | 76 | Q00653 | Nuclear factor NF-kappa-B p100 subunit | NF-kappa-B is a pleiotropic transcription factor present in almost all cell types and is the endpoint of a series of signal transduction events that are initiated by a vast array of stimuli related to many biological processes such as inflammation, immunity, differentiation, cell growth, tumorigenesis and apoptosis. | Q0II00 | - | P51510 | K7FNJ5 | P98152 |
| 90 | R KKRGR | 77 | Q96JN0 | Ligand-dependent corepressor | May act as transcription activator that binds DNA elements with the sequence 5'-CCCTATCGATCGATCTCTACCT-3' | A0PJQ8 | - | A0A1L8FEN4 | - | - |
| 91 | KRRKK RQKLKEKK | 78 | Q9H875 | PRKR-interacting protein 1 | Binds double-stranded RNA. Inhibits EIF2AK2 kinase activity | B2GRY3 | C3Z487 | A0A1L8HDZ0 | - | Q5ZKU0 |
| 92 | ELRLKEELLK GIYA | 79 | Q9UHL0 | ATP-dependent RNA helicase DDX25 | ATP-dependent RNA helicase. Required for mRNA export and translation regulation during spermatid development | Q8JGR1 | C3YRH2 | A0A1L8FPD8 | - | A0A1D5NVY9 |
| 93 | AR RRR | 80 | Q9NUL7 | Probable ATP-dependent RNA helicase DDX28 | Plays an essential role in facilitating the proper assembly of the mitochondrial large ribosomal subunit and its helicase activity is essential for this function | Q08BL1 | C3XRL7 | A0A1L8GKB9 | - | - |
| 94 | RRAKWR RHQR | 81 | Q9BQY4 | Rhox homeobox family member 2 | Transcription factor maybe involved in reproductive processes. Modulates expression of target genes encoding proteins involved in processes relevant to spermatogenesis. | Q6P295 | - | - | - | - |
| 95 | R KHVKAHSAKE QQVRKK | 82 | Q8NBF1 | Zinc finger protein GLIS1 | Acts as both a repressor and activator of transcription. Binds to the consensus sequence 5'-GACCACCCAC-3' | F1QRD5 | C3XY75 | A0A1L8HY67 | - | F1ND41 |
| 96 | KRKR | 83 | Q04206 | Transcription factor p65 | NF-kappa-B is a pleiotropic transcription factor present in almost all cell types and is the endpoint of a series of signal transduction events that are initiated by a vast array of stimuli related to many biological processes such as inflammation, immunity, differentiation, cell growth, tumorigenesis and apoptosis. | Q6DGE6 | C3XR39 | A0A1L8GK02 | K7FNJ5 | P98152 |
| 97 | RLKRIWNWRA CEAAKKTKAD DR | 84 | O60563 | Cyclin-T1 | Regulatory subunit of the cyclin-dependent kinase pair (CDK9/cyclin-T1) complex, also called positive transcription elongation factor B (P-TEFb), which is proposed to facilitate the transition from abortive to productive elongation by phosphorylating the CTD (carboxy-terminal domain) of the large subunit of RNA polymerase II (RNA Pol II). | Q803Q8 | - | A0A1L8HHN4 | K7F433 | F1NIJ9 |
| 98 | KRRKKK | 85 | Q01201 | Transcription factor RelB | NF-kappa-B is a pleiotropic transcription factor which is present in almost all cell types and is involved in many biological processed such as inflammation, immunity, differentiation, cell growth, tumorigenesis and apoptosis. | Q0II00 | C3XR39 | P51510 | K7FNJ5 | Q04861 |
| 99 | MPPKGKSGSG KAGKGGAASG SDSADKK | 86 | Q9Y237 | Peptidyl-prolyl cis-trans isomerase NIMA-interacting 4 | Involved as a ribosomal RNA processing factor in ribosome biogenesis. Binds to tightly bent AT-rich stretches of double-stranded DNA | Q503Y7 | - | - | - | - |
| 100 | RRRR | 87 | Q9UGN5 | Poly [ADP-ribose] polymerase 2 | Involved in the base excision repair (BER) pathway, by catalyzing the poly(ADP-ribosyl)ation of a limited | Q7ZVB0 | C3Z2P3 | Q5M9A2 | - | E1BSI0 |
| 101 | KKTR R | | | | | | | | | |
| 102 | K AKKQK | 88 | Q04864 | Proto-oncogene c-Rel | Proto-oncogene that may play a role in differentiation and lymphopoiesis. | Q6K199 | - | Q91833 | K7FNJ5 | - |
| 103 | KKLQEQ EK | 89 | O75391 | Sperm-associated antigen 7 | Nucleic acid binding | Q7SYJ9 | C3XPX9 | A0A1L8EK33 | - | - |
| 104 | RRGEEWDPQ KAEEKRKLK | | | | | | | | | |
| 105 | KKGR RNRFK | 90 | P20823 | Hepatocyte nuclear factor 1-alpha | Transcriptional activator that regulates the tissue specific expression of multiple genes, especially in pancreatic islet cells and in liver. Required for the expression of several liver specific genes. Binds to the inverted palindrome 5'-GTTAATNATTAAC-3'. | Q8UVH4 | - | - | - | - |
| 106 | RK KVKYRK | 91 | O94983 | Calmodulin-binding transcription activator 2 | Transcription activator. May act as tumor suppressor. | UPI0004F4AA16 | C3ZIH0 | A0A1L8H472 | - | A0A1D5PBI0 |
| 107 | KKQESNHLQI HLCRKK | 92 | Q9NZ71 | Regulator of telomere | ATP-dependent DNA helicase implicated in telomere- | P0C928 | C3YXX3 | A0A1L8ET84 | K7G641 | F1NE49 |

| | | | | | | | | | | |
|---|---|---|---|---|---|---|---|---|---|---|
| 108 | RGGRKK | | | elongation helicase 1 | length regulation, DNA repair and the maintenance of [illegible] | | | | | |
| 109 | RKKVKYRK | 93 | Q9Y6Y1 | Calmodulin-binding transcription activator 1 | Transcriptional activator. May act as a tumor suppressor. | UPI00038359AD | C3ZIH0 | A0A1L8H472 | - | A0A1D5PBI0 |
| 110 | KIKEL YRRR | 94 | O75925 | E3 SUMO-protein ligase PIAS1 | Functions as an E3-type small ubiquitin-like modifier (SUMO) ligase, stabilizing the interaction between [illegible] | X1WDM3 | C3Z9X9 | Q5XHH3 | K7FKR8 | A0A1D5P1W4 |
| 111 | KKP TWVCPVCDKK | | | | | | | | | |
| 112 | KRKADGY NQPDSKRR | 95 | O60506 | Heterogeneous nuclear ribonucleoprotein Q | Heterogenous nuclear ribonucleoprotein (hnRNP) implicated in mRNA processing mechanisms. Component of the CRD-mediated complex that promotes MYC mRNA stability. | Q7SZC9 | C3ZLB6 | Q6NS22 | - | A0A1D5PJG4 |
| 113 | KD KKKK | 96 | Q08211 | ATP-dependent RNA helicase A | Multifunctional ATP-dependent nucleic acid helicase that unwinds DNA and RNA in a 3' to 5' direction and that plays important roles in many processes, such as DNA replication, transcriptional activation, post-transcriptional RNA regulation, mRNA translation and RNA-mediated gene silencing | E7F525 | C3Y332 | A0A1L8I2C1 | K7FFE8 | E1C388 |
| 114 | KR KQLHQLLPNH VLQKKKKH | 97 | P51003 | Poly(A) polymerase alpha | Polymerase that creates the 3'-poly(A) tail of mRNA's. Also required for the endoribonucleolytic cleavage reaction at some polyadenylation sites. May acquire specificity through interaction with a cleavage and polyadenylation specificity factor (CPSF) at its C-terminus. | Q5SNY2 | C3Z3M0 | A0A1L8F081 | - | A0A1D5P5T8 |
| 115 | KKRVHR | 98 | O75840 | Krueppel-like factor 7 | Transcriptional activator. Binds in vitro to the CACCC motif of the beta-globin promoter and to the SP1 recognition sequence. | Q502L8 | C3ZT77 | A0A1L8EX25 | K7FC91 | F1P0T6 |
| 116 | KRKEVKR | 99 | Q14872 | Metal regulatory transcription factor 1 | Activates the metallothionein I promoter. Binds to the metal responsive element (MRE). | A0A0G2KFP7 | C3YX65 | Q0VGW3 | K7FDS6 | Q5ZLE7 |
| 117 | KRKNAKMI SDIEKKRQRM | 100 | Q71F23 | Centromere protein U | Component of the CENPA-NAC (nucleosome-associated) complex, a complex that plays a central role [illegible] | E7FCV0 | - | A0A1L8HU11 | - | Q2Z1W2 |
| 118 | RRRPR PHRSEGARRS KNT | | | | | | | | | |
| 119 | KRPR | 101 | O15151 | Protein Mdm4 | Inhibits p53/TP53- and TP73/p73-mediated cell cycle arrest and apoptosis by binding its transcriptional activation domain. Inhibits degradation of MDM2. | F1QD25 | C3Y7L6 | Q7ZYI3 | K7GDW4 | F1NGX6 |
| 120 | KKFGAQNV ARRIEFRKK | 102 | Q9NYB0 | Telomeric repeat-binding factor 2-interacting protein 1 | Acts both as a regulator of telomere function and as a transcription regulator. | A0A0R4IRZ2 | C3XRX4 | A0A1L8ERQ2 | K7FQB6 | Q7T0L4 |
| 121 | AVSVI TISSDTDEEE TSQRHSLREC KGSLDCEACQ STLNIDRMCS LSSPDST | 103 | Q9H422 | Homeodomain-interacting protein kinase 3 | Serine/threonine-protein kinase involved in transcription regulation, apoptosis and steroidogenic gene expression. | F6NWN5 | C3Y5H6 | A0A1L8GJ10 | K7G117 | E1BUN6 |
| 122 | KRPR | 104 | Q06587 | E3 ubiquitin-protein ligase RING1 | Constitutes one of the E3 ubiquitin-protein ligases that mediate monoubiquitination of 'Lys-119' of histone H2A, thereby playing a central role in histone code and gene regulation. | A0A0H2UKC6 | C3ZPC6 | A0A1L8F8A8 | K7G2D5 | A0A1D5P1F4 |
| 123 | KKTSGL QQKNVE | 105 | Q9UBU8 | Mortality factor 4-like protein 1 | Component of the NuA4 histone acetyltransferase (HAT) complex which is involved in transcriptional activation of select genes principally by acetylation of nucleosomal histones H4 and H2A. | A8E7S8 | C3ZKT2 | Q66J61 | K7GG49 | A0A1L1RTQ0 |
| 124 | KKERKRL KEEKQKRKKE | 106 | Q8TAE8 | Growth arrest and DNA damage-inducible proteins-interacting protein 1 | Acts as a negative regulator of G1 to S cell cycle phase progression by inhibiting cyclin-dependent kinases. | UPI0002026CE4 | C3YM53 | - | - | - |
| 125 | RCVLAACSTY FKKLFKK | 107 | O43829 | Zinc finger and BTB domain-containing protein 14 | Transcriptional activator of the dopamine transporter (DAT), binding it's promoter at the consensus sequence 5'-CCTGCACAGTTCACGGA-3'. | Q7SYJ3 | C3ZBA4 | A0A1L8F220 | K7FD47 | Q92010 |
| 126 | KKKRLR | 108 | Q5T1R4 | Transcription factor HIVEP3 | Plays a role of transcription factor; binds to recognition signal sequences (Rss heptamer) for somatic recombination of immunoglobulin and T-cell receptor gene segments; Binds also to the kappa-B motif of gene such as S100A4, involved in cell progression and differentiation. | A0A286Y903 | - | Q5MYB9 | K7GD73 | F1P1B9 |
| 127 | KTPR K | 109 | Q02241 | Kinesin-like protein KIF23 | Component of the centralspindlin complex that serves as a microtubule-dependent and Rho-mediated signaling required for the myosin contractile ring formation during the cell cycle cytokinesis. | A0A286Y967 | C3Z0T9 | A0A1L8H194 | - | A0A1D5PCT6 |
| 128 | SKP KKK | 110 | Q9NZJ0 | Denticleless protein homolog | Substrate-specific adapter of a DCX (DDB1-CUL4-X-box) E3 ubiquitin-protein ligase complex required for cell cycle control, DNA damage response and translesion DNA synthesis. | Q5RHI5 | C3Y089 | Q6GPU3 | K7FX52 | Q5ZJW8 |
| 129 | KK SFSTLFLETV KRK | 111 | Q03188 | Centromere protein C | Component of the CENPA-NAC (nucleosome- | UPI0004572D87 | - | C8YR60 | K7F4L0 | O57392 |

| | | | | | | | | | | |
|---|---|---|---|---|---|---|---|---|---|---|
| 130 | KKSSTRK DKEESKKKR | | | | associated) complex, a complex that plays a central role in assembly of kinetochore proteins, mitotic progression and chromosome segregation. | | | | | |
| 131 | RKS TKKTNQSSKN IRKK | | | | | | | | | |
| 132 | KRKAKENIGKV NKKSNKKR | | | | | | | | | |
| 133 | RQ RKRHK | 112 | Q00987 | E3 ubiquitin-protein ligase Mdm2 | E3 ubiquitin-protein ligase that mediates ubiquitination of p53/TP53, leading to its degradation by the | A0A0R4ISQ1 | C3Y7L6 | A0A1L8GYI7 | K7GDW4 | F1NGX6 |
| 134 | KKLKK RNK | | | | | | | | | |
| 135 | RKRPRR | 113 | Q96EB6 | NAD-dependent protein deacetylase sirtuin-1 | NAD-dependent protein deacetylase that links transcriptional regulation directly to intracellular | A1L2B7 | C3Z3T3 | Q5U250 | - | A0A1P8D6T9 |
| 136 | KRKKRK | | | | | | | | | |
| 137 | KRKAKA | 114 | Q15025 | TNFAIP3-interacting protein 1 | Inhibits NF-kappa-B activation and TNF-induced NF-kappa-B-dependent gene expression by regulating A20/TNFAIP3-mediated deubiquitination of IKBKG; proposed to link A20/TNFAIP3 to ubiquitinated IKBKG. | A1L1N2 | C3Z5J9 | A0A1L8GWT2 | K7G737 | A0A1D5PHA8 |
| 138 | RRKRSKSE DMDNVQSKRR R | 115 | Q6VMQ6 | Activating transcription factor 7-interacting protein 1 | Recruiter that couples transcriptional factors to general transcription apparatus and thereby modulates transcription regulation and chromatin formation. | A0JME2 | - | A0A1L8GNC0 | K7F6D4 | - |
| 139 | KKRK PK | 116 | Q13769 | THO complex subunit 5 homolog | Acts as component of the THO subcomplex of the TREX complex which is thought to couple mRNA transcription, processing and nuclear export, and which specifically associates with spliced mRNA and not with unspliced pre-mRNA. | Q6NY52 | C3Z469 | A0A1L8HZU7 | K7FFP2 | Q5ZJK1 |
| 140 | KLLKDLPEL | 117 | Q53EL6 | Programmed cell death protein 4 | Inhibits translation initiation and cap-dependent translation. | Q7SYL0 | C3Y3Z6 | Q7T0M4 | K7FSZ2 | Q98TX3 |
| 141 | KAK RRLRK | | | | | | | | | |
| 142 | RRKH F | 118 | Q8N9U0 | Tandem C2 domains nuclear protein | Many C2 domains are Ca2+-dependent membrane-targeting modules that bind a wide variety of substances including bind phospholipids, inositol polyphosphates, and intracellular proteins. Most C2 domain proteins are either signal transduction enzymes that contain a single C2 domain, such as protein kinase C, or membrane trafficking proteins which contain at least two C2 domains, such as synaptotagmin 1. | UPI0004F450CE | - | A0A1L8F9H6 | K7G6Y4 | A0A1D5NW06 |
| 143 | K IK | 119 | Q16644 | MAP kinase-activated protein kinase 3 | Stress-activated serine/threonine-protein kinase involved in cytokines production, endocytosis, cell | Q6DHN7 | C3YQW4 | Q6IRB4 | K7F3M5 | E1BT35 |
| 144 | KRRKK | | | | | | | | | |
| 145 | RRR TVQEARALLR CRRAG | 120 | Q96S44 | EKC/KEOPS complex subunit TP53RK | Component of the EKC/KEOPS complex that is required for the formation of a threonylcarbamoyl group on adenosine at position 37 (t6A37) in tRNAs that read codons beginning with adenine | F1QB90 | C3ZAV8 | Q4V7S0 | - | - |
| 146 | KKQALKEKE LGNDAYKKK | 121 | P31948 | Stress-induced-phosphoprotein 1 | Acts as a co-chaperone for HSP90AA1 | Q5RKM3 | C3Y009 | Q7ZWU1 | - | - |
| 147 | RRLRIWR | 122 | Q7KZF4 | Staphylococcal nuclease domain-containing protein 1 | Endonuclease that mediates miRNA decay of both protein-free and AGO2-loaded miRNAs | Q5RGK8 | C3Y7K6 | Q7ZY98 | - | - |
| 148 | KKL RP | | | | | | | | | |
| 149 | KKER RVGTPQSTKK KKESRR | 123 | P54274 | Telomeric repeat-binding factor 1 | Binds the telomeric double-stranded 5'-TTAGGG-3' repeat and negatively regulates telomere length. | Q6NW47 | C3ZQ21 | Q1WM12 | - | F1NSI5 |
| 150 | K RRK | 124 | Q01658 | Protein Dr1 | The association of the DR1/DRAP1 heterodimer with TBP results in a functional repression of both activated and basal transcription of class II genes. This interaction precludes the formation of a transcription-competent complex by inhibiting the association of TFIIA and/or TFIIB with TBP. | Q08BR1 | C3ZFY4 | A0A1L8GMS5 | - | Q5ZMV3 |
| 151 | RKGKGQ IEKRKLREKR R | 125 | Q96IZ0 | PRKC apoptosis WT1 regulator protein | Pro-apoptopic protein capable of selectively inducing apoptosis in cancer cells, sensitizing the cells to diverse apoptotic stimuli and causing regression of tumors in animal models. | Q5XJ95 | - | Q66JA2 | - | F1NA74 |
| 152 | KRRQTSMTDF YHSKRR | 126 | P38936 | Cyclin-dependent kinase inhibitor 1 | May be involved in p53/TP53 mediated inhibition of cellular proliferation in response to DNA damage. | Q5RGY9 | C3YUU6 | Q91646 | K7FV40 | Q8JIV2 |
| 153 | KKNAEKEKQ QRNQKKKK | 127 | Q9BXJ9 | N-alpha-acetyltransferase 15, NatA auxiliary subunit | Auxillary subunit of the N-terminal acetyltransferase A (NatA) complex which displays alpha (N-terminal) acetyltransferase activity. | F1QE25 | C3XX70 | A0A1L8HUJ1 | K7FHQ9 | F1NVR6 |
| 154 | KRSVVSFDKVKE PRKSR | 128 | Q15424 | Scaffold attachment factor B1 | Binds to scaffold/matrix attachment region (S/MAR) DNA and forms a molecular assembly point to allow the formation of a 'transcriptosomal' complex (consisting of SR proteins and RNA polymerase II) coupling transcription and RNA processing | Q803X6 | C3Z0S4 | A0A1L8HWX4 | K7FD64 | H9L022 |
| 155 | KKRK | 129 | Q04724 | Transducin-like enhancer protein 1 | Transcriptional corepressor that binds to a number of transcription factors. | O13168 | C3ZWE3 | B1H1Y7 | - | A0A1D5PGR2 |

| | | | | | | | | | | |
|---|---|---|---|---|---|---|---|---|---|---|
| 156 | RASIER KRQRALMLRQ ARLAARP | 130 | P23025 | DNA repair protein complementing XP-A cells | Involved in DNA excision repair. Initiates repair by binding to damaged sites with various affinities, depending on the photoproduct and the transcriptional state of the region. Required for UV-induced CHEK1 phosphorylation and the recruitment of CEP164 to cyclobutane pyrimidine dimmers (CPD), sites of DNA damage after UV irradiation. | Q7SY02 | C3YQX9 | P27088 | - | R4GJS9 |
| 157 | KKAKLR | 131 | Q99661 | Kinesin-like protein KIF2C | In complex with KIF18B, constitutes the major microtubule plus-end depolymerizing activity in mitotic cells | F1RAK1 | C3YLZ9 | Q91636 | K7F465 | A0A1D5PBU5 |
| 158 | KRKR | 132 | Q9H814 | Phosphorylated adapter RNA export protein | A phosphoprotein adapter involved in the XPO1-mediated U snRNA export from the nucleus. Bridge | Q68EH5 | C3YBG9 | Q5U568 | - | Q5ZLY0 |
| 159 | KRK R | | | | | | | | | |
| 160 | KK RSR | 133 | Q8IXJ9 | Putative Polycomb group protein ASXL1 | Probable Polycomb group (PcG) protein involved in transcriptional regulation mediated by ligand-bound nuclear hormone receptors, such as retinoic acid receptors (RARs) and peroxisome proliferator-activated receptor gamma (PPARG). | F1Q5H5 | C3YBN3 | A0A1L8ERK1 | - | A0A1D5PLQ0 |
| 161 | GGKPCSQHI ISVTGFVDS | 134 | Q6ZW49 | PAX-interacting protein 1 | Involved in DNA damage response and in transcriptional regulation through histone methyltransferase (HMT) complexes. | A0A0R4ID17 | C3YDM2 | A0A1L8FQI7 | - | A0A1D5NZ11 |
| 162 | KQRA | 135 | Q04725 | Transducin-like enhancer protein 2 | Transcriptional corepressor that binds to a number of transcription factors. Inhibits the transcriptional activation mediated by CTNNB1 and TCF family members in Wnt signaling. The effects of full-length TLE family members may be modulated by association with dominant-negative AES | F1R8R4 | C3ZWE3 | A0A1L8HPC6 | - | A0A1D5PVL6 |
| 163 | R LRRAARPHRR LGPTGKEVHA LKRLR | 136 | Q6NXT1 | Ankyrin repeat domain-containing protein 54 | Plays an important role in regulating intracellular signaling events associated with erythroid terminal differentiation. | Q6DGX3 | C4A083 | A0A1L8GNU4 | - | F1NBL4 |
| 164 | RLKAK | 137 | Q99697 | Pituitary homeobox 2 | Controls cell proliferation in a tissue-specific manner and is involved in morphogenesis. During embryonic development, exerts a role in the expansion of muscle progenitors. | Q9W5Z2 | C3Z639 | Q9PWR3 | K7F389 | O93385 |
| 165 | RLKSK | 138 | P78337 | Pituitary homeobox 1 | Sequence-specific transcription factor that binds gene promoters and activates their transcription. | B2LT56 | C3Z639 | Q6GNM5 | K7F389 | A0A1D5P373 |
| 166 | RRARGGAV SARYVLPDEA ARAR | 139 | Q16690 | Dual specificity protein phosphatase 5 | Dual specificity protein phosphatase; active with phosphotyrosine, phosphoserine and phosphothreonine residues. The highest relative activity is toward ERK1. | Q6DBR5 | C3Z8W6 | A0A1L8HWG3 | - | Q9PW71 |
| 167 | RKVGRSLQRG CRALR | 140 | Q15628 | Tumor necrosis factor receptor type 1-associated DEATH domain protein | The nuclear form acts as a tumor suppressor by preventing ubiquitination and degradation of isoform p19ARF/ARF of CDKN2A by TRIP12: acts by interacting with TRIP12, leading to disrupt interaction between TRIP12 and isoform p19ARF/ARF of CDKN2A | B2GNQ4 | - | A0A1L8GKL2 | K7F8S3 | - |
| 168 | RL KAK | 141 | Q9Y2V3 | Retinal homeobox protein Rx | Plays a critical role in eye formation by regulating the initial specification of retinal cells and/or their subsequent proliferation. Binds to the photoreceptor conserved element-I (PCE-1/Ret 1) in the photoreceptor cell-specific arrestin promoter. | O42358 | C3ZEF8 | Q0ZLI0 | K7FM49 | Q9PVY0 |
| 169 | RKRK | 142 | Q9UKZ4 | Teneurin-1 | Teneurin C-terminal-associated peptide: Plays a role in the regulation of neuroplasticity in the limbic system. Mediates a rapid reorganization of actin- and tubulin-based cytoskeleton elements with an increase in dendritic arborization and spine density formation of neurons in the hippocampus and amygdala. | A0A286Y8Z9 | C3YZB2 | A0A1L8HUA7 | - | F1NN19 |
| 170 | RK AKKDDQMLKR RNVS | 143 | P52292 | Importin subunit alpha-1 | Functions in nuclear protein import as an adapter protein for nuclear receptor KPNB1. | Q6DI01 | C3XTJ8 | Q6IRB7 | K7FQJ3 | Q5ZLF7 |
| 171 | RKPVTA QERQREREEK RRRRQERAKE REKRRQERER | 144 | Q13164 | Mitogen-activated protein kinase 7 | Plays a role in various cellular processes such as proliferation, differentiation and cell survival. The upstream activator of MAPK7 is the MAPK kinase MAP2K5. Upon activation, it translocates to the nucleus and phosphorylates various downstream targets including MEF2C. | Q7ZVK8 | C3Z484 | Q6DFK6 | K7FFZ9 | F1N862 |
| 172 | QLFKRRNV A | 145 | P52294 | Importin subunit alpha-5 | Functions in nuclear protein import as an adapter protein for nuclear receptor KPNB1. | A0A0R4I9E5 | C3ZM42 | Q70PC5 | K7GFN3 | Q5ZML1 |

| | | | | | | | | | | |
|---|---|---|---|---|---|---|---|---|---|---|
| 173 | KKS ILKKKEQSH | 146 | Q9HBY8 | Serine/threonine-protein kinase Sgk2 | Serine/threonine-protein kinase which is involved in the regulation of a wide variety of ion channels, membrane transporters, cell growth, survival and proliferation. Up-regulates Na+ channels: SCNN1A/ENAC, K+ channels: KCNA3/Kv1.3, KCNE1 and KCNQ1, amino acid transporter: SLC6A19, glutamate transporter: SLC1A6/EAAT4, glutamate receptors: GRIA1/GLUR1 and GRIK2/GLUR6, Na+/H+ exchanger: SLC9A3/NHE3, and the Na+/K+ ATPase. | Q08BW4 | C3Y291 | A0A1L8FZ68 | K7G4F8 | A0A1D5PWW9 |
| 174 | KKVK | 147 | P06748 | Nucleophosmin | Involved in diverse cellular processes such as ribosome biogenesis, centrosome duplication, protein chaperoning, histone assembly, cell proliferation, and regulation of tumor suppressors p53/TP53 and ARF. | E7FCU0 | - | Q7ZTP6 | - | P16039 |
| 175 | KYLPQ QQKKK | 148 | Q9UKK3 | Poly [ADP-ribose] polymerase 4 | NAD+ + (ADP-D-ribosyl)(n)-acceptor = nicotinamide + (ADP-D-ribosyl)(n+1)-acceptor. | UPI0007EE98CA | C3Y4W7 | A0A1L8HJ59 | - | A0A1D5PL17 |
| 176 | SAKRPK | 149 | Q09472 | Histone acetyltransferase p300 | Functions as histone acetyltransferase and regulates transcription via chromatin remodeling | A0A0R4IVZ7 | C3ZUG6 | A0A1L8GGS2 | - | E1BSS0 |
| 177 | KK KYHN | 150 | O43683 | Mitotic checkpoint serine/threonine-protein kinase BUB1 | Serine/threonine-protein kinase that performs 2 crucial functions during mitosis: it is essential for spindle-assembly checkpoint signaling and for correct chromosome alignment. | F1QNS6 | C3XYZ6 | Q8JGT8 | - | A0A1D5PR30 |
| 178 | K KKK | 151 | Q13107 | Ubiquitin carboxyl-terminal hydrolase 4 | Hydrolase that deubiquitinates target proteins such as the receptor ADORA2A, PDPK1 and TRIM21. | E7EZD6 | C3Z9Q5 | A0A1L8GHB6 | K7G710 | Q5ZKD8 |
| 179 | KKKRR | 152 | P21675 | Transcription initiation factor TFIID subunit 1 | Largest component and core scaffold of the TFIID basal transcription factor complex | Q1LYC2 | C3Y3A3 | A0A1L8F6Z1 | - | B5AHE5 |
| 180 | RRSKL RKPKPVPLRK | 153 | O75410 | Transforming acidic coiled-coil-containing protein 1 | Likely involved in the processes that promote cell division prior to the formation of differentiated tissues. | Q6Y8G7 | - | A0JMR5 | K7FXY6 | A0A1D5PMU3 |
| 181 | KKAKS RLITSGCKVK K | | | | | | | | | |
| 182 | NKGWSGPKPG F | 154 | Q9NUL3 | Double-stranded RNA-binding protein Staufen homolog 2 | RNA-binding protein required for the microtubule-dependent transport of neuronal RNA from the cell body to the dendrite. | F1QJT6 | C3ZTW2 | A0A1L8FT79 | K7FV25 | A0A1D5PAM6 |
| 183 | KKLPPLPV VEKPKLFFKK RPK | | | | | | | | | |
| 184 | KRKPQEESERAKSD ESIKEEDKDQ DEKRRR | 155 | P26358 | DNA (cytosine-5)-methyltransferase 1 | Methylates CpG residues. Preferentially methylates hemimethylated DNA. | B3DKL7 | C3YZQ2 | P79922 | - | Q92072 |
| 185 | KRGL SVDSAQEVKR FR | 156 | Q9Y4A5 | Transformation/transcription domain-associated protein | Adapter protein, which is found in various multiprotein chromatin complexes with histone acetyltransferase activity (HAT), which gives a specific tag for epigenetic transcription activation. | A0A0R4ITC5 | C3ZR41 | A0A1L8EQF0 | - | E1C796 |
| 186 | LGRR R | 157 | O95831 | Apoptosis-inducing factor 1, mitochondrial | Functions both as NADH oxidoreductase and as regulator of apoptosis. In response to apoptotic stimuli, it is released from the mitochondrion intermembrane space into the cytosol and to the nucleus, where it functions as a proapoptotic factor in a caspase-independent pathway. | F1RBU0 | C3YZW2 | A0A1L8F6P6 | K7FF97 | Q5ZHL7 |
| 187 | MANKRMKVKH | 158 | Q86T24 | Transcriptional regulator Kaiso | Transcriptional regulator with bimodal DNA-binding specificity. Binds to methylated CpG dinucleotides in the consensus sequence 5'-CGCG-3' and also binds to the non-methylated consensus sequence 5'-CTGCNA-3' also known as the consensus kaiso binding site (KBS). | Q6PUQ9 | C3ZBA4 | A0A1L8F758 | K7GIX9 | F6U492 |
| 188 | KRK VSSSKPEENK IR | 159 | Q03933 | Heat shock factor protein 2 | DNA-binding protein that specifically binds heat shock promoter elements (HSE) and activates transcription. In [illegible] | Q90XA8 | C3ZBQ1 | Q5EG56 | K7G0A2 | A0A1D5P919 |
| 189 | KRKRPL LLNTNGAQKK | | | | | | | | | |
| 190 | RLK AK | 160 | O75364 | Pituitary homeobox 3 | Transcriptional regulator which is important for the differentiation and maintenance of meso-diencephalic dopaminergic (mdDA) neurons during development. | B2GRZ8 | C3Z639 | Q9I8K3 | K7GG06 | E1C143 |
| 191 | RREELAQ IRKEEKEKKR RR | 161 | Q9Y5A7 | NEDD8 ultimate buster 1 | Specific down-regulator of the NEDD8 conjugation system. Recruits NEDD8, UBD, and their conjugates to the proteasome for degradation. Isoform 1 promotes the degradation of NEDD8 more efficiently than isoform 2 | A8WFT9 | C3YYI8 | A0A1L8FV51 | K7FNN9 | E1BZ00 |
| 192 | HLLKRRN VP | 162 | O00629 | Importin subunit alpha-3 | Functions in nuclear protein import as an adapter protein for nuclear receptor KPNB1. | Q6DI01 | C3XTJ8 | Q6IRB7 | K7FQJ3 | Q5ZLF7 |

| | | | | | | | | | | |
|---|---|---|---|---|---|---|---|---|---|---|
| 193 | K KRKK | 163 | P23528 | Cofilin-1 | Binds to F-actin and exhibits pH-sensitive F-actin depolymerizing activity. Regulates actin cytoskeleton dynamics. Important for normal progress through mitosis and normal cytokinesis. Plays a role in the regulation of cell morphology and cytoskeletal organization. Required for the up-regulation of atypical chemokine receptor ACKR2 from endosomal compartment to cell membrane, increasing its efficiency in chemokine uptake and degradation | Q6TH32 | C4A075 | A0A1L8GCW3 | K7F4Y0 | P21566 |
| 194 | RGKRKR | 164 | P20700 | Lamin-B1 | Lamins are components of the nuclear lamina, a fibrous layer on the nucleoplasmic side of the inner nuclear membrane, which is thought to provide a framework for the nuclear envelope and may also interact with chromatin. | Q803N8 | C3ZXL7 | Q8AVZ3 | K7FK39 | P14731 |
| 195 | K KKIQKQLEHL KKLRK | 165 | O15553 | Pyrin | Involved in the regulation of innate immunity and the inflammatory response in response to IFNG/IFN-gamma. Organizes autophagic machinery by serving as a platform for the assembly of ULK1, Beclin 1/BECN1, ATG16L1, and ATG8 family members and recognizes specific autophagy targets, thus coordinating target recognition with assembly of the autophagic apparatus and initiation of autophagy. Acts as an autophagy receptor for the degradation of several inflammasome components, including CASP1, NLRP1 and NLRP3, hence preventing excessive IL1B- and IL18-mediated inflammation | Q7ZU70 | - | A0A1L8HI93 | K7FKU0 | A5HUK4 |
| 196 | KLRK | 166 | Q2VIQ3 | Chromosome-associated kinesin KIF4B | Motor protein that translocates PRC1 to the plus ends of interdigitating spindle microtubules during the metaphase to anaphase transition, an essential step for the formation of an organized central spindle midzone and midbody and for successful cytokinesis. | F8W409 | C3Z125 | Q91784 | K7FYR1 | Q90640 |
| 197 | RK HKK | 167 | Q8IWS0 | PHD finger protein 6 | Transcriptional regulator that associates with ribosomal RNA promoters and suppresses ribosomal RNA (rRNA) | Q803W6 | C3ZZY1 | A0A1L8F3A3 | - | V9GVP7 |
| 198 | KSKK KSRKGRPRK | | | | | | | | | |
| 199 | RRMI NKIDKNEDRK KDLS | 168 | Q96RN5 | Mediator of RNA polymerase II transcription subunit 15 | Component of the Mediator complex, a coactivator involved in the regulated transcription of nearly all RNA polymerase II-dependent genes. | F1QFA9 | C3YXZ1 | A0A1L8I0E1 | - | F1P1D8 |
| 200 | KEVGVGFAT RK | 169 | P15090 | Fatty acid-binding protein, adipocyte | Lipid transport protein in adipocytes. Binds both long chain fatty acids and retinoic acid. Delivers long-chain fatty acids and retinoic acid to their cognate receptors in the nucleus. | Q503X5 | C3Y8B1 | Q6PGR8 | K7G6I3 | Q90X55 |
| 201 | KR KRKRV | 170 | Q9UHL9 | General transcription factor II-I repeat domain-containing protein 1 | May be a transcription regulator involved in cell-cycle progression and skeletal muscle differentiation. | E7F1R4 | - | A0A1L8HG89 | K7FZF2 | A0A1D5NWB4 |
| 202 | PAKRKC IF | 171 | O75928 | E3 SUMO-protein ligase PIAS2 | Functions as an E3-type small ubiquitin-like modifier (SUMO) ligase, stabilizing the interaction between UBE2I and the substrate, and as a SUMO-tethering factor. | A2CE48 | C3Z9X9 | Q6IRN4 | K7FKR8 | A0A1D5P1W4 |
| 203 | KRTW VQEEEDIHKE KKI | 172 | Q8NG31 | Kinetochore scaffold 1 | Performs two crucial functions during mitosis: it is essential for spindle-assembly checkpoint signaling and for correct chromosome alignment. | Q56A53 | - | - | - | - |
| 204 | KKKKRK | 173 | Q9ULX6 | A-kinase anchor protein 8-like | Could play a role in constitutive transport element (CTE)-mediated gene expression by association with | A2CEZ5 | - | A0A1L8GMX3 | K7F381 | Q5ZJ02 |
| 205 | KRKL | | | | | | | | | |
| 206 | RKRK | 174 | Q16665 | Hypoxia-inducible factor 1-alpha | Functions as a master transcriptional regulator of the adaptive response to hypoxia. | F1QNB9 | C3XS48 | Q6GP97 | K7FUG2 | Q9YIB9 |
| 207 | RRKEKSKR DREKDKKR | 175 | Q6GQQ9 | OTU domain-containing protein 7B | Negative regulator of the non-canonical NF-kappa-B pathway that acts by mediating deubiquitination of TRAF3, an inhibitor of the NF-kappa-B pathway, thereby acting as a negative regulator of B-cell responses. | X1WEZ5 | C3YXE8 | - | K7G872 | H9L017 |
| 208 | KKRKK | 176 | P42568 | Protein AF-9 | Chromatin reader component of the super elongation complex (SEC), a complex required to increase the catalytic rate of RNA polymerase II transcription by suppressing transient pausing by the polymerase at multiple sites along the DNA | Q568M5 | C3Y1V7 | Q68EY8 | K7FL35 | F1NLL5 |

| | | | | | | | | | | |
|---|---|---|---|---|---|---|---|---|---|---|
| 209 | KKRKK | 177 | Q9UIS9 | Methyl-CpG-binding domain protein 1 | Transcriptional repressor that binds CpG islands in promoters where the DNA is methylated at position 5 of cytosine within CpG dinucleotides. | Q7SZX5 | - | A0A1L8F5I6 | - | - |
| 210 | PSKRPKA | 178 | P78347 | General transcription factor II-I | Interacts with the basal transcription machinery by coordinating the formation of a multiprotein complex at the C-FOS promoter, and linking specific signal responsive activator complexes. | E7F1R4 | - | A0A1L8HG89 | K7FZF2 | A0A1D5NWB4 |
| 211 | RRYGPK | 179 | P50461 | Cysteine and glycine-rich protein 3 | Positive regulator of myogenesis. Acts as cofactor for myogenic bHLH transcription factors such as MYOD1, and probably MYOG and MYF6. | F1QJC0 | C3ZK45 | Q68FI9 | K7FLU1 | F1NWZ2 |
| 212 | KAKTEEK DLKKLKKQEK EEKDFRKKFK | 180 | O15117 | FYN-binding protein 1 | Acts as an adapter protein of the FYN and LCP2 signaling cascades in T-cells. Modulates the expression of interleukin-2 (IL-2). Involved in platelet activation. Prevents the degradation of SKAP1 and SKAP2. May play a role in linking T-cell signaling to remodeling of the actin cytoskeleton. | UPI0004F429CD | C3YID4 | A0A1L8I2G1 | - | E1C908 |
| 213 | RKRRK | 181 | Q8WW38 | Zinc finger protein ZFPM2 | Transcription regulator that plays a central role in heart morphogenesis and development of coronary vessels from epicardium, by regulating genes that are essential during cardiogenesis. | Q2QKJ8 | C3YFM2 | A0A1L8FZV8 | K7F9W8 | A0A1D5PTA6 |
| 214 | RKR GKPASGAGAG AGAGKRRRKA DSAGDRGKSK | 182 | O43818 | U3 small nucleolar RNA-interacting protein 2 | Component of a nucleolar small nuclear ribonucleoprotein particle (snoRNP) thought to participate in the processing and modification of pre-ribosomal RNA. | F1R3Q7 | C3Z2P5 | A0A1L8GHE5 | - | - |
| 215 | KRSE SWTLKKPKSV SKKLK | 183 | Q8IYM9 | E3 ubiquitin-protein ligase TRIM22 | Interferon-induced antiviral protein involved in cell innate immunity. The antiviral activity could in part be mediated by TRIM22-dependent ubiquitination of viral proteins. | A0A0R4ISE2 | - | A0A1L8HI93 | K7FKU0 | A0A1D5PYD2 |
| 216 | KKKTEGLVKL TPIDKRK | 184 | O15164 | Transcription intermediary factor 1-alpha | Transcriptional coactivator that interacts with numerous nuclear receptors and coactivators and modulates the transcription of target genes. | Q5RHY8 | C3ZKJ9 | Q56R14 | - | F1NBP6 |
| 217 | RAKKGRR | 185 | Q15326 | Zinc finger MYND domain-containing protein 11 | Chromatin reader that specifically recognizes and binds histone H3.3 trimethylated at 'Lys-36' (H3.3K36me3) and regulates RNA polymerase II elongation. | Q08CT0 | C3Z8L5 | A0A1L8FVB2 | K7FYY9 | E1BUC1 |
| 218 | PPS RRHHCRSKAK RSR | 186 | O15534 | Period circadian protein homolog 1 | Transcriptional repressor which forms a core component of the circadian clock. | F1QUT1 | C3XQ83 | Q9DG29 | - | A0A1D5P3Z1 |
| 219 | KRGI TNTLEESSSL KRKR | 187 | P28715 | DNA repair protein complementing XP-G cells | Single-stranded structure-specific DNA endonuclease involved in DNA excision repair. | F1QJS3 | C3ZF00 | A0A1L8HH45 | K7GJZ2 | Q4AEJ2 |
| 220 | KLVPGLFKNE MKRRR | 188 | P35226 | Polycomb complex protein BMI-1 | Component of a Polycomb group (PcG) multiprotein PRC1-like complex, a complex class required to maintain the transcriptionally repressive state of many genes, including Hox genes, throughout development. | B0UXV8 | C3YW14 | Q640D5 | K7FHU3 | F1NWQ4 |
| 221 | RADRD KKKSRKRH | 189 | P19447 | General transcription and DNA repair factor IIH helicase subunit XPB | ATP-dependent 3'-5' DNA helicase, component of the general transcription and DNA repair factor IIH (TFIIH) core complex, which is involved in general and transcription-coupled nucleotide excision repair (NER) of damaged DNA and, when complexed to CAK, in RNA transcription by RNA polymerase II. | Q6PFM0 | C3ZFA3 | A0A1L8EUJ7 | K7FSE8 | Q5ZMK5 |
| 222 | KKKKR K | 190 | Q92889 | DNA repair endonuclease XPF | Catalytic component of a structure-specific DNA repair endonuclease responsible for the 5-prime incision during DNA repair. Involved in homologous recombination that assists in removing interstrand cross-link | Q7SYE8 | C3ZNX8 | A0A1L8EXK1 | - | F1NAV2 |
| 223 | KRPRD DEEEEQKMRR KQT | 191 | Q8WYA6 | Beta-catenin-like protein 1 | Component of the PRP19-CDC5L complex that forms an integral part of the spliceosome and is required for activating pre-mRNA splicing. Participates in AID/AICDA-mediated Ig class switching recombination (CSR). May induce apoptosis. | Q7SYE8 | C4A0X5 | A0A1L8EXK1 | K7FZ60 | E1BW50 |
| 224 | KKRKAP | 192 | Q9P0K7 | Ankycorbin | Plays a role in actin regulation at the ectoplasmic specialization, a type of cell junction specific to testis. Important for establishment of sperm polarity and normal spermatid adhesion. May also promote integrity of Sertoli cell tight junctions at the blood-testis barrier. | F1R8J7 | C3YNR4 | A0A1L8I2I5 | K7G1V5 | A0A1D5P991 |

| | | | | | | | | | |
|---|---|---|---|---|---|---|---|---|---|
| 225 | KRFARGDKR GKLPRW | 193 | P18074 | General transcription and DNA repair factor IIH helicase subunit XPD | ATP-dependent 5'-3' DNA helicase, component of the general transcription and DNA repair factor IIH (TFIIH) core complex, which is involved in general and transcription-coupled nucleotide excision repair (NER) of damaged DNA and, when complexed to CAK, in RNA transcription by RNA polymerase II. | F1R345 | C3ZNC0 | A0A1L8HG08 | - | F1NN99 |
| 226 | RRRVRSF ISPIPSKR | 194 | Q9UGU0 | Transcription factor 20 | Transcriptional activator that binds to the regulatory region of MMP3 and thereby controls stromelysin expression. It stimulates the activity of various transcriptional activators such as JUN, SP1, PAX6 and [illegible] | E7F726 | - | 0A1L8GGN6 | - | E1BXI6 |
| 227 | KPKKQ RQRRERRKPG AQPRKRKTKQ | | | | | | | | | |
| 228 | RFKRRH RS | | | | | | | | | |
| 229 | RAKKVRR | 195 | Q9UEW8 | STE20/SPS1-related proline-alanine-rich protein kinase | May act as a mediator of stress-activated signals. Mediates the inhibition of SLC4A4, SLC26A6 as well as CFTR activities by the WNK scaffolds, probably through phosphorylation. | A5PLJ7 | C3ZQX3 | A0JPG0 | - | F1NSM5 |
| 230 | KKLY QVKGKK | 196 | P40121 | Macrophage-capping protein | Calcium-sensitive protein which reversibly blocks the barbed ends of actin filaments but does not sever preformed actin filaments. May play an important role in macrophage function. May play a role in regulating cytoplasmic and/or nuclear structures through potential interactions with actin. May bind DNA. | E7FC92 | C3YDZ9 | Q6GMC0 | K7FNI5 | P02640 |
| 231 | KRK PLP | 197 | P42681 | Tyrosine-protein kinase TXK | Non-receptor tyrosine kinase that plays a redundant role with ITK in regulation of the adaptive immune response. | O42200 | C3Z633 | A0A1L8HTV0 | - | E1BTT3 |
| 232 | KKRSK | 198 | Q01831 | DNA repair protein complementing XP-C cells | Involved in global genome nucleotide excision repair (GG-NER) by acting as damage sensing and DNA-binding factor component of the XPC complex. | Q1LVE4 | C3YAV9 | A0A1L8GPS3 | K7GA72 | E1BUG1 |
| 233 | KRPMEEDGEE KSPSKKKKK | 199 | Q12906 | Interleukin enhancer-binding factor 3 | RNA-binding protein that plays an essential role in the biogenesis of circular RNAs (circRNAs) which are produced by back-splicing circularization of pre-mRNAs. | Q6NXA4 | C4A0T6 | Q6DCD0 | - | A0A1D5NXI6 |
| 234 | RKRKK | 200 | P54132 | Bloom syndrome protein | ATP-dependent DNA helicase that unwinds single- and double-stranded DNA in a 3'-5' direction | Q498Z7 | C3YKR1 | Q32NN1 | K7F4Y6 | Q8AYS4 |
| 235 | KKKKK | 201 | P38935 | DNA-binding protein SMUBP-2 | 5' to 3' helicase that unwinds RNA and DNA duplices in an ATP-dependent reaction. | M9MM98 | C3YJY8 | A0A1L8GJ95 | - | E1BY42 |
| 236 | KDEVRRGD TSTEDIQEEK DKK | 202 | Q7Z401 | C-myc promoter-binding protein | Probable guanine nucleotide exchange factor (GEF) which may activate RAB10. Promotes the exchange of GDP to GTP, converting inactive GDP-bound Rab proteins into their active GTP-bound form. | A0A286Y972 | C3Y2D3 | A0A1L8HXV3 | K7FZL9 | A0A1D5PD21 |
| 237 | KKKKK | 203 | P00519 | Tyrosine-protein kinase ABL1 | Non-receptor tyrosine-protein kinase that plays a role in many key processes linked to cell growth and survival such as cytoskeleton remodeling in response to [illegible] | A0A0R4IYB8 | - | Q6P282 | K7FHZ2 | F1NCD9 |
| 238 | S KRFLR | | | | | | | | | |
| 239 | PRKRAGE | | | | | | | | | |
| 240 | KKR REKQRRR | 204 | O43823 | A-kinase anchor protein 8 | Anchoring protein that mediates the subcellular compartmentation of cAMP-dependent protein kinase (PKA type II) | A0A0H2UKW6 | - | A0A1L8GMX3 | K7F381 | A0A1D5PZG6 |
| 241 | K RKRKSR | 205 | Q8IWZ8 | SURP and G-patch domain-containing protein 1 | Plays a role in pre-mRNA splicing. | E7FGI4 | C3ZA13 | Q4V7M9 | - | A0A1D5P6Z4 |
| 242 | KKK RLR | 206 | P31629 | Transcription factor HIVEP2 | This protein specifically binds to the DNA sequence 5'-GGGACTTTCC-3' which is found in the enhancer elements of numerous viral promoters such as those of SV40, CMV, or HIV1. | F1Q5D6 | - | Q5MYB9 | K7FQK1 | F1P1B9 |
| 243 | KKEMIAQQR EYKKKKALKK | 207 | O75940 | Survival of motor neuron-related-splicing factor 30 | Necessary for spliceosome assembly. Overexpression causes apoptosis. | Q9W6S8 | C3Y8H2 | Q6NU78 | K7G033 | Q98SU9 |
| 244 | KRCIWSTPKR RHKKK | 208 | Q9HB58 | Sp110 nuclear body protein | Transcription factor. May be a nuclear hormone receptor coactivator. Enhances transcription of genes [illegible] | F1R6N2 | - | A0A1L8GBC5 | - | - |
| 245 | KKK EKDICSSSKR RFQK | | | | | | | | | |
| 246 | PGLTKRVK KSK | 209 | Q96EZ8 | Microspherule protein 1 | Modulates the transcription repressor activity of DAXX by recruiting it to the nucleolus | Q6P0E3 | C3YYL7 | Q7T0W8 | - | A0A1D5PQU2 |
| 247 | KKRKR | 210 | Q15637 | Splicing factor 1 | Necessary for the ATP-dependent first step of spliceosome assembly. Binds to the intron branch point sequence (BPS) 5'-UACUAAC-3' of the pre-mRNA. May act as transcription repressor. | A0A0R4IBT0 | C3YFS6 | - | - | - |

| | | | | | | | | | | |
|---|---|---|---|---|---|---|---|---|---|---|
| 248 | KKRNKKKK | 211 | Q9UBZ9 | DNA repair protein REV1 | Deoxycytidyl transferase involved in DNA repair. Transfers a dCMP residue from dCTP to the 3'-end of a DNA primer in a template-dependent reaction. May assist in the first step in the bypass of abasic lesions by the insertion of a nucleotide opposite the lesion. Required for normal induction of mutations by physical and chemical agents. | B0UY97 | C3Y6K1 | A0A1L8HGZ7 | - | F1P5H4 |
| 249 | RRWNK LR | 212 | Q03468 | DNA excision repair protein ERCC-6 | Essential factor involved in transcription-coupled nucleotide excision repair which allows RNA [illegible] | F1RDN1 | C3Z749 | A0A1L8FKT9 | - | E1BYA8 |
| 250 | KRRI QPAFGADHDV PKRKK | | | | | | | | | |
| 251 | RKRSR | 213 | O43502 | DNA repair protein RAD51 homolog 3 | Essential for the homologous recombination (HR) pathway of DNA repair. Involved in the homologous recombination repair (HRR) pathway of double-stranded DNA breaks arising during DNA replication or induced by DNA-damaging agents. | F1QQC9 | - | A0A1L8HG81 | K7G3U0 | A0A1L1RWC9 |
| 252 | KDKFN | 214 | Q9UBB9 | Tuftelin-interacting protein 11 | Involved in pre-mRNA splicing, specifically in spliceosome disassembly during late-stage splicing events. | Q6DI35 | - | Q66J74 | - | Q5ZII9 |
| 253 | KEKFKENGKM FKDK | 215 | Q03112 | MDS1 and EVI1 complex locus protein | Functions as a transcriptional regulator binding to DNA sequences in the promoter region of target genes and regulating positively or negatively their expression. Oncogene which plays a role in development, cell proliferation and differentiation. May also play a role in apoptosis through regulation of the JNK and TGF-beta signaling. Involved in hematopoiesis. | F1Q833 | C3XYB0 | - | K7FB80 | A0A1D5P950 |
| 254 | PSRKRPR | 216 | Q92833 | Protein Jumonji | Regulator of histone methyltransferase complexes that plays an essential role in embryonic development, including heart and liver development, neural tube fusion process and hematopoiesis. | Q1LVC2 | C3ZYE0 | A0A1L8FY27 | - | Q5F363 |
| 255 | KK TGKNRKLKSK RVKPR | 217 | O15055 | Period circadian protein homolog 2 | Transcriptional repressor which forms a core component of the circadian clock. | F1QUT1 | C3XQ83 | Q9DG29 | - | A0A1D5P3Z1 |
| 256 | RK KRRRQKSRK | 218 | Q5H9F3 | BCL-6 corepressor-like protein 1 | Transcriptional corepressor. May specifically inhibit gene expression when recruited to promoter regions by sequence-specific DNA-binding proteins such as BCL6. This repression may be mediated at least in part by histone deacetylase activities which can associate with this corepressor. | F1RCL8 | - | A0A1L8F7G7 | K7G927 | A0A1D5P6S5 |
| 257 | RKRPL EAPPEAGSTK RTN | 219 | P51513 | RNA-binding protein Nova-1 | May regulate RNA splicing or metabolism in a specific subset of developing neurons | Q1LYC7 | - | Q6GPZ4 | K7FI95 | R4GG35 |
| 258 | KRK MMQPFNKPSG TFIKKPKLAK | 220 | Q5BKZ1 | DBIRD complex subunit ZNF326 | Core component of the DBIRD complex, a multiprotein complex that acts at the interface between core mRNP particles and RNA polymerase II (RNAPII) and integrates transcript elongation with the regulation of alternative splicing: the DBIRD complex affects local transcript elongation rates and alternative splicing of a large set of exons embedded in (A + T)-rich DNA regions. | A2CEZ5 | - | A0A1L8GMX3 | K7F381 | Q5ZJ02 |
| 259 | KRKRE KKEKKKKRK | 221 | P25440 | Bromodomain-containing protein 2 | Binds hyperacetylated chromatin and plays a role in the regulation of transcription, probably by chromatin remodeling. Regulates transcription of the CCND1 gene. Plays a role in nucleosome assembly. | A8CYR1 | C3ZPB3 | A0A1L8H2F1 | - | F1NS89 |
| 260 | KKP KRKKRRRR | 222 | Q96CK0 | Zinc finger protein 653 | Transcriptional repressor. May repress NR5A1, PPARG, NR1H3, NR4A2, ESR1 and NR3C1 [illegible] | UPI00038372B4 | C3ZAX3 | Q08B30 | - | - |
| 261 | KEPEKRR RSKRSR | | | | | | | | | |
| 262 | KRKIR | 223 | P0C1S8 | Wee1-like protein kinase 2 | Oocyte-specific protein tyrosine kinase that phosphorylates and inhibits CDK1 and acts as a key regulator of meiosis during both prophase I and metaphase II. | Q6DRI0 | C3Y7K0 | Q8AYK6 | K7FYG3 | Q5F400 |
| 263 | RKRRR | 224 | Q8WXF8 | DNA-binding death effector domain-containing protein 2 | May play a critical role in death receptor-induced apoptosis and may target CASP8 and CASP10 to the [illegible] | Q6DHN2 | C3XZG0 | Q5PQ51 | K7GE20 | A0A1D5PXF2 |
| 264 | KRQRRS RGRPSGGARR RRR | | | | | | | | | |
| 265 | KDKEKEK QRKEKDKTR | 225 | Q8TE49 | OTU domain-containing protein 7A | Has deubiquitinating activity towards 'Lys-11'-linked polyubiquitin chains. | X1WEZ5 | C3YXE8 | A0A1L8H082 | K7G872 | E1C900 |
| 266 | RKK | 226 | Q9Y473 | Zinc finger protein 175 | Down-regulates the expression of several chemokine receptors. Interferes with HIV-1 replication by suppressing Tat-induced viral LTR promoter activity. | A9JTD1 | C3XPD3 | A0A1L8FUG5 | K7FQR3 | Q5F3E3 |
| 267 | KKKTGARK K | 227 | Q9Y3S2 | Zinc finger protein 330 | zinc ion binding | Q7ZUZ8 | C3YZ79 | Q7ZXJ2 | K7GHZ0 | E1C2N4 |

| | | | | | | | | | | |
|---|---|---|---|---|---|---|---|---|---|---|
| 268 | KKHK | 228 | Q6ECI4 | Zinc finger protein 470 | May be involved in transcriptional regulation. | Q4V8R6 | C3XSV2 | Q6GN31 | K7GAQ9 | - |
| 269 | RHRR | | | | | | | | | |
| 270 | KRRKF | 229 | Q86X24 | HORMA domain-containing protein 1 | Plays a key role in meiotic progression. Regulates 3 different functions during meiosis: ensures that sufficient numbers of processed DNA double-strand breaks (DSBs) are available for successful homology search by increasing the steady-state numbers of single-stranded DSB ends. | A2BF66 | C3Y7J4 | A0A1L8F4F2 | - | E1C2C4 |
| 271 | K NKKKKEK | 230 | Q96T21 | Selenocysteine insertion sequence-binding protein 2 | Binds to the SECIS element in the 3'-UTR of some mRNAs encoding selenoproteins. Binding is stimulated by SELB. | Q7ZV23 | C3ZJL2 | A0A1L8HYH8 | K7GDC3 | F1NS30 |
| 272 | RREREAEHGY ASMLPYNNKD RDALKRRNK | 231 | Q05195 | Max dimerization protein 1 | Transcriptional repressor. MAD binds with MAX to form a sequence-specific DNA-binding protein complex which recognizes the core sequence 5'-CAC[GA]TG-3'. MAD thus antagonizes MYC transcriptional activity by competing for MAX. | A3KQG0 | C3Y472 | B7ZPQ0 | K7G942 | Q5ZJ78 |
| 273 | KKTDKLKKKK KRKRR | 232 | Q86YI8 | PHD finger protein 13 | Modulates chromatin structure. Required for normal chromosome condensation during the early stages of mitosis. Required for normal chromosome separation during mitosis. | A0A0G2L648 | C3ZH10 | Q5PQ67 | K7FYN5 | R4GGG1 |
| 274 | R KHVKAHSSKE QQARKKLR | 233 | Q8NEA6 | Zinc finger protein GLIS3 | Acts as both a repressor and activator of transcription. Binds to the consensus sequence 5'-GACCACCCAC-3' | A0A0R4IU95 | C3XY75 | A0A1L8HY67 | - | E1C1Q5 |
| 275 | KRPA SDMGKKPKTP KKKKKK | 234 | O94900 | Thymocyte selection-associated high mobility group box protein TOX | May play a role in regulating T-cell development. | B0V356 | - | A0A1L8FZ33 | K7FWE5 | A0A1D5PSC7 |
| 276 | KKKTTKL | 235 | Q96NK8 | Neurogenic differentiation factor 6 | Activates E box-dependent transcription in collaboration with TCF3/E47. May be a trans-acting factor involved in the development and maintenance of the mammalian nervous system. Transactivates the promoter of its own gene | O42202 | C3ZXA2 | A0A1L8HIF8 | K7EYZ4 | P79766 |
| 277 | KKRRH x{9}HK KERKVY | 236 | Q13627 | Dual specificity tyrosine-phosphorylation-regulated kinase 1A | Dual-specificity kinase which possesses both serine/threonine and tyrosine kinase activities. May play a role in a signaling pathway regulating nuclear functions of cell proliferation. Modulates alternative splicing by phosphorylating the splice factor SRSF6 | A1L1U2 | C3Z2G7 | A0A1L8HCE8 | K7FXM2 | Q8JG32 |
| 278 | KKVRK | 237 | P15923 | Transcription factor E2-alpha | Transcriptional regulator. Involved in the initiation of neuronal differentiation. | B1H1L4 | C3ZAG1 | A0A1L8HWH1 | K7F8X5 | A0A1D5P7W4 |
| 279 | VKKSIR | 238 | P06748 | Nucleophosmin | Involved in diverse cellular processes such as ribosome biogenesis, centrosome duplication, protein chaperoning, histone assembly, cell proliferation, and regulation of tumor suppressors p53/TP53 and ARF. | E7FCU0 | - | Q7ZTP6 | - | P16039 |
| 280 | KRKH RKIPFSKRK | 239 | Q9UKK3 | Poly [ADP-ribose] polymerase 4 | NAD+ ADP-ribosyltransferase activity | UPI0007EE98CA | C3Y4W7 | A0A1L8HJ59 | - | A0A1D5PL17 |
| 281 | KKNK | 240 | Q86Z02 | Homeodomain-interacting protein kinase 1 | Serine/threonine-protein kinase involved in transcription regulation and TNF-mediated cellular apoptosis. | E7F544 | C3Y5H6 | Q5U4V9 | K7F7D3 | E1BW82 |
| 282 | KKERNS SGMARKAKRT K | 241 | P35659 | Protein DEK | Involved in chromatin organization. | F1R6C4 | - | Q2KHP5 | K7G4P3 | Q5F422 |
| 283 | KRRKK | 242 | O75398 | Deformed epidermal autoregulatory factor 1 homolog | Transcription factor that binds to sequence with multiple copies of 5'-TTC[CG]G-3' present in its own promoter and that of the HNRPA2B1 gene. | B3DHN2 | C3ZC54 | A0A1L8GJK0 | - | A0A1L1RRQ7 |
| 284 | KKK | 243 | P35251 | Replication factor C subunit 1 | The elongation of primed DNA templates by DNA polymerase delta and epsilon requires the action of the accessory proteins PCNA and activator 1. | F1Q5P6 | C3Y4Y3 | A0A1L8HTJ6 | - | F1NEP9 |
| 285 | KIPKFKK | 244 | Q09666 | Neuroblast differentiation-associated protein AHNAK | May be required for neuronal cell differentiation. | F1QZ50 | C3YDK3 | A0A1L8F3W5 | - | - |
| 286 | KK SKFKMPK | | | | | | | | | |
| 287 | KIKAKK | | | | | | | | | |
| 288 | KLKKS KIKMPK | | | | | | | | | |
| 289 | KRPRK | 245 | Q15554 | Telomeric repeat-binding factor 2 | Binds the telomeric double-stranded 5'-TTAGGG-3' repeat and plays a central role in telomere maintenance and protection against end-to-end fusion of chromosomes. | Q6NW47 | C3ZQ21 | Q1WM12 | - | F1NSI5 |

| | | | | | | | | | | |
|---|---|---|---|---|---|---|---|---|---|---|
| 290 | KK RRR | 246 | Q86Y91 | Kinesin-like protein KIF18B | In complex with KIF2C, constitutes the major microtubule plus-end depolymerizing activity in mitotic cells. Its major role may be to transport KIF2C and/or MAPRE1 along microtubules | Q7ZUW9 | C3Y5L4 | A0A1L8GJ38 | K7F9Y7 | Q5ZLK6 |
| 291 | KKKKPK | 247 | Q8IX01 | SURP and G-patch domain-containing protein 2 | May play a role in mRNA splicing. | E7FGI4 | C3ZA13 | Q4V7M9 | - | E1C5N4 |
| 292 | KAKKRKR | 248 | O14776 | Transcription elongation regulator 1 | Transcription factor that binds RNA polymerase II and inhibits the elongation of transcripts from target promoters. Regulates transcription elongation in a TATA box-dependent manner. Necessary for TAT-dependent activation of the human immunodeficiency virus type 1 (HIV-1) promoter. | E7F3A6 | C3Y4A6 | - | K7GCU0 | A0A1D5NTY1 |
| 293 | R RREAAAEARK SHSPVKRPRK | 249 | P78549 | Endonuclease III-like protein 1 | Bifunctional DNA N-glycosylase with associated apurinic/apyrimidinic (AP) lyase function that catalyzes the first step in base excision repair (BER), the primary repair pathway for the repair of oxidative DNA damage. | F1QBP9 | C3Z7H4 | - | - | A7M7B9 |
| 294 | KK PKIRRKKS | 250 | Q7Z7C8 | Transcription initiation factor TFIID subunit 8 | Transcription factor TFIID is one of the general factors required for accurate and regulated initiation by RNA polymerase II. | Q6P0T2 | C3ZC47 | Q7ZYA2 | K7FSI0 | R4GIW1 |
| 295 | HRKRK R | 251 | Q7Z2E3 | Aprataxin | DNA-binding protein involved in single-strand DNA break repair, double-strand DNA break repair and base excision repair | P61799 | C3ZFB4 | Q7T287 | - | A0A1D5PAK1 |
| 296 | RPRRRSRR | 252 | Q7Z628 | Neuroepithelial cell-transforming gene 1 protein | Acts as guanine nucleotide exchange factor (GEF) for RhoA GTPase. May be involved in activation of the [illegible] | Q1LXH8 | C3Z2Q7 | A0A1L8GZH7 | K7FZV1 | Q5ZJM0 |
| 297 | KRKRR EK | | | | | | | | | |
| 298 | KR RRPSEGR | 253 | Q7Z5J4 | Retinoic acid-induced protein 1 | Transcriptional regulator of the circadian clock components: CLOCK, ARNTL/BMAL1, [illegible] | E7F726 | - | A0A1L8EYP3 | - | - |
| 299 | KRKSAFMA PVPTKKRNLV | | | | | | | | | |
| 300 | KKSSQTK KKKKAFNSPK | 254 | Q9NPC7 | Myoneurin | RNA polymerase II transcription factor activity, sequence-specific DNA binding | A9JSZ9 | C3XQ44 | A0A1L8HK33 | K7F9H0 | E1C3P7 |
| 301 | KRKR GK | | | | | | | | | |
| 302 | KRLEH TERQFRNRRK | 255 | Q8IY67 | Ribonucleoprotein PTB-binding 1 | Cooperates with PTBP1 to modulate regulated alternative splicing events. Promotes exon skipping. Cooperates with PTBP1 to modulate switching between mutually exclusive exons during maturation of the TPM1 pre-mRNA | F1QAP4 | C3YIG5 | A0A1L8GLV5 | - | A0A1D5P0I3 |
| 303 | KMISRKK KHQA | 256 | Q14517 | Protocadherin Fat 1 | Plays an essential role for cellular polarization, directed cell migration and modulating cell-cell contact. | A2BGT8 | C3Y3A1 | A0A1L8GWK1 | K7FFG3 | F1NWW5 |
| 304 | KRPAEPG PARVGKKGKK | 257 | Q8N543 | Prolyl 3-hydroxylase OGFOD1 | Prolyl 3-hydroxylase that catalyzes 3-hydroxylation of 'Pro-62' of small ribosomal subunit uS12 (RPS23), thereby regulating protein translation termination efficiency. Involved in stress granule formation | F1QTC6 | - | Q6DE73 | - | - |
| 305 | RK KF | 258 | P07992 | DNA excision repair protein ERCC-1 | Non-catalytic component of a structure-specific DNA repair endonuclease responsible for the 5'-incision during DNA repair. | Q6NY87 | C3Y937 | A0A1L8FKK3 | - | - |
| 306 | KKKK MTK | 259 | Q13562 | Neurogenic differentiation factor 1 | Acts as a transcriptional activator: mediates transcriptional activation by binding to E box-containing promoter consensus core sequences 5'-CANNTG-3'. | O42202 | - | A0A1L8ERW7 | K7F0H1 | P79765 |
| 307 | KKRKKK | 260 | O94842 | TOX high mobility group box family member 4 | Component of the PTW/PP1 phosphatase complex, which plays a role in the control of chromatin structure and cell cycle progression during the transition from mitosis into interphase. | F1QXS5 | - | A0A1L8HPM5 | K7FLV4 | A0A1D5PEI4 |
| 308 | RKKRGRK | 261 | Q14781 | Chromobox protein homolog 2 | Component of a Polycomb group (PcG) multiprotein PRC1-like complex, a complex class required to maintain the transcriptionally repressive state of many genes, including Hox genes, throughout development | Q8JIQ9 | - | A0A1L8EM82 | - | - |
| 309 | KRKRx{21}KFRKSKKKKR | 262 | P57052 | Splicing regulator RBM11 | Tissue-specific splicing factor with potential implication in the regulation of alternative splicing during neuron and germ cell differentiation. | Q6NWB3 | C3Z7L7 | Q6GM50 | K7FCD7 | A0A1L1RL91 |
| 310 | RKRLSD EKNLPLDGKR QRF | 263 | Q6PIY7 | Poly(A) RNA polymerase GLD2 | Cytoplasmic poly(A) RNA polymerase that adds successive AMP monomers to the 3'-end of specific RNAs, forming a poly(A) tail. In contrast to the canonical nuclear poly(A) RNA polymerase, it only adds poly(A) to selected cytoplasmic mRNAs | Q503I9 | C3Y1Z0 | A0A1L8I240 | K7FPN3 | E1C833 |

| | | | | | | | | | | |
|---|---|---|---|---|---|---|---|---|---|---|
| 311 | RKPR | 264 | Q9NR83 | SLC2A4 regulator | Transcription factor involved in SLC2A4 and HD gene transactivation. Binds to the consensus sequence 5'-GCCGGCG-3' | F1R143 | C3YKU5 | A0A1L8GC36 | K7FHK4 | A0A1D5PYE8 |
| 312 | KPKRKRR KKRGHGWSRM RMRR | 265 | Q13342 | Nuclear body protein SP140 | Component of the nuclear body, also known as nuclear domain 10, PML oncogenic domain, and KR body (PubMed:8910577). May be involved in the pathogenesis of acute promyelocytic leukemia and viral infection | A0A0R4IMH4 | - | A0A1L8GBC5 | - | - |
| 313 | KRKGPDS LSDGPACKR | 266 | Q9BYH8 | NF-kappa-B inhibitor zeta | Involved in regulation of NF-kappa-B transcription factor complexes. | UPI0007EEA726 | C3ZHK3 | A0A1L8H8D6 | K7GAL9 | F1NL80 |
| 314 | H KRKR | 267 | Q8NEA9 | Germ cell-less protein-like 2 | Possible function in spermatogenesis. Probable substrate-specific adapter of an E3 ubiquitin-protein [illegible] | Q6NYQ2 | C3Y031 | A0A1L8H1G4 | K7FJA3 | - |
| 315 | RKKL R | | | | | | | | | |
| 316 | KKRGPKKRK MTK | 268 | Q15784 | Neurogenic differentiation factor 2 | Transcriptional regulator implicated in neuronal determination. Mediates calcium-dependent transcription activation by binding to E box-containing promoter. | Q9W6C8 | C3ZXA2 | Q6GPW1 | K7EXA8 | P79765 |
| 317 | H KRKR | 269 | Q96IK5 | Germ cell-less protein-like 1 | Possible function in spermatogenesis. Enhances the degradation of MDM2 and increases the amount of p53 [illegible] | Q6NYQ2 | C3Y031 | A0A1L8H1G4 | K7FJA3 | - |
| 318 | PRRKKL K | | | | | | | | | |
| 319 | KRKNE KAGSKRKK | 270 | Q9Y3E1 | Hepatoma-derived growth factor-related protein 3 | Enhances DNA synthesis and may play a role in cell proliferation. | A0A0R4I9P3 | - | A0A1L8HXV6 | - | R4GLJ4 |
| 320 | FKRR | 271 | Q86VE0 | Myb-related transcription factor, partner of profilin | Transcriptional repressor; DNA-binding protein that specifically recognizes the core sequence 5'-YAAC[GT]G-3'. Dimerization with PFN1 reduces its [illegible] | UPI00038387C8 | - | A0A1L8F865 | - | - |
| 321 | HKRRK | | | | | | | | | |
| 322 | R KRR | | | | | | | | | |
| 323 | KK R | 272 | Q92630 | Dual specificity tyrosine-phosphorylation-regulated kinase 2 | Serine/threonine-protein kinase involved in the regulation of the mitotic cell cycle, cell proliferation, apoptosis, organization of the cytoskeleton and neurite outgrowth. | A0A0R4ITS5 | C3Y4L0 | Q5PPW5 | K7F6T0 | Q5ZIU3 |
| 324 | KKEHKKVK | 273 | P42166 | Lamina-associated polypeptide 2, isoform alpha | May be involved in the structural organization of the nucleus and in the post-mitotic nuclear assembly. Plays an important role, together with LMNA, in the nuclear anchorage of RB1.<br>TP and TP5 may play a role in T-cell development and function. TP5 is an immunomodulating pentapeptide | Q5RGX2 | - | - | - | - |
| 325 | QKRKT | 274 | P41218 | Myeloid cell nuclear differentiation antigen | May act as a transcriptional activator/repressor in the myeloid lineage. Plays a role in the granulocyte/monocyte cell-specific response to interferon. Stimulates the DNA binding of the transcriptional repressor protein YY1 | UPI00038371A7 | - | - | - | - |
| 326 | KKTQRRH | 275 | Q9P0U3 | Sentrin-specific protease 1 | Protease that catalyzes two essential functions in the SUMO pathway | A4IFY7 | C3Z396 | - | K7GGL1 | A0A1D5PIW4 |
| 327 | YF RKR | | | | | | | | | |
| 328 | KKRK | | | | | | | | | |
| 329 | RKKREK | 276 | P22626 | Heterogeneous nuclear ribonucleoproteins A2/B1 | Heterogeneous nuclear ribonucleoprotein (hnRNP) that associates with nascent pre-mRNAs, packaging them into hnRNP particles. | Q803K3 | - | Q6IP29 | - | Q5ZME1 |
| 330 | KKRKER | 277 | O43390 | Heterogeneous nuclear ribonucleoprotein R | Component of ribonucleosomes, which are complexes of at least 20 other different heterogenious nuclear ribonucleoproteins (hnRNP). hnRNP play an important role in processing of precursor mRNA in the nucleus. | Q7SZC9 | C3ZLB6 | A0A1L8G9H1 | - | A0A1D5P155 |
| 331 | RRRR | 278 | Q9H4L4 | Sentrin-specific protease 3 | Protease that releases SUMO2 and SUMO3 monomers from sumoylated substrates, but has only weak activity [illegible] | A2BEN4 | C3XYA3 | A0A1L8G3T3 | K7GGL1 | F1NYZ3 |
| 332 | WGRHRGRRR | | | | | | | | | |
| 333 | HKVLPR | 279 | P06401 | Progesterone receptor | The steroid hormones and their receptors are involved in the regulation of eukaryotic gene expression and affect cellular proliferation and differentiation in target tissues. Depending on the isoform, progesterone receptor functions as transcriptional activator or repressor. | A4GT83 | - | Q9DDU9 | - | P07812 |
| 334 | KKKKRK R | 280 | Q9NQB0 | Transcription factor 7-like 2 | Participates in the Wnt signaling pathway and modulates MYC expression by binding to its promoter in a sequence-specific manner. | Q9YHE8 | C3Z5P5 | A0A1L8HM07 | K7GJ95 | A0A1D5P979 |
| 335 | KRKAASQER EAKETERKRR K | 281 | O94850 | Dendrin | Promotes apoptosis of kidney glomerular podocytes. Podocytes are highly specialized cells essential to the ultrafiltration of blood, resulting in the extraction of urine and the retention of protein | E7FAR1 | - | - | - | - |

| | | | | | | | | | | |
|---|---|---|---|---|---|---|---|---|---|---|
| 336 | K RTSSPHKEES PKKTK | 282 | P51003 | Poly(A) polymerase alpha | Polymerase that creates the 3'-poly(A) tail of mRNA's. Also required for the endoribonucleolytic cleavage reaction at some polyadenylation sites. | Q5SNY2 | C3Z3M0 | A0A1L8F081 | - | A0A1D5P5T8 |
| 337 | KKKRR | 283 | Q8IZX4 | Transcription initiation factor TFIID subunit 1-like | May act as a functional substitute for TAF1/TAFII250 during male meiosis, when sex chromosomes are transcriptionally silenced. | Q1LYC2 | C3Y3A3 | A0A1L8F6Z1 | - | B5AHE5 |
| 338 | KRKR | 284 | Q9NZC7 | WW domain-containing oxidoreductase | Putative oxidoreductase. Acts as a tumor suppressor and plays a role in apoptosis. Required for normal bone development | Q803A8 | C3ZVC1 | Q63ZM3 | - | Q5F389 |
| 339 | KKKR K | 285 | Q9Y3Y4 | Pygopus homolog 1 | Involved in signal transduction through the Wnt pathway. | A8WG61 | C3Z441 | A0A1L8H0D5 | - | E1BWE1 |
| 340 | KP RKR | 286 | Q9H4Z3 | Phosphorylated CTD-interacting factor 1 | May play a role in transcription elongation or in coupling transcription to pre-mRNA processing through its association with the phosphorylated C-terminal domain (CTD) of RNAPII largest subunit | A0A0R4IKJ1 | C3YV22 | A0A1L8ETX1 | - | E1BR74 |
| 341 | RRVKEEVLKQ LPPKK | 287 | Q9H4L7 | SWI/SNF-related matrix-associated actin-dependent regulator of chromatin subfamily A containing DEAD/H box 1 | DNA helicase that possesses intrinsic ATP-dependent nucleosome-remodeling activity and is both required for DNA repair and heterochromatin organization. | E7F1C4 | C3ZNR8 | A0A1L8HMC3 | K7FTH0 | A0A1D5PHR6 |
| 342 | KKKKKKDR KKKKF | 288 | Q86X95 | Corepressor interacting with RBPJ 1 | May modulate splice site selection during alternative splicing of pre-mRNAs | Q504G8 | C3Z8J5 | A0A1L8EW84 | K7GHQ5 | E1BVI7 |
| 343 | RKKRKKNK | | | | | | | | | |
| 344 | NRLRR KR | 289 | Q9BTC0 | Death-inducer obliterator 1 | Putative transcription factor, weakly pro-apoptotic when overexpressed | A1A5T6 | C3YI02 | A0A1L8GA10 | K7G4A0 | A0A1D5NXL6 |
| 345 | RLRK KRR | | | | | | | | | |
| 346 | KS PEKKRRK | 290 | Q9BRQ0 | Pygopus homolog 2 | Involved in signal transduction through the Wnt pathway. | A8WG61 | C3Z441 | Q8JHB8 | - | E1BWE1 |
| 347 | KRRPGPG PGVPPKRAR | 291 | P28340 | DNA polymerase delta catalytic subunit | As the catalytic component of the trimeric (Pol-delta3 complex) and tetrameric DNA polymerase delta complexes (Pol-delta4 complex), plays a crucial role in high fidelity genome replication, including in lagging strand synthesis, and repair. | Q2KNE0 | C3Y5K6 | Q642R8 | - | F1NQT0 |
| 348 | KKKGR | 292 | Q9Y6F1 | Poly [ADP-ribose] polymerase 3 | Involved in the base excision repair (BER) pathway, by catalyzing the poly(ADP-ribosyl)ation of a limited number of acceptor proteins involved in chromatin architecture and in DNA metabolism. | Q7ZVB0 | C3Z2P3 | Q5M9A2 | - | E1BSI0 |
| 349 | KKKRRSR | 293 | P36402 | Transcription factor 7 | Transcriptional activator involved in T-cell lymphocyte differentiation. Necessary for the survival of CD4+ CD8+ immature thymocytes. Isoforms lacking the N-terminal CTNNB1 binding domain cannot fulfill this role. | A5A8K1 | - | - | - | - |
| 350 | KKKRKHR | 294 | Q92800 | Histone-lysine N-methyltransferase EZH1 | Polycomb group (PcG) protein. Catalytic subunit of the PRC2/EED-EZH1 complex, which methylates 'Lys-27' of histone H3, leading to transcriptional repression of the affected target gene. | Q08BS4 | C3YCV4 | Q4V863 | - | A0A1D5PN50 |
| 351 | KREGDDW SQLNVLKKRR | 295 | Q9BWG6 | Sodium channel modifier 1 | Plays a role in RNA splicing, possibly contributing to the recognition of non-consensus donor sites. | Q2YDS5 | - | A0A1L8FCL5 | K7GHG9 | - |
| 352 | KEDPPNV GKVKKAAKR | 296 | Q8IZA3 | Histone H1oo | May play a key role in the control of gene expression during oogenesis and early embryogenesis, presumably through the perturbation of chromatin structure. | Q4V956 | - | Q5XHJ2 | - | - |
| 353 | KKKKRKREK | 297 | Q9HCS4 | Transcription factor 7-like 1 | Participates in the Wnt signaling pathway. Binds to DNA and acts as a repressor in the absence of CTNNB1, and as an activator in its presence. | Q9YHE8 | C3Z5P5 | A0A1L8GRU8 | K7GJ95 | A0A1D5P979 |
| 354 | KRRKKRRH SKVNGLQRGR KK | 298 | P23497 | Nuclear autoantigen Sp-100 | Together with PML, this tumor suppressor is a major constituent of the PML bodies, a subnuclear organelle involved in a large number of physiological processes including cell growth, differentiation and apoptosis. | F1R6N2 | - | A0A1L8GBC5 | - | - |
| 355 | KKRW QQRGRKANTR PLKRRRKRGPR | | | | | | | | | |
| 356 | KKCS ETWKTIFAKE KGKF | | | | | | | | | |
| 357 | RRKRR | 299 | Q8NDT2 | Putative RNA-binding protein 15B | RNA-binding protein that acts as a key regulator of N6-methyladenosine (m6A) methylation of RNAs, thereby regulating different processes, such as alternative splicing of mRNAs and X chromosome inactivation mediated by Xist RNA | F1RCY7 | - | A0A1L8GHD9 | - | F1NRZ4 |

| | | | | | | | | | | |
|---|---|---|---|---|---|---|---|---|---|---|
| 358 | K QKKK | 300 | Q8IXJ9 | Putative Polycomb group protein ASXL1 | Probable Polycomb group (PcG) protein involved in transcriptional regulation mediated by ligand-bound nuclear hormone receptors, such as retinoic acid receptors (RARs) and peroxisome proliferator-activated receptor gamma (PPARG). | F1Q5H5 | C3YBN3 | A0A1L8ERK1 | - | A0A1D5PLQ0 |
