## Supplementary Table S2 for "Origin of the nuclear proteome on the basis of pre-existing nuclear localization signals in prokaryotic proteins"

**Supplementary Table S2.** Prokaryotic proteins with or without predicted NLSs.

| Protein | UniProt ID | Organism | Molecular weight, kDa | Molecular weight of EGFP fusion, kDa | Predicted NLSs | In-domain localization of predicted NLS | $F_{nuc}/F_{cyt}$, mean±s.d. |
|---|---|---|---|---|---|---|---|
| EGFP | - | - | 29.4 | - | no | no | 1.16±0.10 |
| Prokaryotic proteins with predicted NLSs | | | | | | | |
| KiaC (S.sp) | Q79PF4 | *Synechococcus sp.* | 58.0 | 87.4 | KYRARR | KaiC 1 domain (DNA binding) | 0.10±0.05 |
| | | | | | RRELFRLVAR | | |
| | | | | | RRRRT | | |
| | | | | | FKMRGSWH | KaiC 2 domain (DNA binding) | |
| Alr4331 | Q8YP69 | *Anabaena sp.* | 76.2 | 105.5 | RLKRT | S1 domain (interaction with pre-RNA) | 0.29±0.07 |
| | | | | | RRARTRRSR | no | |
| | | | | | RRRRRR | no | |
| KiaC (A.sp) | Q8YT40 | *Anabaena sp.* | 57.9 | 87.3 | RLQYAIRKYKAKR | KaiC 1 domain (DNA binding) | 0.33±0.08 |
| LigA (E.coli) | C3T152 | *Escherichia coli* | 74.0 | 103.4 | RTTLRHHEYLYH | DNA ligase | 0.37±0.09 |
| | | | | | RITAKRP | Nucleotide-binding pocket | |
| GyrB | D0XEB0 | *Vibrio harveyi* | 89.0 | 118.4 | RRGLSLQRYKGLGEMNPDQLWETTMDPETRR | DNA gyrase B subunit | 0.48±0.09 |
| LigA (S.sp) | Q5N2P4 | *Synechococcus sp.* | 74.3 | 103.7 | RSWDQRWRK | nucleotide binding pocket | 0.56±0.18 |
| PriA | Q31QI6 | *Synechococcus* | 89.2 | 118.6 | RRSQRRIRAR | no | 1.19±0.27 |

| | | | | | | | |
|---|---|---|---|---|---|---|---|
| | | *sp* | | | RIARRHRW | Primosomal DNA replication, repair, and recombination domains | |
| RecQ | P15043 | *Escherichia coli* | 70.0 | 99.4 | RIVALKPK | no | 1.26±0.2 |
| | | | | | RKLFAKLRKLRKS | The HRDC (interactions with DNA and protein) | |
| Lig | C6A2U9 | *Thermococcus sibiricus* | 63.4 | 92.8 | RRFRRKY | DNA-binding site | 1.35±0.25 |
| | | | | | RLKGGR | no | |
| PolB | P21189 | *Escherichia coli* | 90.0 | 119.4 | RLVYRKRLRRPLSEY | DNA polymerase type-II subfamily catalytic domain | 1.62±0.30 |
| SigA1 | P38023 | *Synechococcus sp.* | 45.7 | 75.1 | KAKAKVRKTY | no | 4.89±1.84 |
| | | | | | RRRLFRGRR | Sigma-70 factor domain-2, RNA synthesis | |
| | | | | | KKYMNR | no | |
| Dcm | P0AED9 | *Escherichia coli* | 53.0 | 82.4 | WKYLYRYAKKH | SAM-dependent MTase C5-type (DNA interaction, methylation) | 7.30±2.71 |
| Control prokaryotic proteins (wighout predicted NLSs) | | | | | | | |
| NblS | Q8RQ68 | *Synechococcus sp.* | 73.9 | 103.3 | no | no | 0.20±0.06 |
| Tuf | Q8YP63 | *Anabaena sp.* | 44.8 | 74.2 | no | no | 0.26±0.05 |
| ClpB | P63284 | *Escherichia coli* | 95.6 | 125.0 | no | no | 0.28±0.10 |
| GlpK | P0A6F3 | *Escherichia coli* | 56.2 | 85.6 | no | no | 0.31±0.08 |
| Pgi | Q8YY05 | *Anabaena sp.* | 57.8 | 87.2 | no | no | 0.31±0.07 |

| | | | | | | | |
|---|---|---|---|---|---|---|---|
| YgiK | P42592 | *Escherichia coli* | 88.3 | 117.7 | no | no | 0.42±0.09 |
| NifJ | Q06879 | *Anabaena sp.* | 132.2 | 161.6 | no | no | 0.44±0.12 |
| AccC | P24182 | *Escherichia coli* | 49.3 | 78.7 | no | no | 0.45±0.13 |
| TynA | P46883 | *Escherichia coli* | 84.4 | 113.8 | no | no | 0.48±0.08 |
| FusA | P0A6M8 | *Escherichia coli* | 77.6 | 107.0 | no | no | 0.50±0.13 |
| Ppc | Q31KY7 | *Synechococcus sp.* | 117.3 | 146.7 | no | no | 0.51±0.12 |
| AchB | Q31PT6 | *Synechococcus sp.* | 92.3 | 121.7 | no | no | 0.63±0.15 |
| DmsA | P18775 | *Escherichia coli* | 90.4 | 119.8 | no | no | 0.79±0.12 |
| PrfC | Q8YP23 | *Anabaena sp.* | 61.3 | 90.7 | no | no | 1.06±0.11 |
| CasA | Q46901 | *Escherichia coli* | 55.5 | 84.9 | no | no | 1.71±0.25 |
