## Supplementary Table S3 for "Origin of the nuclear proteome on the basis of pre-existing nuclear localization signals in prokaryotic proteins"

**Supplementary Table S3.** Predicted NLSs from prokaryotic proteins are able to target EGFP to the cell nucleus.

| Protein | Predicted NLSs | In-domain localization of predicted NLS | $F_{nuc}/F_{cyt}$, mean±s.d. |
|---|---|---|---|
| EGFP | - | - | 1.16±0.10 |
| LigA (S. sp.) | RSWDQRWRK | Nucleotide binding pocket | 1.29±0.11 |
| PriA | RRSQRRIRAR | no | 3.24±0.69 |
| | RIARRHRW | Primosomal DNA replication, repair, and recombination domains | 1.98±0.52 |
| RecQ | RIVALKPK | no | 1.31±0.14 |
| | RKLFAKLRKLRKS | The HRDC (interactions with DNA and proteins) | 4.01±0.49 |
| Lig | RRFRRKY | DNA-binding site | 2.32±0.29 |
| | RLKGGR | no | 1.34±0.49 |
| PolB | RLVYRKRLRRPLSEY | DNA polymerase type-II subfamily catalytic domain | 4.03±0.46 |
| SigA1 | KAKAKVRKTY | no | 1.74±0.30 |
| | RRRLFRGRR | Sigma-70 factor domain-2, RNA synthesis | 2.87±0.41 |
| | KKYMNR | no | 1.80±0.58 |
| Dcm | WKYLYRYAKKH | SAM-dependent MTase C5-type (DNA interaction, methylation) | 1.80±0.27 |
