## Supplementary Table S4 for "Origin of the nuclear proteome on the basis of pre-existing nuclear localization signals in prokaryotic proteins"

**Supplementary Table S4.** Detection of NLSs inside prokaryotic proteins by site-specific mutagenesis.

| **Protein** | **Protein $F_{nuc}/F_{cyt}$, mean±s.d.** | **Mutant** | **Mutant $F_{nuc}/F_{cyt}$, mean±s.d.** |
|---|---|---|---|
| PriA | 1.19±0.27 | mNLS1 | 1.05±0.15 |
| | | mNLS2 | 1.23±0.31 |
| | | mNLS1+mNLS2 | 1.14±0.18 |
| SigA1 | 4.89±1.84 | mNLS1 | 1.52±0.27 |
| | | mNLS2 | 0.96±0.16 |
| | | mNLS3 | 1.28±0.16 |
| | | mNLS1+ mNLS2 | 0.66±0.16 |
| | | mNLS1+ mNLS3 | 1.10±0.12 |
| | | mNLS2+ mNLS3 | 0.49±0.15 |
| | | mNLS1+ NLS2+mNLS3 | 0.90±0.21 |
| LigA (S.sp) | 0.56±0.18 | mNLS | 0.79±0.17 |
| PolB | 1.62±0.30 | mNLS | 1.07±0.14 |
| Dcm | 7.30±2.71 | mNLS | 0.76±0.12 |
| RecQ | 1.26±0.2 | mNLS1 | 1.32±0.12 |
| | | mNLS2 | 0.96±0.17 |
| | | mNLS1+mNLS2 | 1.01±0.08 |
| Lig | 1.35±0.25 | mNLS1 | 1.09±0.14 |
| | | mNLS2 | 1.10±0.12 |
| | | mNLS1+mNLS2 | 1.06±0.12 |
